# Identification of molecular pathways involved in the anti-liver cancer activity of madecassic acid

**DOI:** 10.1101/2025.08.17.670739

**Authors:** Geraud N. Sansom, Emily P. Friar, David Shorthouse, Thao Thi Phuong Tran, Fernando Sialana, Thao Thi Do, Chien Van Tran, Yen Thi Hoang Be, Sung Van Tran, Michelle D. Garrett, Christopher J. Serpell

## Abstract

Natural products are a time-tested source of medicinal compounds, and there is huge interest in discovery of new drugs from plant sources. However, in many cases the effects are not as well understood, strong, or selective, as would be hoped. An example of this is madecassic acid (MA), which has promising activity against liver cancer, activity which can be improved through chemical modifications, although a molecular understanding of its mechanism of action is unknown. In this report we have used chemical proteomics to identify the proteins with which madecassic interacts in liver cancer cells, and used RNAseq to support those findings, as well as showing more precisely how chemical modifications can focus and amplify MA activity. Our results show that madecassic acid interacts with a number of proteins relating to metabolism, nucleic acid processing, and protein folding, which have been previously identified as linked to liver cancer. This provides a route from phenotypic- to target based-drug discovery and develop new potential treatments for a globally challenging disease.

## Introduction

Liver cancer is a global health concern, with nearly 900,000 new cases and 800,000 deaths per year.^1^ In some locations the problem is particularly severe: in Viet Nam, for example, liver cancer kills more than 25,000 people each year; it is the country’s most prevalent (15%) and fatal (21%) cancer.^2–4^ The unusual severity in Viet Nam^5^ is due to globally leading hepatitis burden,^4,6,7^ vaccine hesitancy^8^, alcohol consumption^9–11^ and unregulated agrochemicals.^12–14^ It is also a leading cancer in nearby Laos, Thailand, Singapore, and Cambodia.^5,15^ Outcomes for liver cancer patients in Viet Nam are poor – the best-case median survival is 10 months^16^ due to 80% of diagnoses occurring late.^2,4^ Only 12% of patients receive treatments at cure, while 56% get merely symptomatic relief.^17^ Prohibitive personal healthcare costs healthcare are compacted in rural areas where provision is under resourced and long journeys are required.^16^ The associated costs put 37% of families of patients into poverty.^2,18,19^

However, Viet Nam also has resources, which could reduce liver cancer burden. Herbal products are widely used in cancer treatment^20–22^ and are a source of modern medicines.^23^ Madecassic acid (MA), a compound from *Centella asiatica*, a medicinal and nutritional herb popular in Viet Nam,^24^ can kill liver cancer cells, with its efficacy improved by chemical modification.^25^ Development of MA derivatives as a new drug could give better outcomes for liver cancer patients, and the drug could be produced sustainably and equitably inside Viet Nam. However, for this to occur, it is important to understand how MA exerts its effects and whether these could be fine-tuned through chemical modification to influence certain targets

We here report mechanistic investigation of the activity of madecassic acid in liver cancer cells by combining photoaffinity labelling with proteomics and RNAseq. Our results show that madecassic acid works on specific proteins and pathways, resulting in an anticancer effect which is consistent with hepatic specificity and shows potential for fine-tuning through chemical modification.

## Results

We adopted a photoaffinity approach^26–29^ to identifying the proteins with which MA interacts in liver cancer cells. This strategy was selected as it would allow us to identify interactions even if they have relatively weak binding constants, since a new covalent link would be formed with the photoaffinity probe. The photoaffinity process requires MA to be appended with a photoaffinity tag, in this case an aryl diazirine combined with an alkyne for subsequent attachment of a biotin which would allow pulldown using streptavidin coated beads. The photoaffinity tag must be installed on MA in a position which does not impair its biological activity, and thus bias the proteomic results. To identify such a location, we needed to performed structure-activity relationship experiments which would compare MA (**1**) along with the more potent 1,10-diaminodecane amidated version (**3**)^25^ and a series of new derivatives with blocking groups on the potential points for modification: the 2-OH, 3-OH or 23-OH locations, and terminal amine on the modified version (**Scheme 1**).

**Scheme 1.**
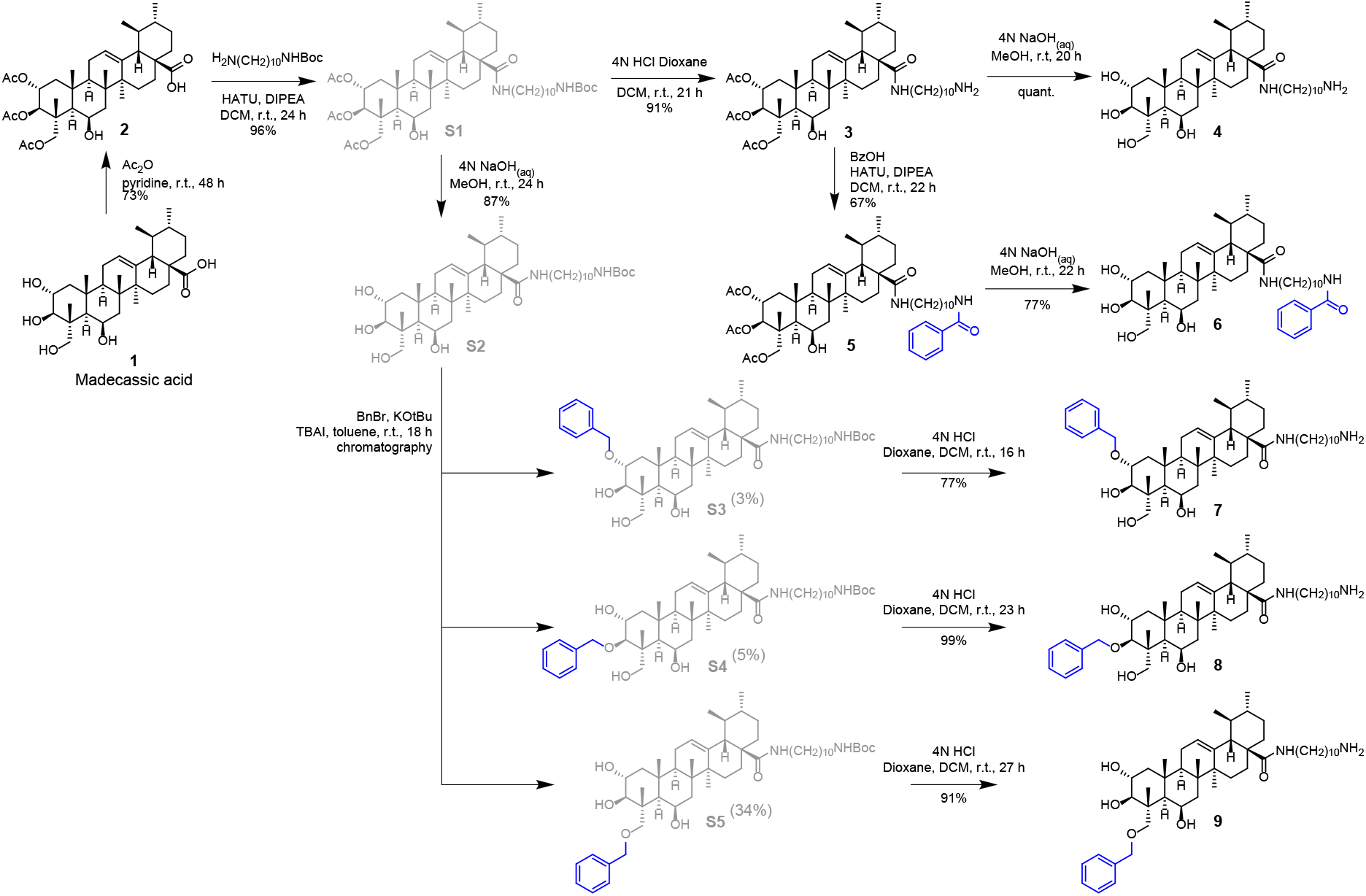
Synthesis of MA derivatives for structure-activity relationship study.

Synthesis of derivatives started from MA isolated from *C. asiatica* grown in Hue province, Viet Nam, extracted according to our optimised procewas produced by using benzene as a multifunctional core from whichdures.^30^ Selective functionalisation of 2-OH, 3-OH or 23-OH presented challenges and there was no literature precedent for selective functionalisation of any of the three accessible hydroxyls at will. A 1:1.06:0.21 mixture of 2,23-:3,23:2,3-diacetylated products, as well as some triacetylated material, could be obtained using 3.5 eq. of acetic anhydride, but separation of the mixture proved intractable. Expecting a bulky electrophile to react preferentially with the primary 23-OH, we reacted MA with TBSCl (*tert*-butyldimethylsilyl chloride), but obtained the 23-OTBS product in only 5% yield, which we regarded as impractical. Instead, we were able to couple the amide using *tert*-butyl 10-aminodecylcarbamate, HATU (hexafluorophosphate azabenzotriazole tetramethyl uronium), *N,N*-diisopropylethylamine, and dichloromethane (Scheme S1). Benzylation of the alcohol positions was achieved in a 1.:1.10:7.93 mixture of 2-:3-:23-benzylated products which were more amenable to chromatography separation. The amino position was benzoylated using benzoic acid, HATU, DIPEA and dichloromethane.

Using a 72 hour MTT assay MA itself (**1**) was found to have an IC_50_ of 109 µM against the HepG2 hepatocellular carcinoma (HCC) cell line, and the acetylated and 1,10-diaminodecane amidated derivative (**3**) gave a value of 1.56 µM, both in line with previously reported values (**Table 1)**.^25^ We found that each of the modifications (acetylation of OH-2, -3, and -23 to give **2**, or amidation with 1,10-diaminodecane to give **4**) resulted in an order-of-magnitude reduction in IC_50_, with their combination (i.e. **3**) giving an approximately hundred-fold reduction, indicating that these modifications were complementary and orthogonal in their effects. Addition of a benzoyl group to the end of the amide/amine chain resulted in loss of activity, whether the alcohols were acetylated (**5**) or not (**6**), thus ruling out that position as an option for attachment of the tag. Conversely, any of the three accessible alcohols (2-OH, 3-OH, 23-OH) could be benzylated, leading to minimal effects on the IC_50_ compared to the non-benzylated analogue **4** with the ranking of **8** > **7** > **9** in terms of similarity. Since the activities of **8** and **7** were close, but **7** was significantly easier to access from a synthetic point of view, we decided to attach the tag to the 3-OH.]

**Table 1.** Viability of HepG2 HCC cells after treatment with MA and derivatives, tested using the MTT assay.

| Compound | R<sub>1</sub> | R<sub>2</sub> | R<sub>3</sub> | R<sub>4</sub> | IC<sub>50</sub> (μM) | Std Dv. (n) |
| --- | --- | --- | --- | --- | --- | --- |
| **Ellipticine** (control) | - | - | - | - | 0.6702 | 0.1622 (6) |
| **1** (madecassic acid) | H | H | H | OH | 109.4 | 11.02 (3) |
| **2** | Ac | Ac | Ac | OH | 19.37 | 2.212 (3) |
| **3** | Ac | Ac | Ac | NH(CH<sub>2</sub>)<sub>10</sub>NH<sub>2</sub> | 1.562 | 0.3910 (3) |
| **4** | H | H | H | NH(CH<sub>2</sub>)<sub>10</sub>NH<sub>2</sub> | 11.33 | 1.116 (4) |
| **5** | Ac | Ac | Ac | NH(CH<sub>2</sub>)<sub>10</sub>NHBz | >250 | N.A. (4) |
| **6** | H | H | H | NH(CH<sub>2</sub>)<sub>10</sub>NHBz | >250 | N.A. (4) |
| **7** (selected) | Bn | H | H | NH(CH<sub>2</sub>)<sub>10</sub>NH<sub>2</sub> | 6.503 | 0.8404 (5) |
| **8** | H | Bn | H | NH(CH<sub>2</sub>)<sub>10</sub>NH<sub>2</sub> | 8.511 | 1.082 (5) |
| **9** | H | H | Bn | NH(CH<sub>2</sub>)<sub>10</sub>NH<sub>2</sub> | 2.134 | 0.02625 (5) |

**Scheme 2.**
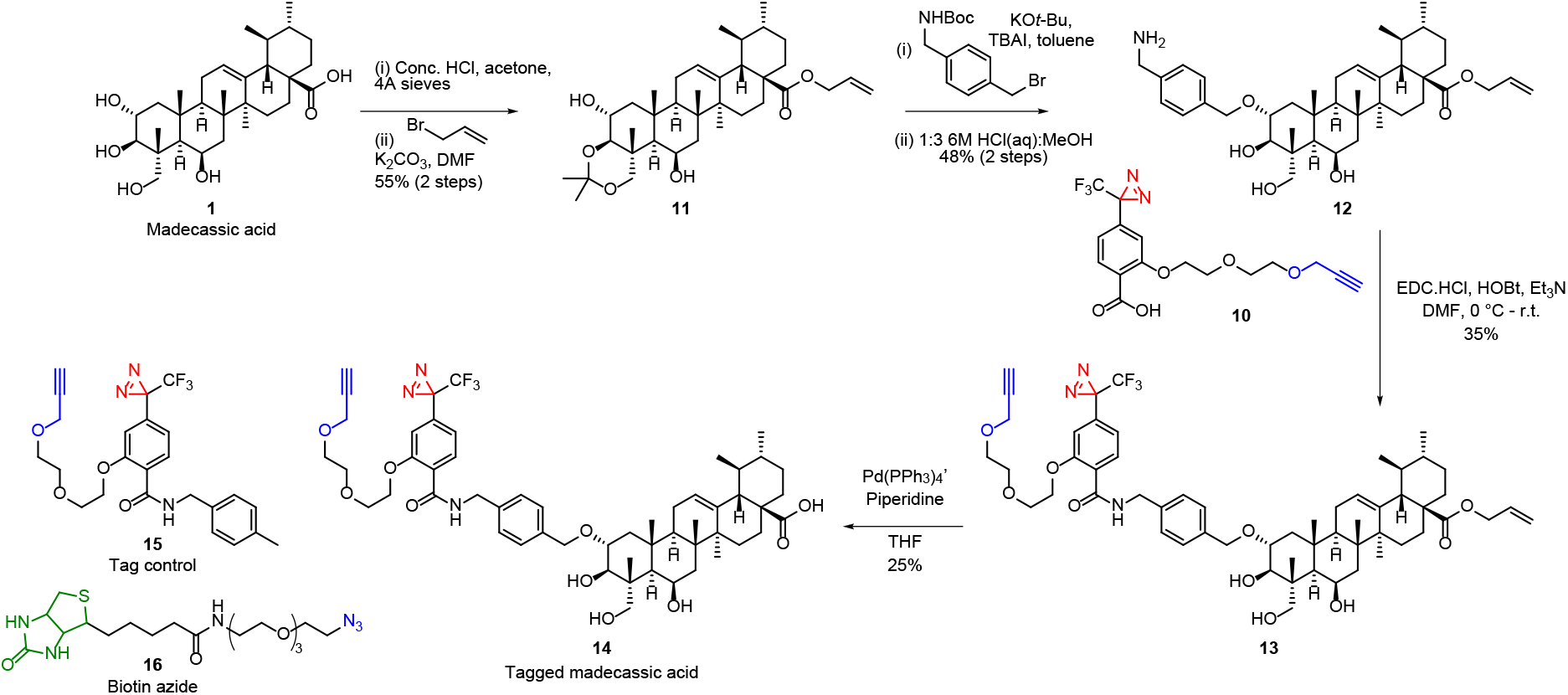
Synthesis of tagged MA.

The diazirine/alkyne tag **10** was produced by using benzene as a multifunctional core from which diazirine, alkyne, and bioconjugation unit (carboxylic acid) extend (**Scheme S2/S3**). Synthesis of the tagged madecassic acid (**Scheme 2**) was achieved by acetonide protection of 3- and 23-OH following procedures previously used for the related asiatic acid,^31^ followed by allyl protection of the carboxylic acid to give **11**. The remaining alcohol was converted to an ether by reaction with a Boc-protected benzyl amine featuring a para-bromomethyl unit. Removal of the Boc group (with concurrent alcohol deprotection) gave **12**, allowing attachment of the photoaffinity tag **10** through an amide (**13**). The carboxylic acid was recovered after allyl deprotection, giving the final tagged MA (**14**).

**Scheme 3.**
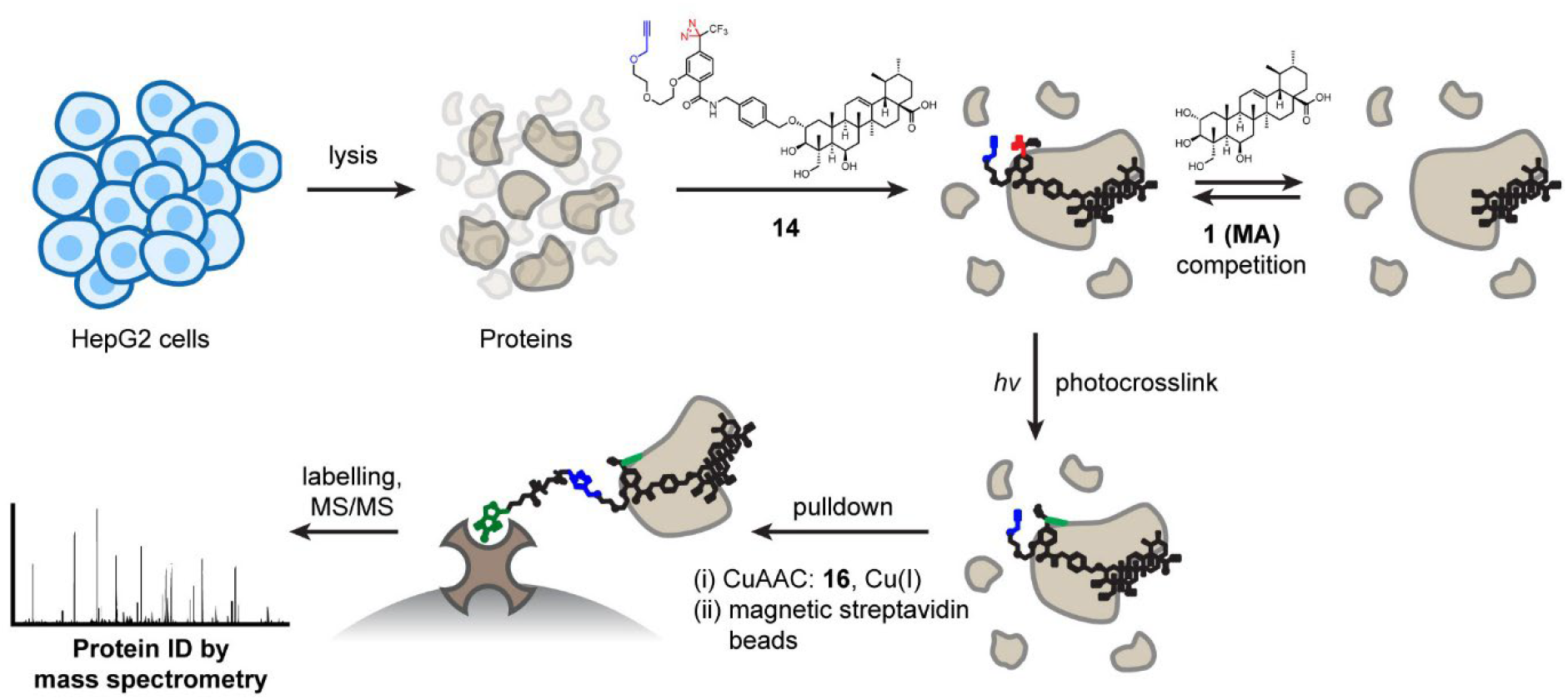
Workflow for photoaffinity pulldown-enabled proteomics.

To optimise conditions for the crosslinking reaction, the irradiation time required to activate the photoaffinity probe was tested. The tag control compound **15** (without MA) was irradiated at 365 nm, and the loss of optical absorbance at the same wavelength was measured at timepoints up to 256 seconds. It was found that, under these conditions, the half-life of the tag was 24.9 seconds (see **Figure S1**). Pulldown experiments were then conducted using the MA-photoaffinity compound **14** as shown in **Scheme 3**. HepG2 cells were lysed, and any proteins which displayed endogenous biotin were removed by passing through streptavidin columns. The protein content was then determined by Bradford assay, and concentrations were normalised to 3 mg/mL protein. Compound **14** was then added to the cell lysate and incubated at 4 °C to allow association with protein targets. UV irradiation was then applied for five minutes, to form covalent bonds with target proteins. The attached alkyne was then reacted with biotin-azide **16** to give a triazole linkage via copper catalysed azide-alkyne cycloaddition (CuAAC) using CuSO_4_, sodium ascorbate, and the ligand THPTA (tris-hydroxypropyltriazolylmethylamine) to generate Cu(I) *in situ*. After workup to remove reagents and side products, the proteins were isolated by centrifugation. Pulldown was achieved by dissolving the pellets in HEPES buffer and collecting proteins which have been labelled with biotin *via* the photoaffinity probe using magnetic streptavidin coated beads. The captured proteins were digested on bead and the released peptides labelled with TMTpro, multiple samples are combined for LC-MS/MS analysis using SPS-MS^3^ with real-time search.^32^ This enabled multiplexed quantification of enriched proteins across three conditions (all performed as three biological repeats): a DMSO control (no compound added); the tagged-MA compound **14**; and then **14** in the presence of excess (18.5 µM) **MA** as a competitor. The proteins which increase in abundance in the tagged-MA sample compared to the DMSO control sample represent candidates from photolabeling reactions, rather than non-specific adherence to the streptavidin beads. The proteins which are due to specific interactions with the MA moiety (rather than interactions with the tag, or new non-specific interactions due to modification) should then decrease in abundance in the competition sample compared with the no competition sample, due to the excess of MA.

Proteomics data was transformed to log2+0.1, normalised, and then differential expression analysis was performed with each normalised data set using limma open source software for **14** (tagged MA) versus DMSO and **14** versus [**14** plus MA competition] (**Fig. 1a**). After comparison of normalisation techniques, 67 proteins were identified (**Fig. 1b**), which were both substantially (log2 fold change ≥ 1.5) and significantly (P < 0.05) increased in abundance in the Tagged sample compared to the DMSO sample, and decreased in abundance in the Competition sample versus the Tagged sample with examples highlighted in the volcano plots in **Figure 1a**. Metascape analysis of these 67 proteins (**Fig. 1c**) revealed that the top specific cell processes associated with the hits included metabolic reprogramming in colon cancer, protein folding and aerobic glycolysis (−log(P) > 10). Interestingly, the entire second half of the glycolytic cascade is represented within the list of 67 proteins (ALDOA, ENO1, LDHA (the bacterial analogue is a known MA binder^33^), PGAM1, PGK1, PKM), which is important from an oncology perspective, as the shift of glucose metabolism away from oxidative phosphorylation to glycolysis, even in aerobic conditions (the Warburg effect) is a characteristic of cancer. Indeed, there is building evidence for its role in liver cancer,^34^ meaning that treatment with madecassic acid (or derivatives) could selectively target liver cancer cells. Among these proteins, ENO1 (Enolase 1) stood out as one of the most enriched (Log2(fold change) = 1.8 – 3.0 depending on normalisation method), which is an important result because the protein is overexpressed in many cancers,^35^ including hepatocellular carcinoma where it is a prognostic marker.^36–38^ It has also been associated with liver cancer cell proliferation via CRISPR screens.^39,40^ With respect to protein folding, four out of eight members of the chaperone complex (CCT4, CCT6A, CCT7, and TCP1) were also present in the list. Overexpression of these proteins has been found to be a biomarker for a poorer prognosis in hepatocellular carcinoma,^41,42^ suggesting them as potential drug targets.^43,44^ Six heat shock proteins (HSP90B1, HSPA4, HSPA5, HSPA8, HSPA9, HSPD1) were found, all of which are upregulated in HCC.^45^ It is worth noting that many of the metabolic and protein-folding proteins mentioned above are high-abundance,^46^ and although the use of the competition experiment should limit non-specific interactions, these results need validating. Other pathways included the cellular response to interleukin 7 and interleukin 12 signalling. PDIA3 was also identified, which along with HSPD1 and ENO1 is part of the cellular response to interleukin-7 pathway, while interleukin-7 supports CD8+ T-cell activity in HCC.^47^ Immune response to cancer cells in HCC is also boosted by interleukin-12,^48^ signalling of which is indicated in identification of HSPA9, TCP1, and RPLP0. While these interleukin pathways may be useful for cancer treatment, their reliance on immune cells means that they cannot be the cause of the efficacy of our compounds in HepG2 culture. Finally, DNA-PK-Ku was identified in the Metascape list, which correlates with the identification of a series of DNA repair and nucleotide processing proteins in the list of 67 proteins, many of which have been linked to cell survival/proliferation (ATIC,^49^ HNRNPM,^50^ MTHFD1L,^51^ XRCC5^52^), poor prognosis (MTHFD1,^53^ PAICS^54^), or as potential therapeutic targets (HNRNPM,^55,56^ PRKDC^57^) in HCC. The proteins identified were regulated by HYF1A, MYC, and JUN (Fig. 1d), and consistent with proteins specifically associated with the liver and the HepG2 cell line specifically (Fig. 1e), providing an internal validation to the experiment. Given the wide nature of proteins identified in this pulldown experiment, and their relationship to processes related to HCC, it is possible that MA interacts weakly with a variety of targets, with the overall profile of changes resulting in the liver cancer cell killing activity. This would make it challenging to improve the therapeutic potential of MA in a specific manner, so comparison with a chemically modified analogue was needed.

**Figure 1.**
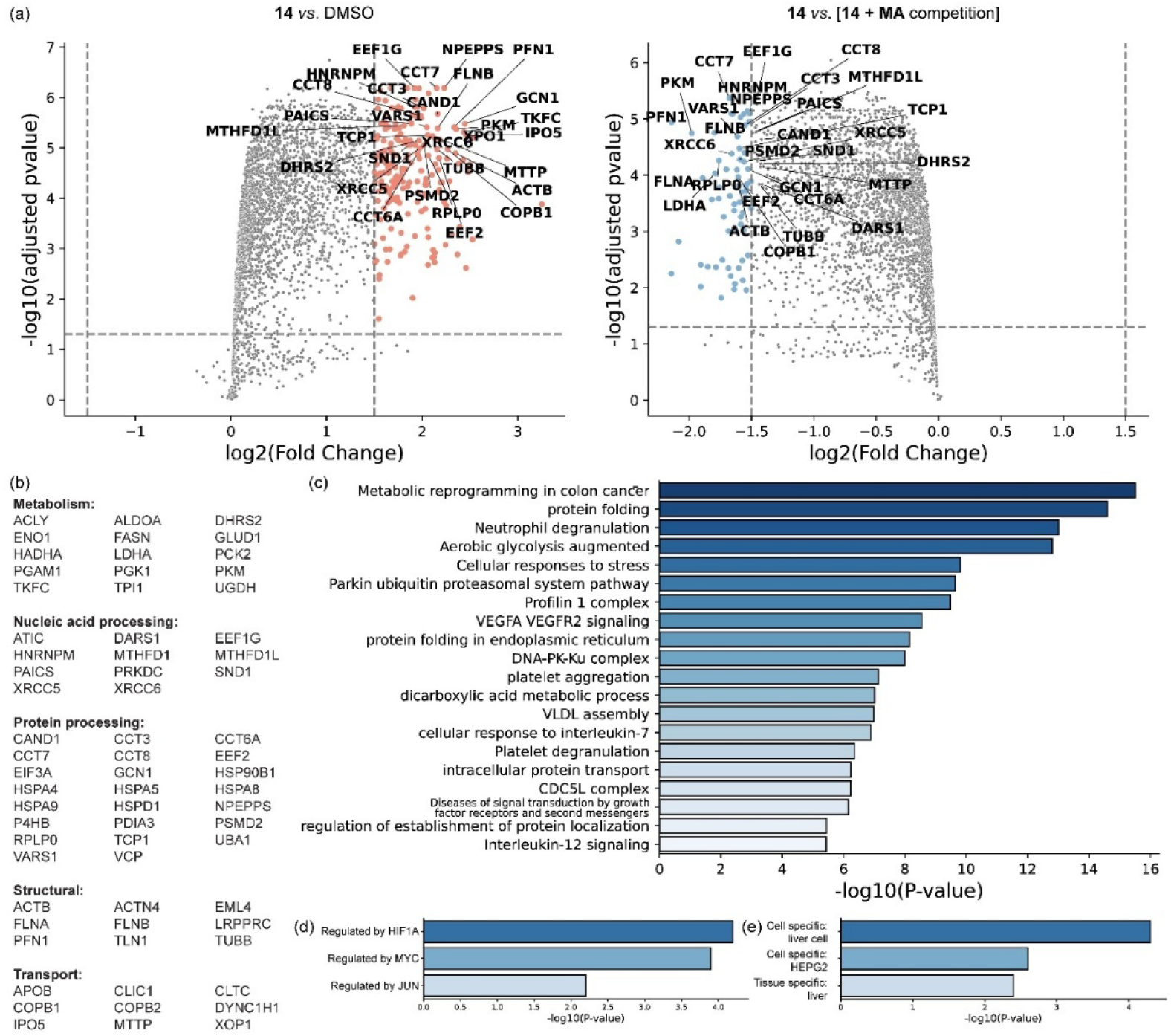
Proteomic analysis. (a) Volcano plots showing selection for proteins which are both more represented in the Tagged sample (**14**) vs DMSO control (left hand side), and reduced in the Tagged sample (**14**) when **MA** is added as a competitor (right hand side). (b) 67 hit proteins. (c) Enriched pathways identified by Metascape analysis (d) Predicted transcriptional regulators of identified proteins. (e) Cell type associations inferred from protein expression.

To both assess the cellular response to **MA**, and verify whether the greater cytotoxicity of the modified MA analogues such as **3** or **7** was due to similar mechanisms, RNAseq experiments were performed. Cells were treated with either MA or **7** for 6 and 24 hours at 1 × and 3 × GI_50_, with gene expression compared to a control experiment using only the same quantity of DMSO (**Fig. 2a, b**). The differences between the expression of the various genes were much more pronounced (in terms of hierarchical clustering and principal component analysis) at the later timepoint and higher dose. Across all samples, more genes were upregulated relative to the control than were downregulated. There was an excellent overlap of the genes influenced by both **MA** and **7**, with **7** appearing to affect a subset of those genes altered in expression by **MA** (**Fig. 2c**). The same overlap is seen if pathways are analysed instead of genes. This is consistent with the modifications applied to **MA** to give **7** limiting its interactions, resulting in a cleaner effect profile. Gene set enrichment analysis was used to look at gene ontology biological process pathway overlaps for both **MA** and **7** with 3 × GI_50_ for each compound at the 24 hour time point where the effect is strongest (Fig. **2d**). In line with the proteomics results, pathways relating to metabolism of sugars, and nucleic acid processing both rank highly. Consistent with the structure of madecassic acid itself, cholesterol biosynthesis was also a prominently influenced pathway. In more detail, (ESI) metabolic processing and biosynthesis of sterols, especially the estradiol response, dominated the upregulated pathways, while nucleic acid processing (ribosome biogenesis, tRNA, DNA, gene expression) dominated the downregulated pathways. MA bears structural similarities to estradiol, so this response may be the cell processing the compound. At the 3 × GI_50_ concentration, interleukin (IL-6 particularly), TNFα and JUN (and its promotor ATF) signalling were also all implicated. The differences were weaker at the 6 hour timepoint, but the overlap between **MA** and **7** remained, as did the estradiol signature, although it was weaker indicating that it is an adaptive response. In addition, there was a strong increase in UV damage, ER stress, and apoptotic pathways.

**Figure 2.**
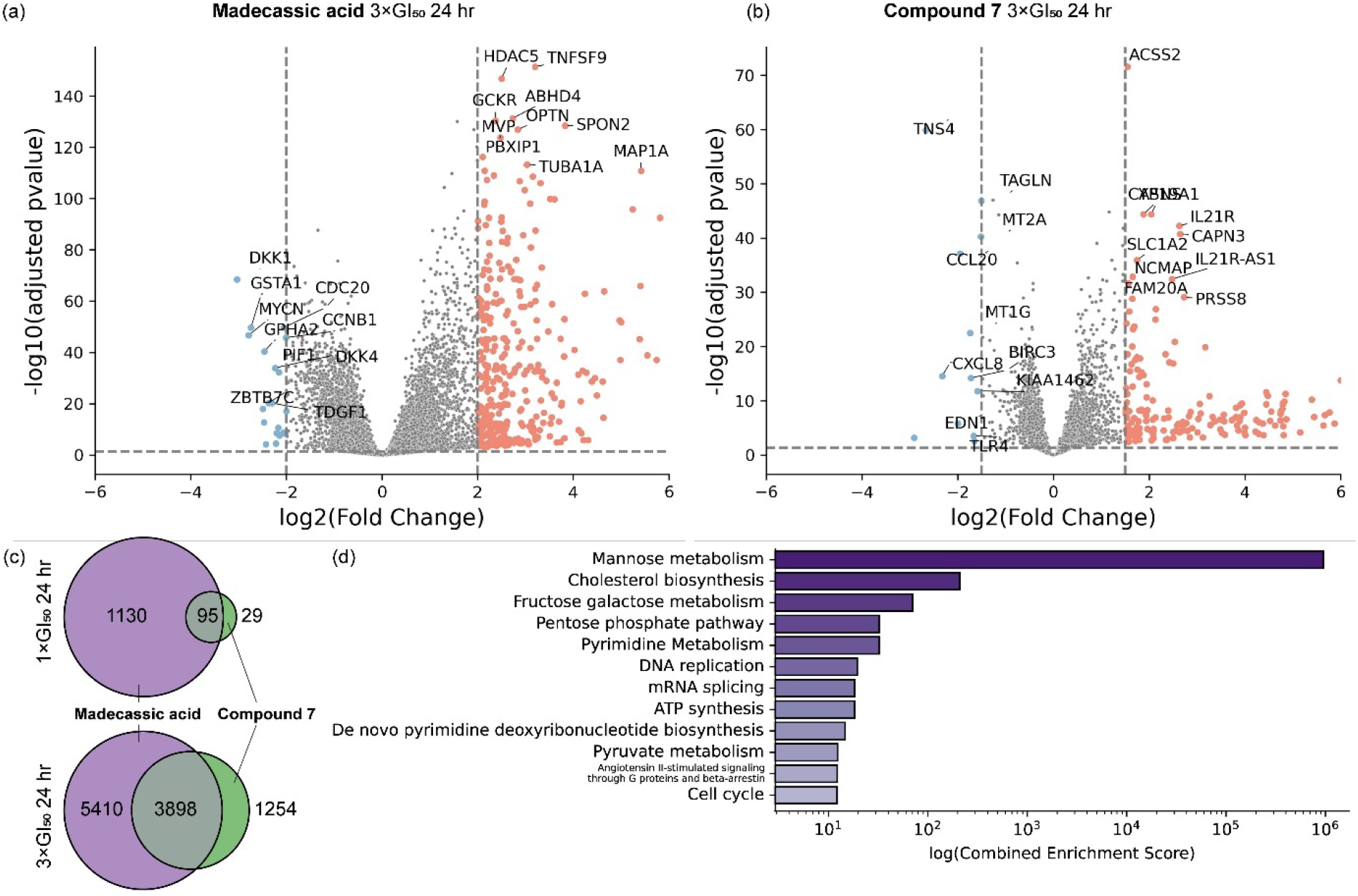
RNAseq data. Volcano plots for RNA expression in HepG2 cells treated with (a) **MA** and (b) **7** for 24 hours at 3 x the GI_50_. (c) Venn diagrams showing the overall between changes in RNA expression for **MA** and **7** at 1 and 3 x the GI_50_ after 24 hr. (d) Pathway analysis of genes regulated under influence of both **MA** and **7** at 3 x the GI_50_ after 24 hr.

Proteomics and RNAseq are two quite distinct ways of looking at how a compound can influence cellular behaviour, and we expect to see different results and since binding of a certain protein to a compound such as **MA** or **7** is not necessarily related to whether expression of that protein is upregulated. Moreover, the proteomics was performed using lysate of cells which had not previously been exposed to **MA** or **7**, while the RNAseq gives us a snapshot of the transcriptional response of the cell some hours of exposure. Nonetheless, the similarities in terms of metabolism and nucleic acid processing are remarkable. There is also a further important link between the two datasets in that the proteomics hits are regulated by the same transcription factors found in the RNAseq: HIF1A (−log_10_P >4), MYC (−log_10_P >3.5), and JUN (−log_10_P >2)(**Fig. 1d**).

Interestingly, we did not find any links to proteins previously reported to interact with madecassic acid or similar compounds. **MA** itself has been shown to inhibit tyrosine-protein phosphatase non-receptor type 1,^58^ and to act as an agonist of PPARγ.^59^ Although many similar triterpenes inhibit glycogen phosphorylase, the C6-OH of **MA** removes potency.^60^ All of these are primarily under investigation in relation to diabetes, obesity, and colitis rather than liver cancer, and the methodology used in those papers was focused on predetermined (candidate) proteins in certain conditions, while our unbiased approach looked at all proteins or RNAs within the HepG2 liver cancer cell line. **MA** also interacts with cytochrome P450 enzymes,^61^ although this is unremarkable for a drug compound. Related compounds have been investigated against liver-prevalent enzymes related to diabetes: carboxylesterase 1 and pancreatic lipase (which did not show up in our experiments) have been reported as targets of oleanolic acid and ursolic acid,^62^ however this was again a targeted study.

## Discussion

Nature continues to be a rich source of starting points for the discovery of medicines. Plants, animals, fungi, and bacteria produce a raft of compounds which all have some kind of bioactivity, and many of these can be found which interact with human biology for a variety of effects. Alongside notable successes in drug discovery using compounds which originate in natural sources, such as aspirin, paclitaxel, and artemisinin, there are an enormous number of publications reporting preliminary activity for natural products against all kinds of diseases. Sometimes, the same compound appears to work against all the most pressing medical needs of the day. In some cases, and notoriously for curcumin,^63^this is a result of it belonging to a compound class known as ‘pan-assay interfering compounds’ (PAINS),^64,65^ which have properties such as redox cycling, membrane disruption, aggregation, or covalent reactivity. For many other examples, there are no obvious PAINS properties, but nonetheless research has not progressed beyond initial testing. ^66^

It is often the case that these studies are conducted in countries which are rich in biodiversity, and hence the raw materials for natural-products-based drug discovery, and the diseases selected are targeted towards the specific healthcare needs of those nations. At the same time, there may not be an established history of drug discovery from these same countries. There is a need to create links which will enable drug discovery to happen globally in a more equitable sense, and the tools of chemical biology which enable us to understand how molecules influence biological systems have a role to play here in progressing natural product bioactivity beyond initial reports or development of nutritional products or dietary recommendations for which impact is hard to measure due to variability of composition as well as challenges of monitoring.

We have used photoaffinity labelling and mass spectrometric proteomics, combined with RNAseq, to identify the protein partners and down stream cellular effects of madecassic acid and its derivatives in liver cancer cells. To achieve this, we first identified a position on the molecule which did not impede its valuable phenotypic effect (killing of liver cancer cells). This structure-activity relationship enabled us to attach a photoaffinity tag and identify which proteins **MA** interacts with. Proteins relating to metabolic activity in cancer, protein folding, interleukin signalling, and DNA repair were identified, which have pre-established strong links to liver cancer progression and prognosis. RNAseq was then used to observe the cellular response to **MA** treatment, and whether the more potent derivative, **7** is working through the same processes. There was considerable overlap in response for both compounds, with **7** influencing a subset of those genes activated by **MA**, and links between the two datasets through interleukin signalling and transcription factors.

These results show that the bioactivity of the natural product has an intermediate specificity – it exerts its influence through specific pathways, and is not a PAINS compound – but in its natural form it is less specific than final compounds arising from target-based drug discovery. The fact that the modified version **7**, alters transcription of a subset of the genes influenced by the parental compound shows that fine-tuning of bioactivity is possible through chemical modifications. The next steps in testing the potential of madecassic acid derivatives as drugs for liver cancer is to perform structure-based optimisations for binding on each of the pathways identified, alongside cellular testing. In this way, the successes of phenotypic drug discovery can be combined with the information-rich precision of target-based drug discovery.^67,68^

## Conclusion

Through combination of chemical biology methods with bioinformatics, we have been able to identify the pathways through which madecassic acid and its derivatives exert their anti-liver cancer effect. These pathways relate to processes in metabolism, processing of nucleic acids and proteins, and transport, as well as structural proteins. Our investigations have shown that the selection of pathways influenced can be narrowed down, providing new leads for development of MA derivatives as sustainably and equitably developed and produced drugs. The development of MA derivatives against liver cancer is particularly relevant for Viet Nam and nearby countries where there is both the medical need, and the capacity to produce MA at scale through agricultural techniques.^30^ Further elucidation and advancement could lead to significant gains in wellbeing in the region, in line with the UN’s Sustainable Development Goals, while also having an impact on liver cancer patients worldwide.

## Supporting information

All supplementary data

## Acknowledgements

We thank the UK’s Engineering and Physical Science Research Council and Global Challenges Research Fund (EP/T020164/1) and the University of Kent, United Kingdom and Vietnam Academy of Science and Technology (VAST) for financial support of this project.

## Notes

### Competing Interest Statement

The authors have declared no competing interest.

## References

(1) Bray, F.; Laversanne, M.; Sung, H.; Ferlay, J.; Siegel, R. L.; Soerjomataram, I.; Jemal, A. Global Cancer Statistics 2022: GLOBOCAN Estimates of Incidence and Mortality Worldwide for 36 Cancers in 185 Countries. CA: A Cancer Journal for Clinicians 2024, 74 (3), 229–263. 10.3322/caac.21834.

(2) Pham, T.; Bui, L.; Kim, G.; Hoang, D.; Tran, T.; Hoang, M. Cancers in Vietnam-Burden and Control Efforts: A Narrative Scoping Review. Cancer Control 2019, 26 (1), 1073274819863802. 10.1177/1073274819863802.

(3) International Agency for Research on Cancer. Viet Nam Fact Sheet. https://gco.iarc.fr/today/data/factsheets/populations/704-viet-nam-fact-sheets.pdf.

(4) Nguyen-Dinh, S.-H.; Do, A.; Pham, T. N. D.; Dao, D. Y.; Nguy, T. N.; Jr, M. S. C. High Burden of Hepatocellular Carcinoma and Viral Hepatitis in Southern and Central Vietnam: Experience of a Large Tertiary Referral Center, 2010 to 2016. World Journal of Hepatology 2018, 10 (1), 116–123. 10.4254/wjh.v10.i1.116.

(5) Sung, H.; Ferlay, J.; Seigel, R. L.; Laversanne, M.; Soerjomataram, I.; Jemal, A.; Bray. Global Cancer Statistics 2020: GLOBOCAN Estimates of Incidence and Mortality Worldwide for 36 Cancers in 185 Countries. CA: a cancer journal for clinicians 2021, 71 (3), 209–249. 10.3322/caac.21660.

(6) Schweitzer, A.; Horn, J.; Mikolajczyk, R. T.; Krause, G.; Ott, J. J. Estimations of Worldwide Prevalence of Chronic Hepatitis B Virus Infection: A Systematic Review of Data Published between 1965 and 2013. The Lancet 2015, 386 (10003), 1546–1555. 10.1016/S0140-6736(15)61412-X.

(7) Le, V. Q.; Nguyen, V. H.; Nguyen, V. H.; Nguyen, T. L.; Sudenga, S. L.; Trinh, L. H.; Nguyen, V. T.; Nguyen, T. T. H. Epidemiological Characteristics of Advanced Hepatocellular Carcinoma in the Northern Region of Vietnam. Cancer Control 2019, 26 (1), 1073274819862793. 10.1177/1073274819862793.

(8) Thanh Thi Le, X.; Ishizumi, A.; Thi Thu Nguyen, H.; Thi Duong, H.; Thi Thanh Dang, H.; Manh Do, C.; Thi Pham, Q.; Thi Le, H.; Iijima, M.; Tohme, R. A.; Patel, P.; Abad, N. Social and Behavioral Determinants of Attitudes towards and Practices of Hepatitis B Vaccine Birth Dose in Vietnam. Vaccine 2020, 38 (52), 8343–8350. 10.1016/j.vaccine.2020.11.009.

(9) Ha, L.; Tran, A.; Bui, L.; Giovannucci, E.; Mucci, L.; Song, M.; Le, P. D.; Hoang, M.; Tran, H.; Kim, G.; Pham, T. Proportion and Number of Cancer Cases and Deaths Attributable to Behavioral Risk Factors in Vietnam. International Journal of Cancer 2023, 153 (3), 524–538. 10.1002/ijc.34549.

(10) Gish, R. G.; Bui, T. D.; Nguyen, C. T. K.; Nguyen, D. T.; Tran, H. V.; Tran, D. M. T.; Trinh, H. N.; International Group for Liver Health in Viet Nam. Liver Disease in Viet Nam: Screening, Surveillance, Management and Education: A 5-Year Plan and Call to Action. Journal of Gastroenterology and Hepatology 2012, 27 (2), 238–247. 10.1111/j.1440-1746.2011.06974.x.

(11) Huu Bich, T.; Thi Quynh Nga, P.; Ngoc Quang, L.; Van Minh, H.; Ng, N.; Juvekar, S.; Razzaque, A.; Ashraf, A.; Masud Ahmed, S.; Soonthornthada, K.; Kanungsukkasem, U. Patterns of Alcohol Consumption in Diverse Rural Populations in the Asian Region. Global Health Action 2009, 2 (1), 2017. 10.3402/gha.v2i0.2017.

(12) Ngaon, L. T.; Yoshimura, T. Liver Cancer in Viet Nam: Risk Estimates of Viral Infections and Dioxin Exposure in the South and North Populations. Asian Pacific journal of cancer prevention : APJCP 2001, 2 (3), 199–202.

(13) Krishnamurthy, P.; Hazratjee, N.; Opris, D.; Agrawal, S.; Markert, R. Is Exposure to Agent Orange a Risk Factor for Hepatocellular Cancer?—A Single-Center Retrospective Study in the U.S. Veteran Population. Journal of Gastrointestinal Oncology 2016, 7 (3), 426–432. 10.21037/jgo.2016.01.09.

(14) Cordier, S.; Thuy, L. T. B.; Verger, P.; Bard, D.; Dai, L. C.; Larouze, B.; Dazza, M. C.; Quinh, H. T.; Abenhaim, L. Viral Infections and Chemical Exposures as Risk Factors for Hepatocellular Carcinoma in Vietnam. International Journal of Cancer 1993, 55 (2), 196–201. 10.1002/ijc.2910550205.

(15) Liu, Z.; Jiang, Y.; Yuan, H.; Fang, Q.; Cai, N.; Suo, C.; Jin, L.; Zhang, T.; Chen, X. The Trends in Incidence of Primary Liver Cancer Caused by Specific Etiologies: Results from the Global Burden of Disease Study 2016 and Implications for Liver Cancer Prevention. Journal of Hepatology 2019, 70 (4), 674–683. 10.1016/j.jhep.2018.12.001.

(16) Le, D. C.; Nguyen, T. M.; Nguyen, D. H.; Nguyen, D. T.; Nguyen, L. T. M. Survival Outcome and Prognostic Factors Among Patients With Hepatocellular Carcinoma: A Hospital-Based Study. Clin Med Insights Oncol 2023, 17, 11795549231178171. 10.1177/11795549231178171.

(17) Phan, T.; Nguyen, T.; Dinh, H. Treatment Cost of Hepatocellular Carcinoma from The Health Insurance’s Perspective in Viet Nam. Value in Health 2017, 20 (9), A429. 10.1016/j.jval.2017.08.180.

(18) Ho, H. T.; Jenkins, C.; Nghiem, H. L. P.; Hoang, M. V.; Santin, O. Understanding Context: A Qualitative Analysis of the Roles of Family Caregivers of People Living with Cancer in Vietnam and the Implications for Service Development in Low-Income Settings. Psycho-Oncology 2021, 30 (10), 1782–1788. 10.1002/pon.5746.

(19) Jenkins, C.; Ho, H. T.; Nghiem, H. P. L.; Prue, G.; Lohfeld, L.; Donnelly, M.; Hoang, M. V.; Santin, O. A Qualitative Study on the Needs of Cancer Caregivers in Vietnam. Global Health Action 2021, 14 (1), 1961403. 10.1080/16549716.2021.1961403.

(20) Li, Y.; Martin, R. C. G. Herbal Medicine and Hepatocellular Carcinoma: Applications and Challenges. Evidence-Based Complementary and Alternative Medicine 2011, 2011, eneq044. 10.1093/ecam/neq044.

(21) Nguyen, H. M.; Nguyen, N. Y. T.; Chau, N. T. N.; Nguyen, A. B. T.; Tran, V. K. T.; Hoang, V.; Le, T. M.; Wang, H.-C.; Yen, C.-H. Bioassay-Guided Discovery of Potential Partial Extracts with Cytotoxic Effects on Liver Cancer Cells from Vietnamese Medicinal Herbs. Processes 2021, 9 (11), 1956. 10.3390/pr9111956.

(22) Nguyen, S. T.; Vo, P. H.; Nguyen, T. D.; Do, N. M.; Le, B. H.; Dinh, D. T.; Truong, K. D.; Pham, P. V. Ethanol Extract of Ginger Zingiber Officinale Roscoe by Soxhlet Method Induces Apoptosis in Human Hepatocellular Carcinoma Cell Line. Biomedical Research and Therapy 2019, 6 (11), 3433–3442. 10.15419/bmrat.v6i11.572.

(23) Newman, D. J.; Cragg, G. M. Natural Products as Sources of New Drugs over the Nearly Four Decades from 01/1981 to 09/2019. J. Nat. Prod. 2020, 83 (3), 770–803. 10.1021/acs.jnatprod.9b01285.

(24) Babu, T. D.; Kuttan, G.; Padikkala, J. Cytotoxic and Anti-Tumour Properties of Certain Taxa of Umbelliferae with Special Reference to Centella Asiatica (L.) Urban. Journal of Ethnopharmacology 1995, 48 (1), 53–57. 10.1016/0378-8741(95)01284-K.

(25) Van Loc, T.; Nhu, V. T. Q.; Van Chien, T.; Ha, L. T. T.; Thao, T. T. P.; Van Sung, T. Synthesis of Madecassic Acid Derivatives and Their Cytotoxic Activity. Zeitschrift für Naturforschung B 2018, 73 (2), 91–98. 10.1515/znb-2017-0172.

(26) Kita, M.; Hirayama, Y.; Yamagishi, K.; Yoneda, K.; Fujisawa, R.; Kigoshi, H. Interactions of the Antitumor Macrolide Aplyronine A with Actin and Actin-Related Proteins Established by Its Versatile Photoaffinity Derivatives. J. Am. Chem. Soc. 2012, 134 (50), 20314–20317. 10.1021/ja310495p.

(27) Smith, E.; Collins, I. Photoaffinity Labeling in Target- and Binding-Site Identification. Future Medicinal Chemistry 2015, 7 (2), 159–183. 10.4155/fmc.14.152.

(28) Zeng, Q.; Deng, H.; Li, Y.; Fan, T.; Liu, Y.; Tang, S.; Wei, W.; Liu, X.; Guo, X.; Jiang, J.; Wang, Y.; Song, D. Berberine Directly Targets the NEK7 Protein to Block the NEK7–NLRP3 Interaction and Exert Anti-Inflammatory Activity. J. Med. Chem. 2021, 64 (1), 768–781. 10.1021/acs.jmedchem.0c01743.

(29) Cheng, Y.-S.; Zhang, T.; Ma, X.; Pratuangtham, S.; Zhang, G. C.; Ondrus, A. A.; Mafi, A.; Lomenick, B.; Jones, J. J.; Ondrus, A. E. A Proteome-Wide Map of 20(S)-Hydroxycholesterol Interactors in Cell Membranes. Nat Chem Biol 2021, 17 (12), 1271–1280. 10.1038/s41589-021-00907-2.

(30) Tran Thi, P. T.; Pham Thi, N.; Nguyen, X. N.; Tran Van, C.; Tran Van, L.; Nguyen, T. L.; Huynh, T. N. N.; Nguyen, L. T.; Be Thi Hoang, Y.; Serpell, C. J.; Garrett, M. D.; Tran Van, S. Comparison of Organic and Conventional Production Methods in Accumulation of Biomass and Bioactive Compounds in Centella Asiatica (L.) Urban. ChemRxiv. 10.26434/chemrxiv-2023-q9vvc.

(31) Tran Van, L.; Vo Thi, Q. N.; Tran Van, C.; Tran Thi, P. T.; Pham Thi, N.; Nguyen Tuan, T.; Le Thi, T. H.; Nguyen Thi, N.; Do Thi, T.; Tran Van, S. Synthesis of Asiatic Acid Derivatives and Their Cytotoxic Activity. Medicinal Chemistry Research 2018, 27 (6), 1609–1623. 10.1007/s00044-018-2176-y.

(32) Kleifeld, O.; Doucet, A.; Prudova, A.; auf dem Keller, U.; Gioia, M.; Kizhakkedathu, J. N.; Overall, C. M. Identifying and Quantifying Proteolytic Events and the Natural N Terminome by Terminal Amine Isotopic Labeling of Substrates. Nat Protoc 2011, 6 (10), 1578–1611. 10.1038/nprot.2011.382.

(33) Oluyemi, W. M.; Samuel, B. B.; Adewumi, A. T.; Adekunle, Y. A.; Soliman, M. E. S.; Krenn, L. An Allosteric Inhibitory Potential of Triterpenes from Combretum Racemosum on the Structural and Functional Dynamics of Plasmodium Falciparum Lactate Dehydrogenase Binding Landscape. Chemistry & Biodiversity 2022, 19 (2), e202100646. 10.1002/cbdv.202100646.

(34) Feng, J.; Li, J.; Wu, L.; Yu, Q.; Ji, J.; Wu, J.; Dai, W.; Guo, C. Emerging Roles and the Regulation of Aerobic Glycolysis in Hepatocellular Carcinoma. Journal of Experimental & Clinical Cancer Research 2020, 39 (1), 126. 10.1186/s13046-020-01629-4.

(35) Huang, C. K.; Sun, Y.; Lv, L.; Ping, Y. ENO1 and Cancer. Molecular Therapy - Oncolytics 2022, 24, 288–298. 10.1016/j.omto.2021.12.026.

(36) Hamaguchi, T.; Iizuka, N.; Tsunedomi, R.; Hamamoto, Y.; Miyamoto, T.; Iida, M.; Tokuhisa, Y.; Sakamoto, K.; Takashima, M.; Tamesa, T.; Oka, M. Glycolysis Module Activated by Hypoxia-Inducible Factor 1α Is Related to the Aggressive Phenotype of Hepatocellular Carcinoma. International Journal of Oncology 2008, 33 (4), 725–731. 10.3892/ijo_00000058.

(37) Takashima, M.; Kuramitsu, Y.; Yokoyama, Y.; Iizuka, N.; Fujimoto, M.; Nishisaka, T.; Okita, K.; Oka, M.; Nakamura, K. Overexpression of Alpha Enolase in Hepatitis C Virus-Related Hepatocellular Carcinoma: Association with Tumor Progression as Determined by Proteomic Analysis. PROTEOMICS 2005, 5 (6), 1686–1692. 10.1002/pmic.200401022.

(38) Yu, S.; Li, N.; Huang, Z.; Chen, R.; Yi, P.; Kang, R.; Tang, D.; Hu, X.; Fan, X. A Novel lncRNA, TCONS_00006195, Represses Hepatocellular Carcinoma Progression by Inhibiting Enzymatic Activity of ENO1. Cell Death Dis 2018, 9 (12), 1–13. 10.1038/s41419-018-1231-4.

(39) Chan, K.; Farias, A. G.; Lee, H.; Guvenc, F.; Mero, P.; Brown, K. R.; Ward, H.; Billmann, M.; Aulakh, K.; Astori, A.; Haider, S.; Marcon, E.; Braunschweig, U.; Pu, S.; Habsid, A.; Tong, A. H. Y.; Christie-Holmes, N.; Budylowski, P.; Ghalami, A.; Mubareka, S.; Maguire, F.; Banerjee, A.; Mossman, K. L.; Greenblatt, J.; Gray-Owen, S. D.; Raught, B.; Blencowe, B. J.; Taipale, M.; Myers, C.; Moffat, J. Survival-Based CRISPR Genetic Screens across a Panel of Permissive Cell Lines Identify Common and Cell-Specific SARS-CoV-2 Host Factors. Heliyon 2023, 9 (1). 10.1016/j.heliyon.2022.e12744.

(40) Liang, J.; Zhao, H.; Diplas, B. H.; Liu, S.; Liu, J.; Wang, D.; Lu, Y.; Zhu, Q.; Wu, J.; Wang, W.; Yan, H.; Zeng, Y.-X.; Wang, X.; Jiao, Y. Genome-Wide CRISPR-Cas9 Screen Reveals Selective Vulnerability of ATRX-Mutant Cancers to WEE1 Inhibition. Cancer Research 2020, 80 (3), 510–523. 10.1158/0008-5472.CAN-18-3374.

(41) Li, W.; Liu, J.; Zhao, H. Prognostic Power of a Chaperonin Containing TCP-1 Subunit Genes Panel for Hepatocellular Carcinoma. Front. Genet. 2021, 12. 10.3389/fgene.2021.668871.

(42) Zeng, G.; Wang, J.; Huang, Y.; Lian, Y.; Chen, D.; Wei, H.; Lin, C.; Huang, Y. <p>Overexpressing CCT6A Contributes To Cancer Cell Growth By Affecting The G1-To-S Phase Transition And Predicts A Negative Prognosis In Hepatocellular Carcinoma</P>. OTT 2019, 12, 10427–10439. 10.2147/OTT.S229231.

(43) Yao, L.; Zou, X.; Liu, L. The TCP1 Ring Complex Is Associated with Malignancy and Poor Prognosis in Hepatocellular Carcinoma. International Journal of Clinical and Experimental Pathology 2019, 12 (9), 3329.

(44) Fu, R.; Jiang, S.; Guan, Z.; Li, J.; Zhang, X.; Chen, H. Comprehensive Analysis of the Expression of Chaperonin Containing TCP1 Subunits (CCTs) and Their Influence on Prognosis in Hepatocellular Carcinoma. Translational Cancer Research 2020, 9 (3), 1867. 10.21037/tcr.2020.02.20.

(45) Yang, Z.; Zhuang, L.; Szatmary, P.; Wen, L.; Sun, H.; Lu, Y.; Xu, Q.; Chen, X. Upregulation of Heat Shock Proteins (HSPA12A, HSP90B1, HSPA4, HSPA5 and HSPA6) in Tumour Tissues Is Associated with Poor Outcomes from HBV-Related Early-Stage Hepatocellular Carcinoma. International Journal of Medical Sciences 2015, 12 (3), 256. 10.7150/ijms.10735.

(46) Beck, M.; Schmidt, A.; Malmstroem, J.; Claassen, M.; Ori, A.; Szymborska, A.; Herzog, F.; Rinner, O.; Ellenberg, J.; Aebersold, R. The Quantitative Proteome of a Human Cell Line. Molecular Systems Biology 2011, 7, 549. 10.1038/msb.2011.82.

(47) Teng, D.; Ding, L.; Cai, B.; Luo, Q.; Wang, H. Interleukin-7 Enhances Anti-Tumor Activity of CD8+ T Cells in Patients with Hepatocellular Carcinoma. Cytokine 2019, 118, 115–123. 10.1016/j.cyto.2018.04.003.

(48) Tsai, S.-L. Interleukin-12 Gene Therapy for Hepatocellular Carcinoma: A Dream Come True? Journal of Gastroenterology and Hepatology 2004, 19 (10), 1097–1100. 10.1111/j.1440-1746.2004.03552.x.

(49) Zhang, H.; Xia, P.; Liu, J.; Chen, Z.; Ma, W.; Yuan, Y. ATIC Inhibits Autophagy in Hepatocellular Cancer through the AKT/FOXO3 Pathway and Serves as a Prognostic Signature for Modeling Patient Survival. Int J Biol Sci 2021, 17 (15), 4442–4458. 10.7150/ijbs.65669.

(50) Qiao, L.; Xie, N.; Li, Y.; Bai, Y.; Liu, N.; Wang, J. Downregulation of HNRNPM Inhibits Cell Proliferation and Migration of Hepatocellular Carcinoma through MAPK/AKT Signaling Pathway. Translational Cancer Research 2022, 11 (7). 10.21037/tcr-21-2484.

(51) Lee, D.; Xu, I. M.-J.; Chiu, D. K.-C.; Lai, R. K.-H.; Tse, A. P.-W.; Li, L. L.; Law, C.-T.; Tsang, F. H.-C.; Wei, L. L.; Chan, C. Y.-K.; Wong, C.-M.; Ng, I. O.-L.; Wong, C. C.-L. Folate Cycle Enzyme MTHFD1L Confers Metabolic Advantages in Hepatocellular Carcinoma. J Clin Invest 2017, 127 (5), 1856–1872. 10.1172/JCI90253.

(52) Liu, T.; Long, Q.; Li, L.; Gan, H.; Hu, X.; Long, H.; Yang, L.; Pang, P.; Wang, S.; Deng, W. The NRF2-Dependent Transcriptional Axis, XRCC5/hTERT Drives Tumor Progression and 5-Fu Insensitivity in Hepatocellular Carcinoma. Molecular Therapy - Oncolytics 2022, 24, 249–261. 10.1016/j.omto.2021.12.012.

(53) Yu, H.; Wang, H.; Xu, H.-R.; Zhang, Y.-C.; Yu, X.-B.; Wu, M.-C.; Jin, G.-Z.; Cong, W.-M. Overexpression of MTHFD1 in Hepatocellular Carcinoma Predicts Poorer Survival and Recurrence. Future Oncology 2019, 15 (15), 1771–1780. 10.2217/fon-2018-0606.

(54) Chu, Q.; Gu, X.; Zheng, Q.; Wang, J.; Zhu, H. <p>Phosphoribosyl Pyrophosphate Amido Transferase: A New Prognostic Biomarker for Hepatocellular Carcinoma</P>. IJGM 2022, 15, 353–358. 10.2147/IJGM.S340758.

(55) Kennedy, L. HNRNPM Regulates HCC Tumorigenesis and Cancer Stemness: Identification of a Novel Therapeutic Target? Cellular and Molecular Gastroenterology and Hepatology 2022, 13 (5), 1471–1473. 10.1016/j.jcmgh.2022.02.012.

(56) Zhu, G.-Q.; Wang, Y.; Wang, B.; Liu, W.-R.; Dong, S.-S.; Chen, E.-B.; Cai, J.-L.; Wan, J.-L.; Du, J.-X.; Song, L.-N.; Chen, S.-P.; Yu, L.; Zhou, Z.-J.; Wang, Z.; Zhou, J.; Shi, Y.-H.; Fan, J.; Dai, Z. Targeting HNRNPM Inhibits Cancer Stemness and Enhances Antitumor Immunity in Wnt-Activated Hepatocellular Carcinoma. Cellular and Molecular Gastroenterology and Hepatology 2022, 13 (5), 1413–1447. 10.1016/j.jcmgh.2022.02.006.

(57) Pan, Y.; Zhu, Q.; Hong, T.; Cheng, J.; Tang, X. Targeting PRKDC Activates the Efficacy of Antitumor Immunity While Sensitizing to Chemotherapy and Targeted Therapy in Liver Hepatocellular Carcinoma. Aging (Albany NY) 2024, 16 (10), 9047–9071. 10.18632/aging.205855.

(58) Zhang, Y.-N.; Zhang, W.; Hong, D.; Shi, L.; Shen, Q.; Li, J.-Y.; Li, J.; Hu, L.-H. Oleanolic Acid and Its Derivatives: New Inhibitor of Protein Tyrosine Phosphatase 1B with Cellular Activities. Bioorganic & Medicinal Chemistry 2008, 16 (18), 8697–8705. 10.1016/j.bmc.2008.07.080.

(59) Xu, X.; Wang, Y.; Wei, Z.; Wei, W.; Zhao, P.; Tong, B.; Xia, Y.; Dai, Y. Madecassic Acid, the Contributor to the Anti-Colitis Effect of Madecassoside, Enhances the Shift of Th17 toward Treg Cells via the PPARγ/AMPK/ACC1 Pathway. Cell Death Dis 2017, 8 (3), e2723–e2723. 10.1038/cddis.2017.150.

(60) Wen, X.; Sun, H.; Liu, J.; Cheng, K.; Zhang, P.; Zhang, L.; Hao, J.; Zhang, L.; Ni, P.; Zographos, S. E.; Leonidas, D. D.; Alexacou, K.-M.; Gimisis, T.; Hayes, J. M.; Oikonomakos, N. G. Naturally Occurring Pentacyclic Triterpenes as Inhibitors of Glycogen Phosphorylase: Synthesis, Structure−Activity Relationships, and X-Ray Crystallographic Studies. J. Med. Chem. 2008, 51 (12), 3540–3554. 10.1021/jm8000949.

(61) Wu, J.-J.; Ai, C.-Z.; Liu, Y.; Zhang, Y.-Y.; Jiang, M.; Fan, X.-R.; Lv, A.-P.; Yang, L. Interactions between Phytochemicals from Traditional Chinese Medicines and Human Cytochrome P450 Enzymes. Current Drug Metabolism 2012, 13 (5), 599–614.

(62) Zhang, J.; Pan, Q.-S.; Qian, X.-K.; Zhou, X.-L.; Wang, Y.-J.; He, R.-J.; Wang, L.-T.; Li, Y.-R.; Huo, H.; Sun, C.-G.; Sun, L.; Zou, L.-W.; Yang, L. Discovery of Triterpenoids as Potent Dual Inhibitors of Pancreatic Lipase and Human Carboxylesterase 1. Journal of Enzyme Inhibition and Medicinal Chemistry 2022, 37 (1), 629–640. 10.1080/14756366.2022.2029855.

(63) Nelson, K. M.; Dahlin, J. L.; Bisson, J.; Graham, J.; Pauli, G. F.; Walters, M. A. The Essential Medicinal Chemistry of Curcumin. J. Med. Chem. 2017, 60 (5), 1620–1637. 10.1021/acs.jmedchem.6b00975.

(64) Baell, J. B.; Holloway, G. A. New Substructure Filters for Removal of Pan Assay Interference Compounds (PAINS) from Screening Libraries and for Their Exclusion in Bioassays. J. Med. Chem. 2010, 53 (7), 2719–2740. 10.1021/jm901137j.

(65) Baell, J. B.; Nissink, J. W. M. Seven Year Itch: Pan-Assay Interference Compounds (PAINS) in 2017—Utility and Limitations. ACS Chem. Biol. 2018, 13 (1), 36–44. 10.1021/acschembio.7b00903.

(66) Bisson, J.; McAlpine, J. B.; Friesen, J. B.; Chen, S.-N.; Graham, J.; Pauli, G. F. Can Invalid Bioactives Undermine Natural Product-Based Drug Discovery? J. Med. Chem. 2016, 59 (5), 1671–1690. 10.1021/acs.jmedchem.5b01009.

(67) Vincent, F.; Nueda, A.; Lee, J.; Schenone, M.; Prunotto, M.; Mercola, M. Phenotypic Drug Discovery: Recent Successes, Lessons Learned and New Directions. Nat Rev Drug Discov 2022, 21 (12), 899–914. 10.1038/s41573-022-00472-w.

(68) Sadri, A. Is Target-Based Drug Discovery Efficient? Discovery and “Off-Target” Mechanisms of All Drugs. J. Med. Chem. 2023, 66 (18), 12651–12677. 10.1021/acs.jmedchem.2c01737.

