## Supplementary material for "Identification of molecular pathways involved in the anti-liver cancer activity of madecassic acid": All supplementary data

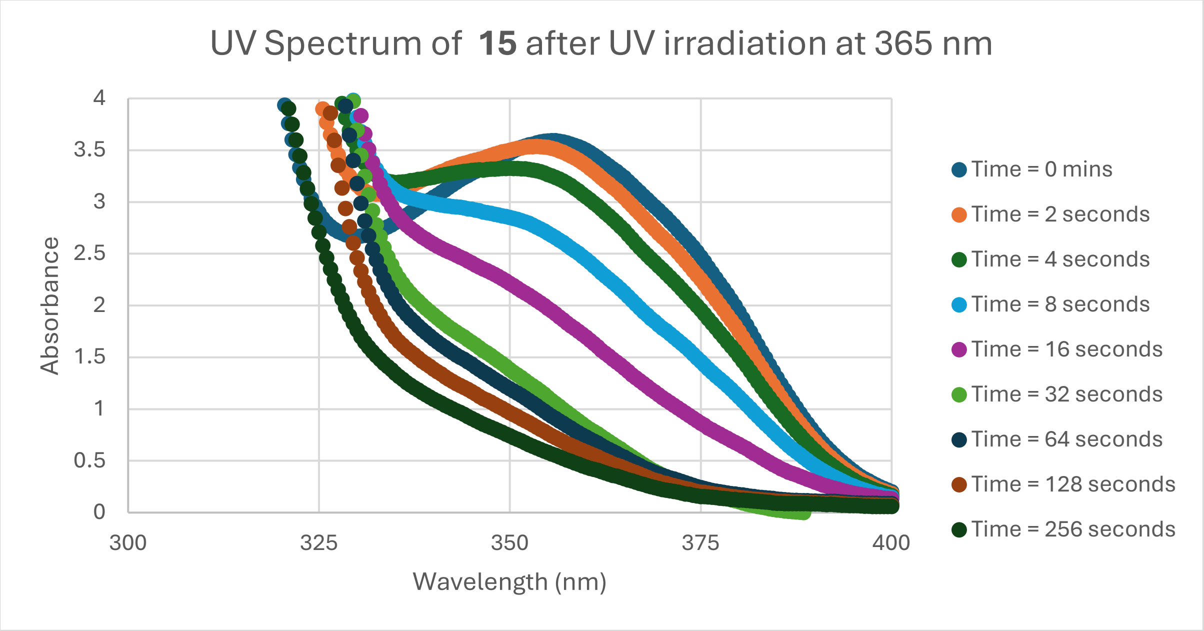


**Figure S1.** Determination of half-life of **15** by absorbance measurements after irradiation at 365 nm.

#### Biology Experimental

##### Materials:

Iscove Modified Dulbecco Medium (IMDM), Iscove Modified Dulbecco Medium with no phenol red, Fetal bovine serum (FBS) and phosphate buffered saline (PBS) were purchased from Gibco. Serological pipettes, sterile 96-well plates and T75 cell culture flasks were purchased from Sarstedt. Sterile DMSO, Sodium fluoride, sodium orthovanadate, bovine serum albumin (BSA), sodium ascorbate, β-glycerophosphate and TWEEN® 20 were purchased from Sigma-Aldrich. Trypsin-EDTA, ethylenediaminetetraacetic acid (EDTA), Streptavidin HP SpinTrap columns, sodium chloride, copper (III) sulphate pentahydrate, tris(hydroxymethyl)aminomethane (Tris), sucrose, TrisHCl, magnesium chloride, methanol, acetic acid and chloroform were purchased from ThermoFisher Scientific. HEPES, free acid and DTT (dithiothreitol) were purchased from Melford biolaboratories Ltd. Streptavidin magnetic beads and triethylammonium bicarbonate buffer (TEAB) were purchased from ThermoScientific. Tris(benzyltriazolylmethyl)amine (THPTA) was purchased from Combi-blocks. Azido-PEG3-biotin conjugate was purchased from Alfa Aesar. SDS micropellets were purchased from Fisher BioReagents. HEPG2 cell line was originally from ATCC.

##### Buffer Preparations

Lysis buffer: 50 mM HEPES pH 7.4, 250 mM NaCl, 0.1% NP40, 1mM Pefabloc/AEBSF*, 10μg/ml Aprotinin*, 20μM Leupeptin*, 1 mM DTT, 1mM EDTA, 1mM NAF, 10mM β-Glycerophosphate, 0.1mM Sodium Orthovanadate *from Boehringer/Roche proteinase inhibitor cocktail tablet.

HEPES buffer: 25 mM HEPES, 150 mM NaCl, 2 mM $MgCl_2$ ,1% SDS, pH: 7.5.

TBS-T buffer: 50 mM Tris pH 8.0, 150 mM NaCl, 0.1% TWEEN® 20.

**Instrumentation**

Absorbances measured for MTT assays were taken using 570 nm in a Perkin-Elmer Victor X4 plate reader. Cells were grown in a Thermo-scientific Forma Series II Water Jacket $CO_2$ incubator at 37 °C with 5% $CO_2$.

**Cell culture methods and equipment**

Cell lines were defrosted from cryopreserved stocks and placed into IMDM supplemented with 10% FBS (v/v). Cells were grown as adherent cultures in T75 cell culture flasks with biweekly passaging of cells and addition of fresh media. Passaging of cells involved the removal of media, a wash with 2.0 mL PBS and then addition of 0.5 mL 0.25% white trypsin (both of which were warmed in a Grant water bath to 37 °C before use after storage at 4 °C). Cells were resuspended in 13.5 mL cell media and 3 mL aliquot taken into a new T75 cell culture flask and 12 mL fresh supplemented media added. Cells were then placed in the incubator. Cell cultures were grown at 37 °C in a 5% CO2 environment. All cell handling was carried out in a sterile laminar-flow fume hood and equipment followed standard sterilisation procedure including autoclaving, 70% IMS spray and aseptic technique.

**MTT Growth Inhibition Assay**

HepG2 were maintained in IMDM supplemented with 10% FBS (v/v). Cytotoxicity of madecassic acid derivatives against the HepG2 human cancer cell line was evaluated by the 3-(4,5-Dimethylthiazol-2-yl)-2,5-Diphenyltetrazolium Bromide (MTT) method. Briefly, $3 \times 10^4$ cells.$mL^{-1}$ (190 µL) were seeded into 96-well microplates. Drugs made up in DMSO were diluted with media (to a maximum concentration of 10% DMSO) to give the desired compound concentrations. Drugs made up in the DMSO/Media mixture (10 µL) was added to each well to give the desired final drug concentrations and a final maximum DMSO concentration of 0.5%. Each concentration was tested in triplicate. After incubation at 37 °C in 5% $CO_2$ for 48 h, culture medium was carefully removed and replaced with fresh 1.2 mM MTT IMDM without phenol red or FBS and the plates incubated for another 4 h. Then the liquid in the well was removed, and DMSO (100 µL) was added to each well, along with thorough mixing to dissolve formazan produced. Ellipticine was used as positive control. The absorbance was recorded on a microplate reader at the wavelength of 570 nm and compared to an undrugged control. Cell viability was measured and $IC_{50}$ values were calculated based on the Raw data software. Growth curves were plotted using Excel and Graphpad Prism 9.

**Photoaffinity Crosslinking Procedure**

HEPG2 cells were pelleted at 20 million cells per pellet. 8 pellets were lysed using lysis buffer (250 µL per pellet) and the pellets were combined into 4 pairs (500 µL each) and each precleared of biotin interacting molecules by passage through a streptavidin HP SpinTrap column. Lysates were combined and cell lysates were diluted with lysis buffer to 3 mg/mL protein (determined with BSA by Bradford assay). Precleared cell lysates (485 µL) were aliquoted into Eppendorf tubes along with 2.5 µL of either DMSO or 3.7 mM madecassic acid **1** in DMSO (for competition control) were added to the lysates, and they were incubated for 1 hour at 4 °C in darkness. Then, 12.5 µL of either DMSO, **14** or **15** (2 mM in DMSO) was added and the lysates incubated for a further 1 hour at 4 °C in darkness. Samples were irradiated through the top of the Eppendorf with an Alonefire SV18 12W 365nm UV Torch placed at a 4 cm distance for 5 min. Samples were then snap frozen at −80 °C for further use. Protocol was repeated in triplicate on separate days.

**Photoaffinity Click Reaction Procedure**

Crosslinked lysates previously prepared were defrosted and to each 500 µL lysate was added 1000 µL PBS buffer and 50 µL of 1 mM biotin azide **16** in DMSO followed by brief vortexing. The following were all added followed by brief vortexing after the addition of each: 100 µL of 100 mM THPTA solution, 100 µL of 20 mM $CuSO_4$ solution and 100 µL of 300 mM sodium ascorbate solution. The reaction was protected from light at room temperature for 3 hours. To each lysate was added the following followed by brief vortexing after each: 3000 µL methanol, 750 µL chloroform and 2000 µL distilled $H_2O$. Lysates were centrifuged at 13,000-20,000 × g, upper aqueous layer removed without disturbing interface layer containing proteins. 2250 µL methanol added, vortexed briefly, centrifuged at 13,000-20,000 × g and supernatant removed. Methanol wash of pellet was repeated. Supernatant of protein pellet completely removed, and the pellet was allowed to air dry for at least 15 minutes.

**Photoaffinity Magnetic Bead Pulldown Procedure**

To a 2.0 mL Eppendorf was added 125 µL of vortexed Pierce magnetic streptavidin beads. Beads were collected to the side of the tube with a magnet and supernatant removed. 1500 µL of washing buffer (TBS-T) was added and beads again collected to the side of the tube using a magnet and supernatant removed. Crosslinked and clicked protein pellets were dissolved in 1500 µL of HEPES buffer containing 1% SDS and added to magnetic beads and they were incubated together for 1 hour at room temperature. Beads were collected to the side of the tube using a magnet and supernatant removed. 750 µL of washing buffer (TBS-T) was added and beads collected to the side of the tube using a magnet and supernatant removed. Beads were washed with TBS-T (750 µL) an additional 2 times (total 3 time) and supernatant removed after each washing. 750 µL TEAB buffer (1.0 M, pH 8.5) was added and beads collected to the side of the tube using a magnet and supernatant removed. Beads were washed with TEAB buffer (750 µL) an additional 2 times (total 3 time) and supernatant removed after each washing. Beads were then stored at −80 °C before analysis at the ICR.

**Photoaffinity Proteomics Procedure**

For sample preparation and TMT labelling, beads from chemical pulldown experiments were resuspended in a buffer containing 100 µL of 0.1 M triethylammonium bicarbonate (TEAB), 5 mM tris(2-carboxyethyl)phosphine (TCEP), and 10 mM iodoacetamide. Samples were sonicated and incubated for 45 minutes at room temperature. Proteins were digested by adding 10 µL of trypsin stock solution (500 ng/µL in 0.1% formic acid, Pierce, #90059) and incubating for 2 hours at 37 °C. This was followed by overnight incubation at room temperature in a shaker at 600 rpm. Beads were magnetised, and the supernatant containing the resulting peptides was dried using a centrifugal vacuum concentrator.

Dried peptides were resuspended in 25 µL of 100 mM TEAB and labelled with a tandem mass tag (TMTpro) multiplex reagent vial (Thermo Scientific, #A44520) according to the manufacturer's instructions. All TMT-labelled samples were combined in equal amounts in a single tube and dried using a centrifugal vacuum concentrator.

For high-pH reversed-phase peptide fractionation, pooled TMT-labelled peptides were fractionated using the Pierce™ High-pH Reversed-Phase Peptide Fractionation Kit (Catalog No. 84868). Proteolytic digests were loaded onto spin columns, where peptides were bound and desalted. A step gradient of acetonitrile (5% to 50%) in 0.1% triethylamine was applied to elute peptides into eight fractions. Fractions were dried and re-suspended in 0.1% formic acid for LC-MS analysis to improve peptide separation and protein identification.

For liquid chromatography–mass spectrometry, peptide analysis was performed on a Dionex UltiMate 3000 UHPLC system coupled with an Orbitrap Ascend Tribrid mass spectrometer (Thermo Scientific). Peptides were separated using a nanoEase M/Z Peptide BEH C18 column (75 µm × 25 cm, 1.7 µm) at 45 °C. Mobile phase A was 0.1% formic acid, and mobile phase B was 80% acetonitrile with 0.1% formic acid. The gradient separation method was as follows: 100-minute gradient to 35% B, followed by 5 minutes to 95% B, 5 minutes isocratic at 95% B, re-equilibration to 5% B over 5 minutes, and 5 minutes isocratic at 5% B.

Mass spectrometry data were acquired using TMT-SPS-$MS^3$ with online real-time database search (RTS). Precursors between 400 and 1600 m/z were selected with a resolution of 120,000, a standard automatic gain control (AGC) target of $2 \times 10^5$, and a maximum injection time (IT) of 50 ms, using the top-speed mode (3-second cycle time). Ion trap $MS^2$ scans were acquired using HCD fragmentation (32% collision energy), standard AGC target, maximum IT of 35 ms, and turbo scan rate. RTS was enabled, and quantitative TMT-SPS-$MS^3$ scans were collected with a resolution of 45,000, an AGC target of $1 \times 10^5$, and a maximum IT of 200 ms.

For protein database search and quantification, the SEQUEST-HT search engine was used to analyse the acquired mass spectra in Proteome Discoverer 2.4 (Thermo Scientific). The precursor mass tolerance was set at 20 ppm, and the fragment ion mass tolerance was 0.02 Da. Spectra were searched for fully tryptic peptides, allowing up to two missed cleavages. TMT on lysine residues and peptide N-termini (+229.163 Da) and carbamidomethylation of cysteine residues (+57.0215 Da) were set as static modifications. Oxidation of methionine (+15.9949 Da) and deamidation of asparagine and glutamine (+0.9848 Da) were set as variable modifications.

Peptide confidence was estimated using Percolator, with the peptide false discovery rate (FDR) set to 0.01, based on the q-value and a decoy database search. All spectra were searched against the UniProt-SwissProt *Homo sapiens* proteome, appended with common contaminants. The Reporter Ion Quantifier node included TMT quantification with an integration window tolerance of 15 ppm and an integration method based on the most confident centroid peak at the $MS^3$ level. Only unique peptides were used for quantification, and protein groups were considered for peptide uniqueness. Peptides with an average reporter signal-to-noise ratio >3 were used for protein quantification. Correction for isotopic impurities in reporter ion quantification values was applied. Only spectra with at least 50% of the SPS fragment ions matching the identified peptides were used for quantification.

**RNA Extraction Protocol**

HepG2 were maintained in IMDM supplemented with 10% FBS (v/v). HepG2 cells were seeded into T25 cell dishes (2.0 mL of $1.15 \times 10^6$ cells.$mL^{-1}$). Either madecassic acid (3 mM in DMSO) or **7** (50 mM in DMSO) (33 µL or 11 µL + 22 µL DMSO) were added to each of the T25 flasks. Additional IMDM supplemented with 10% FBS (v/v) (2967 µL) was added to each T25 flask (total volume 5000 µL). The flasks were gently swirled and incubated at 37 °C in 5% $CO_2$ for either 6 or 24 hours. Media from T25 flasks was removed, a wash with 2.0 mL PBS and then addition of 0.5 mL 0.25% white trypsin (both of which were warmed in a Grant water bath to 37 °C before use after storage at 4 °C). Cells were pelleted at 500 × g and supernatant discarded. RNA extraction was performed using Monarch® total RNA miniprep kit according to the use manual.

**Chemistry Experimental**

**Materials:**

Deuterated solvents were purchased from Cambridge Isotope Laboratories Inc. Chloroform, $CH_2Cl_2$, diethyl ether, DMF, DMSO, ethanol, ethyl acetate (EtOAc), formic acid, hydrochloric acid, hexane, methanol, sodium hydrogen carbonate and magnesium sulphate were purchased from Fisher Scientific. Potassium carbonate was purchased from Thermo Fisher Scientific. HCl (4 M in dioxane), 3-bromoanisole, methyl trifluoroacetate, hydroxylamine hydrochloride, DMAP, triethylamine, dichloromethyl methyl ether, EDC.HCl and silver oxide were purchased from Fluorochem. Sodium hydroxide, dry pyridine, dry $CH_2Cl_2$, dry THF, dry ethanol, 7 M $NH_3$ in methanol, titanium tetrachloride, tetrabutylammonium iodide and n-butyl amine were purchased from Acros Organics. Benzyl bromide, Tosyl chloride and boron tribromide (1 M in $CH_2Cl_2$) were purchased from Alfa Aesar.

**Instrumentation**

A Bruker AVII 400 MHz spectrometer was used to record NMR spectra, and each spectrum was calibrated to the known chemical shift of the residual solvent peak of the deuterated solvent used. Chemical shifts were reported in part per million (ppm) and J coupling values were reported in Hz. Proton NMR spectra were obtained at 400 MHz and carbon-13 spectra were obtained at 101 MHz. Spectral data was processed using MestReNova software. UV-Vis spectra were recorded using a NanoDrop™ One/OneC Microvolume UV-Vis Spectrophotometer. Data was processed using Excel and Graphpad Prism 9. UV irradiation was carried out using an Alonefire SV18 12W 365nm UV Torch. Electrospray mass spectrometry data was obtained using a Thermo MSQPlus instrument fitted with a Zorbax SB-C18 5 μm 3.0 x 150 mm column using $H_2O$ + 0.1% formic acid and MeOH + 0.1% formic acid or $H_2O$ + 0.1% TFA and MeCN + 0.1% TFA mobile phases. Data was analysed using Chromeleon™ Chromatography Data System (CDS) Software. A Bruker micrOTOF-Q LCMS system was used to obtain high-resolution mass spectrometry data, samples were dissolved in HPLC-grade methanol and injected using direct injection mode with a mobile phase system of 50:50 MeOH and $H_2O$. Data was processed using Bruker Compass Hystar software.

**Synthesis of 2**

Madecassic Acid (201.6 mg, 0.40 mmol) was dissolved in 6 mL of dry pyridine under nitrogen and acetic anhydride (0.13 mL, 1.40 mmol) was injected into the reaction slowly. After 24 hours, 10 mL pyridine was added, and additional acetic anhydride (4 mL, 42.3 mmol) was added to the reaction. After 48 hours, TLC (after staining with vanillin) showed complete consumption of starting material. The reaction was transferred to a separating funnel, EtOAc (25 mL) was added, and the reaction organic layer was washed with 1 M $HCl_{(aq)}$ (3 × 50 mL). The organic layer was dried over anhydrous $MgSO_4$, filtered and the solvent removed under reduced pressure. Crude product was purified using gradient silica gel column chromatography (0-50%, EtOAc:$CH_2Cl_2$) giving **2** as a white needle crystalline solid (183 mg 72.6% yield).

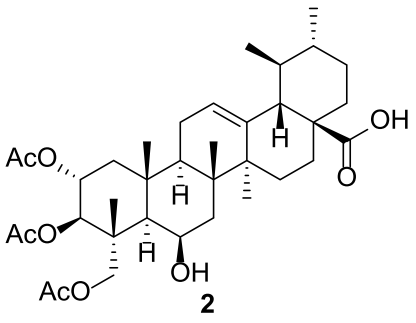


**2**

**$^1$H NMR** (400 MHz, $CDCl_3$) δ 5.27 (d, *J* = 3.6 Hz, 1H), 5.22 (td, *J* = 11.3, 4.8 Hz, 2H), 5.00 (d, *J* = 10.3 Hz, 1H), 4.33 (d, *J* = 3.8 Hz, 1H), 3.93 (d, *J* = 11.9 Hz, 1H), 3.70 (d, *J* = 12.0 Hz, 1H), 2.19 (d, *J* = 11.2 Hz, 1H), 2.15 – 0.80 (m, 46H). **$^{13}$C NMR** (101 MHz, $CDCl_3$) δ 183.9, 171.0, 170.6, 170.6, 137.3, 125.7, 75.0, 70.0, 68.0, 65.5, 52.5, 48.3, 48.0, 47.9, 45.9, 42.5, 40.8, 39.2, 38.9, 38.7, 37.4, 36.7, 30.7, 28.0, 24.2, 23.6, 23.4, 21.3, 21.2, 21.2, 21.0, 20.9, 18.8, 18.6, 17.1, 15.5, 14.3. **HREI-MS**: *m/z* calculated for $C_{36}H_{54}O_9Na$ [M+Na$^+$] 653.3660; observed 653.3672.

**Synthesis of S1**

HATU (131.1 mg, 0.334 mmol) and **2** (175.5 mg, 0.278 mmol) were dissolved in 5 mL dry $CH_2Cl_2$ and DIPEA ( 0.145 mL, 0.835 mmol) was slowly added to the reaction and it was stirred for 1 hour. *Tert*-butyl 10-aminodecylcarbamate (152.8 mg, 0.556 mmol) was dissolved in dry $CH_2Cl_2$ (5 mL) and slowly added to the reaction and it was left to stir overnight. After 22 hours, the reaction was transferred to a separating funnel and diluted with $CH_2Cl_2$ (30 mL). It was then extracted with 1 M $HCl_{(aq)}$ (3 × 50 mL) and brine (1 × 50 mL). Dried over anhydrous $MgSO_4$, filtered and reduced under vacuum. The crude product was purified by isocratic column Chromatography (25%:75%, EtOAc:$CHCl_3$) to yield **S1** as a white solid (236 mg, 95.8%).

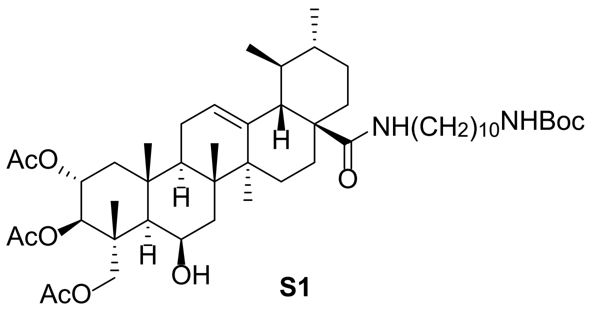


**S1**

**$^1$H NMR** (400 MHz, $CDCl_3$) δ 5.85 (t, *J* = 5.4 Hz, 1H), 5.34 (t, *J* = 3.5 Hz, 1H), 5.22 (td, *J* = 11.0, 4.7 Hz, 1H), 5.01 (d, *J* = 10.3 Hz, 1H), 4.57 (s, 1H), 4.34 (t, *J* = 3.8 Hz, 1H), 3.94 (d, *J* = 11.9 Hz, 1H), 3.71 (d, *J* = 12.0 Hz, 1H), 3.23 (dq, *J* = 13.3, 6.8 Hz, 1H), 3.14 – 2.93 (m, 3H), 2.17 – 0.77 (m, 72H). **$^{13}$C NMR** (101 MHz, $CDCl_3$) δ 177.9, 170.9, 170.5, 170.5, 139.4,

125.5, 79.1, 75.0, 70.1, 67.4, 65.4, 54.2, 48.1, 47.9, 47.8, 46.0, 43.1, 42.5, 40.6, 39.9, 39.6, 39.2, 38.8, 38.8, 37.4, 37.3, 31.0, 30.2, 29.5, 29.5, 29.3, 29.3, 29.3, 29.2, 28.5, 27.9, 27.2, 26.9, 24.9, 23.5, 23.3, 21.3, 21.2, 21.0, 20.9, 18.7, 18.3, 17.3, 15.4. **HREI-MS**: *m/z* calculated for $C_{51}H_{84}N_2O_{10}Na$ [M+Na$^+$] 907.6018; observed 907.6052.

**Synthesis of 3**

**S1** (131.2 mg, 0.148 mmol) was dissolved in $CH_2Cl_2$ (3.7 mL), 4 M HCl in dioxane (3.7 mL, 14.8 mmol) was added to the solution at 0 °C. The reaction was stirred and left to warm to room temperature overnight. After 21 hours, solvent was removed under reduced pressure. The residue was washed into a separating funnel with a mixture of $CH_2Cl_2$ (50 mL) and sat. $NaHCO_{3(aq)}$ (50 mL) and aqueous layer removed. It was then extracted with a further sat. $NaHCO_{3(aq)}$ (2 × 50 mL) and brine (1 × 50 mL). Dried over anhydrous $MgSO_4$, filtered and reduced under vacuum. Extracted product was clean by NMR and required no further purification yielding **3** (105.8 mg, 90.9%).

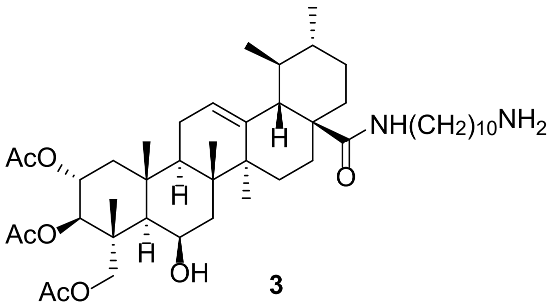


**3**

**$^1$H NMR** (400 MHz, $CDCl_3$) δ 5.92 (t, *J* = 5.4 Hz, 1H), 5.34 (d, *J* = 3.7 Hz, 1H), 5.21 (td, *J* = 10.9, 4.5 Hz, 1H), 5.00 (d, *J* = 10.3 Hz, 1H), 4.83 (s, 2H), 4.31 (d, *J* = 5.1 Hz, 1H), 3.94 (d, *J* = 11.7 Hz, 1H), 3.70 (dd, *J* = 12.5, 2.6 Hz, 1H), 3.28 – 2.94 (m, 3H), 2.85 (s, 2H), 2.25 – 0.73 (m, 62H). **$^{13}$C NMR** (101 MHz, $CDCl_3$) δ 178.1, 171.1, 170.7, 170.6, 139.4, 125.5, 75.0, 70.2, 67.2, 65.4, 54.1, 48.1, 47.9, 47.8, 46.0, 43.8, 43.1, 42.5, 40.4, 39.9, 39.7, 39.2, 38.8, 37.4, 37.3, 31.0, 29.5, 29.4, 29.3, 29.3, 29.2, 29.1, 27.8, 27.1, 26.7, 24.9, 23.5, 23.3, 21.3, 21.3, 21.0, 20.9, 18.7, 18.3, 17.3, 15.4.

**Synthesis of 4**

**3** (52.0 mg, 0.0662 mmol) was dissolved in MeOH (5 mL). 4 M $NaOH_{(aq)}$ solution (0.080 mL, 0.331 mmol) was injected into the reaction, and it was left to stir overnight. After 20 hours, reaction was transferred to a separating funnel, EtOAc (50 mL) was added. The organic layer was extracted with 1 M $NaOH_{(aq)}$ (3 × 50 mL). Dried over anhydrous $MgSO_4$, filtered and reduced under vacuum. Extracted product was pure and required no further purification, yielding **4** (43.6 mg, 99.9%).

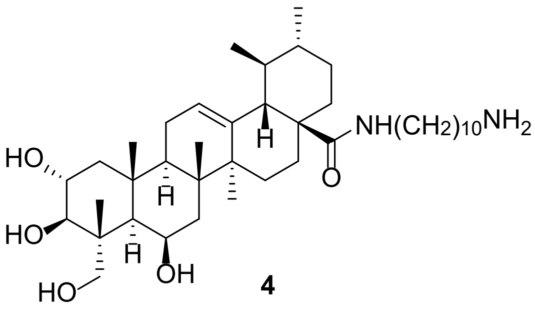

**4**

[1]**H NMR** (400 MHz, Methanol-*d*$_4$) δ 5.39 (t, *J* = 3.6 Hz, 1H), 4.37 (t, *J* = 3.4 Hz, 1H), 3.74 (ddd, *J* = 11.3, 9.3, 4.3 Hz, 1H), 3.58 (d, *J* = 11.1 Hz, 1H), 3.43 (d, *J* = 11.1 Hz, 1H), 3.30 (d, *J* = 8.0 Hz, 1H), 3.21 – 3.01 (m, 2H), 2.75 – 2.66 (m, 2H), 2.20 – 0.79 (m, 54H).[13]**C NMR** (101 MHz, Methanol-*d*$_4$) δ 180.1, 139.7, 127.1, 78.0, 69.6, 68.3, 65.7, 54.3, 50.2, 49.1, 48.9, 44.8, 43.9, 42.0, 41.2, 40.9, 40.8, 40.3, 40.1, 38.8, 38.5, 32.2, 32.0, 30.7, 30.7, 30.7, 30.6, 30.5, 30.3, 28.9, 28.3, 27.9, 25.2, 24.5, 24.1, 21.6, 19.3, 19.3, 17.7, 15.3. **HREI-MS**: *m/z* calculated for $C_{40}H_{71}N_2O_5$ [M+H$^+$] 659.5357; observed 659.5392.

**Synthesis of 5**

Benzoic Acid (20.3 mg, 0.161 mmol) and HATU (49.8 mg, 0.129 mmol) were dissolved in dry $CH_2Cl_2$ (5 mL) and DIPEA (0.093 mL, 0.536 mmol) was slowly added to the reaction, and it was stirred for 1 hour. **3** (88.0 mg, 0.107 mmol) was dissolved in dry $CH_2Cl_2$ (5 mL) and slowly added to the reaction and it was left to stir overnight. After 22 hours and consumption of all starting material, the reaction was diluted with $CH_2Cl_2$ (40 mL) and transferred to a separating funnel. The organic layer was extracted with sat. $NaHCO_{3(aq)}$ (2 × 50 mL) and 1 M $HCl_{(aq)}$ (2 × 50 mL). Organic layer was collected, dried over anhydrous $MgSO_4$, filtered and solvent removed under reduced pressure. Crude product was purified by gradient column chromatography (0-60%, EtOAc:$CHCl_3$) to give the desired product as a single fraction of **5** as a white solid (64.0 mg, 67.2%).

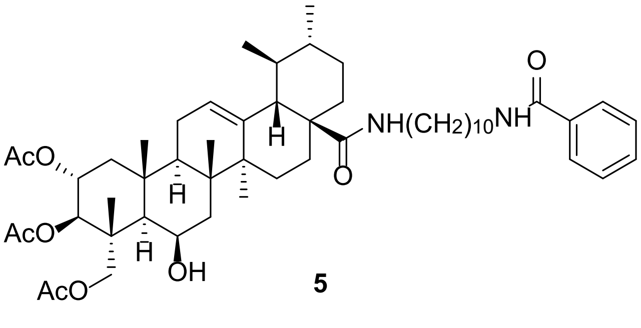

**5**

[1]**H NMR** (400 MHz, $CDCl_3$) δ 7.76 (dt, *J* = 6.9, 1.5 Hz, 2H), 7.49 – 7.43 (m, 1H), 7.43 – 7.37 (m, 2H), 6.53 (t, *J* = 5.7 Hz, 1H), 5.90 (t, *J* = 5.3 Hz, 1H), 5.33 (t, *J* = 3.5 Hz, 1H), 5.21 (td, *J* = 11.0, 4.7 Hz, 1H), 5.00 (d, *J* = 10.3 Hz, 1H), 4.33 (d, *J* = 3.7 Hz, 1H), 3.93 (d, *J* = 11.9 Hz, 1H), 3.69 (d, *J* = 12.0 Hz, 1H), 3.41 (td, *J* = 7.4, 5.7 Hz, 2H), 3.21 (td, *J* = 13.1, 6.7 Hz, 1H), 3.07 – 2.95 (m, 1H), 2.45 – 0.71 (m, 62H). [13]**C NMR** (101 MHz, $CDCl_3$) δ 178.1, 171.0, 170.7, 170.5, 167.8, 139.4, 134.8, 131.4, 128.6, 127.1, 125.5, 74.9, 70.1, 67.4, 65.3, 54.1, 48.1,

47.9, 47.8, 45.9, 43.1, 42.5, 40.6, 40.3, 39.9, 39.7, 39.1, 38.8, 37.4, 37.2, 31.0, 29.7, 29.5, 29.5, 29.4, 29.2, 29.1, 27.8, 27.2, 27.0, 24.8, 23.5, 23.3, 21.3, 21.2, 21.0, 20.9, 18.7, 18.3, 17.3, 15.4. **HREI-MS**: *m/z* calculated for $C_{53}H_{81}N_2O_9$ [M+H$^+$] 889.5937; observed 889.5972.

**Synthesis of 6**

**5** (25.6 mg, 0.0288 mmol) was dissolved in MeOH (5 mL). 4 M $NaOH_{(aq)}$ solution (0.036 mL, 0.144 mmol) was injected into the reaction, and it was left to stir overnight. After 22 hours, solvent was removed under reduced pressure, reaction was transferred to a separating funnel with EtOAc (30 mL). The organic layer was extracted with $H_2O$ (3 × 30 mL). Dried over anhydrous $MgSO_4$, filtered and reduced under vacuum. Crude product was subjected to isocratic column chromatography (100%, EtOAc) to yield **6** as a white solid (16.8 mg, 76.5%).

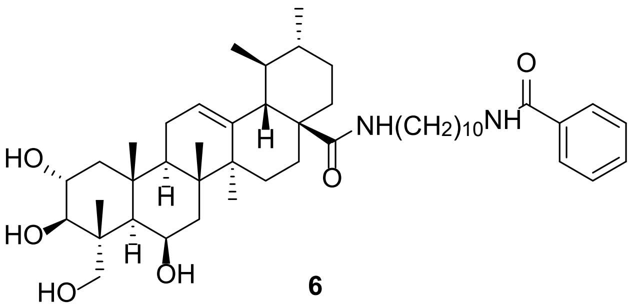


**6**

**$^1$H NMR** (400 MHz, Methanol-*d*$_4$) δ 7.83 – 7.78 (m, 2H), 7.56 – 7.49 (m, 1H), 7.48 – 7.42 (m, 2H), 7.14 (t, *J* = 5.6 Hz, 1H), 5.39 (t, *J* = 3.6 Hz, 1H), 4.38 (t, *J* = 3.8 Hz, 1H), 3.74 (ddd, *J* = 11.3, 9.5, 4.5 Hz, 1H), 3.58 (d, *J* = 11.1 Hz, 1H), 3.43 (d, *J* = 11.0 Hz, 1H), 3.37 (t, *J* = 7.2 Hz, 2H), 3.29 (d, *J* = 8.1 Hz, 1H), 3.21 – 3.02 (m, 2H), 2.19 – 0.83 (m, 54H). **$^{13}$C NMR** (101 MHz, Methanol-*d*$_4$) δ 180.2, 170.2, 139.7, 135.9, 132.5, 129.5, 128.2, 127.2, 78.1, 69.7, 68.3, 65.7, 54.3, 50.3, 49.2, 48.6, 48.4, 44.8, 43.9, 41.2, 41.1, 40.9, 40.9, 40.3, 40.1, 38.8, 38.5, 32.0, 30.7, 30.7, 30.5, 30.5, 30.5, 30.3, 28.9, 28.3, 28.1, 25.2, 24.5, 24.1, 21.6, 19.3, 19.3, 17.7, 15.3. **HREI-MS**: *m/z* calculated for $C_{47}H_{75}N_2O_6$ [M+H$^+$] 763.5620; observed 763.5651.

**Synthesis of S2**

**S1** (404.0 mg, 0.456 mmol) was dissolved in MeOH (10 mL). 4 M $NaOH_{(aq)}$ solution (0.115 mL, 0.456 mmol) was injected into the reaction and it was left to stir overnight. Reaction was transferred to a separating funnel, EtOAc (50 mL) added. The organic layer was extracted with $H_2O$ (3 × 50 mL). Dried over anhydrous $MgSO_4$, filtered and reduced under vacuum. Crude product was purified by isocratic column Chromatography (100%, EtOAc) to give **S2** as a pure white solid (300.3 mg, 86.7%).

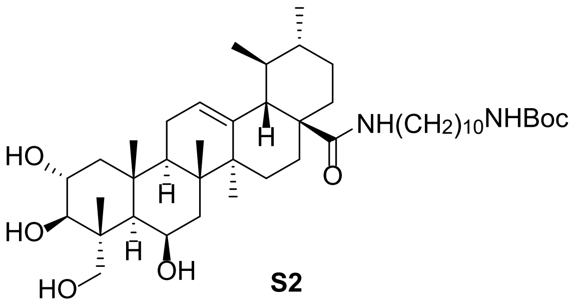

**S2**

**$^1$H NMR** (400 MHz, Methanol-$d_4$) δ 7.08 (t, *J* = 5.6 Hz, 1H), 5.39 (t, *J* = 3.6 Hz, 1H), 4.38 (t, *J* = 3.6 Hz, 1H), 3.75 (ddd, *J* = 11.3, 9.5, 4.6 Hz, 1H), 3.58 (d, *J* = 11.2 Hz, 1H), 3.44 (d, *J* = 11.1 Hz, 1H), 3.30 (d, *J* = 9.7 Hz, 1H), 3.19 – 3.04 (m, 2H), 3.01 (t, *J* = 7.1 Hz, 2H), 2.20 – 0.82 (m, 63H). **$^{13}$C NMR** (101 MHz, Methanol-$d_4$) δ 180.1, 158.5, 139.7, 127.1, 79.7, 78.1, 69.6, 68.3, 65.9, 54.4, 50.2, 49.1, 48.9, 48.7, 44.8, 43.9, 41.4, 41.2, 40.9, 40.9, 40.3, 40.1, 38.8, 38.5, 32.0, 31.0, 30.7, 30.7, 30.5, 30.5, 30.3, 28.9, 28.8, 28.3, 27.9, 25.3, 24.5, 24.1, 21.6, 19.3, 19.3, 17.7, 15.3. **HREI-MS**: *m/z* calculated for $C_{45}H_{79}N_2O_7$ [M+H$^+$] 759.5882; observed 759.5914.

**Synthesis of S3, S4 and S5**

Potassium *tert*-Butoxide (179.2 mg, 1.58 mmol), **S2** (600.0 mg, 0.790 mmol) and tetrabutylammonium iodide (29.0 mg, 0.0790 mmol) were dissolved in 30 mL dry Toluene and stirred for 30 mins. Benzyl Bromide (0.113 mL, 0.948 mmol) was slowly injected into the reaction mixture. The reaction was left to react overnight. After 21 hours, the reaction was diluted with EtOAc (50 mL) and washed with brine (3 × 50 mL). The organic phase was dried over anhydrous magnesium sulphate, filtered and solvent removed under reduced pressure. Crude products were purified by gradient column chromatography (0-100%, EtOAc:$CH_2Cl_2$) to give the products **S3** (29.0 mg, 4.3%), **S4** (32.0 mg, 4.8%), **S5** (230.0 mg, 34.3%) and recovered starting material **S2** (178.0 mg, 29.7%).

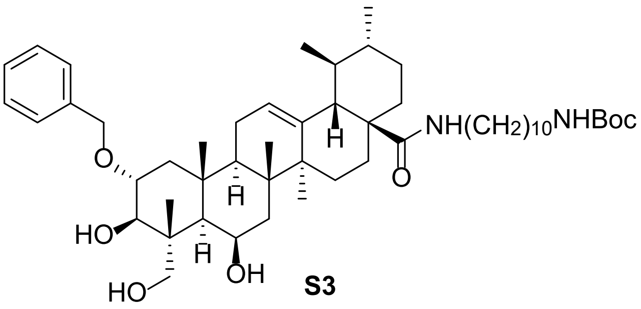

**S3**

**$^1$H NMR** (400 MHz, Methanol-$d_4$) δ 7.42 – 7.38 (m, 2H), 7.36 – 7.30 (m, 2H), 7.29 – 7.24 (m, 1H), 7.13 (t, *J* = 5.6 Hz, 1H), 5.38 (t, *J* = 3.6 Hz, 1H), 4.68 – 4.60 (m, 2H), 4.38 (t, *J* = 4.0 Hz, 1H), 3.67 – 3.58 (m, 2H), 3.48 – 3.40 (m, 2H), 3.20 – 3.04 (m, 2H), 3.01 (t, *J* = 7.1 Hz, 2H), 2.20 – 0.75 (m, 63H). **$^{13}$C NMR** (101 MHz, Methanol-$d_4$) δ 180.2, 158.6, 140.4, 139.7, 129.3, 129.3, 128.6, 127.2, 79.8, 78.5, 76.8, 72.9, 68.2, 65.5, 54.3, 49.3, 48.6, 47.3, 45.0, 44.0, 41.4, 41.2, 40.9, 40.9, 40.3, 40.1, 38.8, 38.4, 32.0, 31.0, 30.7, 30.7, 30.7, 30.6, 30.5, 30.3,

28.9, 28.8, 28.3, 27.9, 25.2, 24.5, 24.1, 21.6, 19.3, 19.2, 17.7, 15.4. **HREI-MS**: *m/z* calculated for $C_{52}H_{84}N_2O_7Na$ [M+Na$^+$] 871.6171; observed 871.6168.

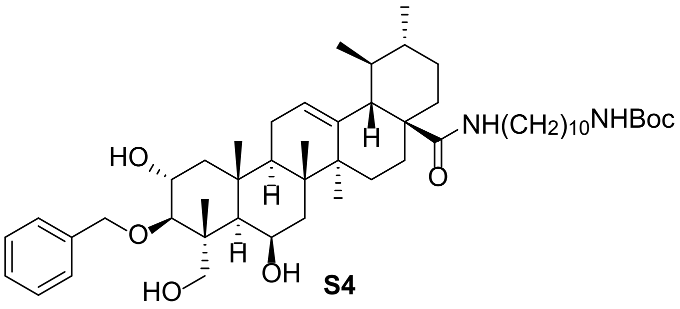


**S4**

**$^1$H NMR** (400 MHz, Methanol-*d*$_4$) δ 7.42 – 7.38 (m, 2H), 7.35 – 7.29 (m, 2H), 7.27 – 7.21 (m, 1H), 7.14 (t, *J* = 5.6 Hz, 1H), 5.44 – 5.37 (m, 1H), 5.01 (d, *J* = 11.3 Hz, 1H), 4.65 (d, *J* = 11.2 Hz, 1H), 4.41 (t, *J* = 3.6 Hz, 1H), 3.94 (ddd, *J* = 11.5, 9.5, 4.5 Hz, 1H), 3.59 – 3.46 (m, 2H), 3.30 (d, *J* = 7.4 Hz, 1H), 3.20 – 3.04 (m, 2H), 3.01 (t, *J* = 7.1 Hz, 2H), 2.21 – 0.87 (m, 63H). **$^{13}$C NMR** (101 MHz, Methanol-*d*$_4$) δ 180.2, 158.6, 141.1, 139.7, 129.3, 129.2, 128.5, 128.2, 127.2, 86.7, 79.8, 76.8, 70.5, 68.2, 65.1, 54.3, 51.0, 49.3, 48.4, 45.9, 44.0, 41.4, 41.2, 40.9, 40.9, 40.3, 40.2, 38.8, 38.4, 32.0, 31.0, 30.7, 30.7, 30.6, 30.5, 30.3, 28.9, 28.8, 28.4, 27.9, 25.3, 24.6, 24.1, 21.6, 19.3, 19.3, 17.7, 16.2. **HREI-MS**: *m/z* calculated for $C_{52}H_{84}N_2O_7Na$ [M+Na$^+$] 871.6171; observed 871.6168.

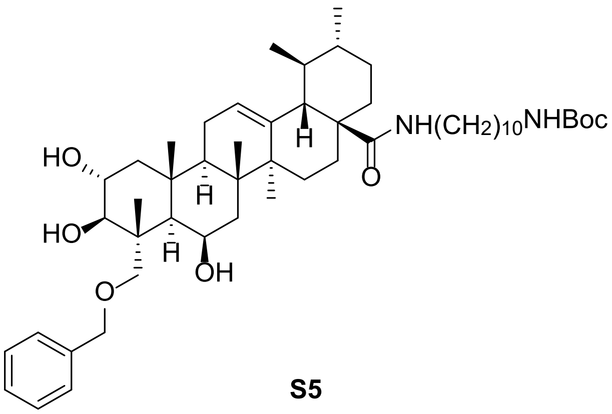


**S5**

**$^1$H NMR** (400 MHz, Methanol-*d*$_4$) δ 7.34 – 7.22 (m, 5H), 7.13 (t, *J* = 5.6 Hz, 1H), 5.39 (t, *J* = 3.6 Hz, 1H), 4.56 (d, *J* = 11.9 Hz, 1H), 4.39 (d, *J* = 11.9 Hz, 1H), 4.24 (t, *J* = 3.8 Hz, 1H), 3.74 (ddd, *J* = 11.3, 9.6, 4.5 Hz, 1H), 3.51 (d, *J* = 9.3 Hz, 1H), 3.42 (d, *J* = 9.7 Hz, 1H), 3.33 (d, *J* = 9.3 Hz, 1H), 3.20 – 3.04 (m, 2H), 3.01 (t, *J* = 7.1 Hz, 2H), 2.21 – 0.83 (m, 63H). **$^{13}$C NMR** (101 MHz, Methanol-*d*$_4$) δ 180.1, 158.5, 140.2, 139.6, 129.3, 128.5, 128.5, 128.5, 127.2, 79.7, 78.3, 73.9, 69.6, 68.3, 54.3, 50.4, 49.2, 48.9, 48.9, 44.8, 43.9, 41.4, 41.3, 40.9, 40.3, 40.1, 38.8, 38.4, 32.0, 31.0, 30.7, 30.7, 30.7, 30.6, 30.5, 30.3, 28.9, 28.8, 28.3, 27.9, 25.3, 24.5, 24.1, 21.6, 19.3, 19.3, 17.8, 15.5. **HREI-MS**: *m/z* calculated for $C_{52}H_{85}N_2O_7$ [M+H$^+$] 849.6351; observed 849.6363.

**Synthesis of 7**

**S3** (20.3 mg, 0.0239 mmol) was dissolved in dry $CH_2Cl_2$ (3 mL), 4 M HCl in dioxane (1.0 mL, 3.98 mmol) was added to the solution. The reaction was then stirred overnight. After 20 hours, reaction solvent was removed under reduced pressure. Reaction mixture redissolved in EtOAc (30 mL) and transferred to a separating funnel. Organic layer was extracted with 1 M $NaOH_{(aq)}$ (3 × 50 mL). Dried over anhydrous $MgSO_4$, filtered and reduced under vacuum to give pure product **7** as a white solid (13.7 mg. 76.5% Yield).

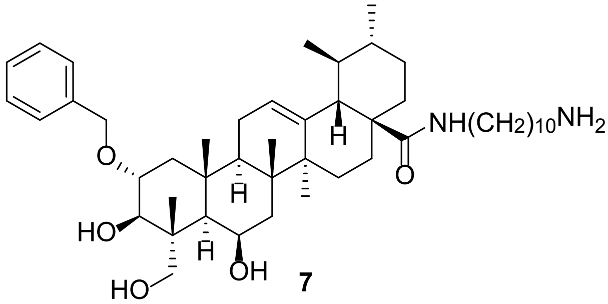

**7**

**$^1$H NMR** (400 MHz, Methanol-*d*$_4$) δ 7.44 – 7.10 (m, 5H), 5.40 (d, *J* = 14.1 Hz, 1H), 4.72 – 4.59 (m, 2H), 4.37 (s, 1H), 3.76 – 3.39 (m, 4H), 3.19 – 3.01 (m, 2H), 2.90 – 2.82 (m, 2H), 2.23 – 0.79 (m, 53H). **$^{13}$C NMR** (101 MHz, Methanol-*d*$_4$) δ 180.1, 140.4, 129.3, 129.2, 129.2, 128.6, 127.1, 78.5, 76.7, 72.8, 68.3, 65.5, 54.3, 49.1, 48.6, 46.7, 45.0, 44.0, 41.0, 40.9, 40.8, 40.3, 40.1, 38.8, 38.4, 33.1, 32.0, 30.8, 30.8, 30.7, 30.5, 30.5, 30.3, 29.3, 28.2, 27.6, 24.2, 24.1, 22.0, 21.6, 19.3, 19.2, 17.7, 17.3, 15.4. **HREI-MS**: *m/z* calculated for $C_{47}H_{77}N_2O_5$ [M+H$^+$] 749.5827; observed 749.5836.

**Synthesis of 8**

**S4** (27.5 mg, 0.0353 mmol) in dry $CH_2Cl_2$ (3.0 mL) and cooled to 0 °C. 4 M HCl in dioxane (1.0 mL, 3.53 mmol) was added to the solution at 0 °C and it was stirred until consumption of starting material. After 25 hours, reaction mixture was diluted with EtOAc (30 mL) and transferred to a separating funnel. Organic layer was extracted with 1 M NaOH (3 × 50 mL). Product was dried over anhydrous $MgSO_4$, filtered and reduced under vacuum giving pure product **8** as a white solid (24.2 mg, 99.8%).

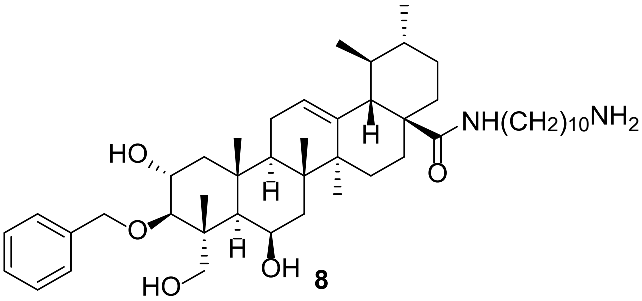

**8**

**$^1$H NMR** (400 MHz, Methanol-*d*$_4$) δ 7.34 – 7.27 (m, 2H), 7.24 – 7.19 (m, 2H), 7.19 – 7.10 (m, 1H), 5.29 (s, 1H), 4.90 (d, *J* = 11.0 Hz, 1H), 4.55 (d, *J* = 11.2 Hz, 1H), 4.31 (s, 1H), 3.89 – 3.77 (m, 1H), 3.54 – 3.37 (m, 2H), 3.20 (d, *J* = 3.1 Hz, 1H), 3.12 – 2.89 (m, 2H), 2.78 (t, *J* = 7.7 Hz, 2H), 2.12 – 0.58 (m, 53H). **$^{13}$C NMR** (101 MHz, Methanol-*d*$_4$) δ 180.2, 141.0, 139.7, 129.2,

128.6, 128.3, 127.1, 86.7, 76.8, 70.5, 68.2, 65.1, 54.3, 51.0, 49.1, 48.4, 45.9, 44.5, 44.0, 40.9, 40.8, 40.8, 40.3, 40.2, 38.8, 38.4, 33.1, 32.0, 30.8, 30.8, 30.7, 30.6, 30.3, 28.9, 28.3, 27.5, 25.2, 24.6, 24.1, 23.8, 21.6, 19.3, 17.7, 16.2, 14.5. **HREI-MS**: *m/z* calculated for $C_{47}H_{77}N_2O_5$ [M+H$^+$] 749.5827; observed 749.5840.

**Synthesis of 9**

**S5** (109.0 mg, 0.128 mmol) was dissolved in $CH_2Cl_2$ (3.2 mL), 4 M HCl in dioxane (3.2 mL, 12.8 mmol) was added to the solution at 0 °C. The reaction was stirred and left to warm to room temperature overnight. After 27 hours, the reaction was transferred to a separating funnel, diluted with $CH_2Cl_2$ (50 mL) and extracted with sat. $NaHCO_{3(aq)}$ (2 × 100 mL) and brine (1 × 100 mL). Dried over anhydrous $MgSO_4$, filtered and reduced under vacuum to give pure **9** as a white solid (87.4 mg, 90.9%).

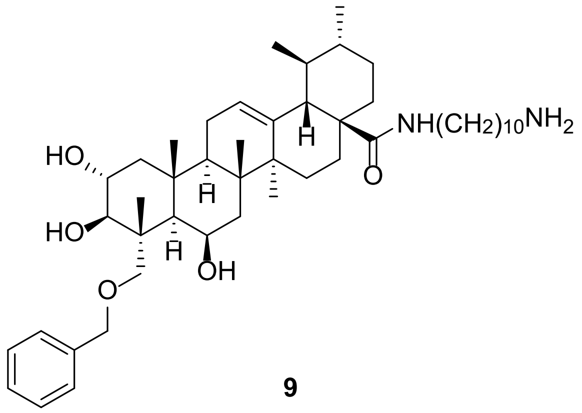


**9**

**$^1$H NMR** (400 MHz, Methanol-*d*$_4$) δ 7.36 – 7.21 (m, 5H), 5.39 (t, *J* = 3.6 Hz, 1H), 4.56 (d, *J* = 11.9 Hz, 1H), 4.39 (d, *J* = 11.9 Hz, 1H), 4.24 (t, *J* = 3.9 Hz, 1H), 3.73 (ddd, *J* = 11.3, 9.6, 4.6 Hz, 1H), 3.51 (d, *J* = 9.3 Hz, 1H), 3.42 (d, *J* = 9.7 Hz, 1H), 3.33 (d, *J* = 9.6 Hz, 1H), 3.19 – 3.02 (m, 2H), 2.72 – 2.65 (m, 2H), 2.19 – 0.82 (m, 53H). **$^{13}$C NMR** (101 MHz, Methanol-*d*$_4$) δ 180.1, 140.2, 139.6, 129.3, 128.5, 128.5, 127.2, 78.3, 74.0, 74.0, 69.6, 68.3, 54.3, 50.4, 49.3, 48.9, 44.8, 43.9, 42.1, 41.3, 40.9, 40.8, 40.3, 40.1, 38.8, 38.5, 32.4, 32.0, 30.7, 30.7, 30.6, 30.6, 30.6, 30.3, 28.9, 28.3, 27.9, 25.2, 24.5, 24.1, 21.6, 19.3, 19.3, 17.7, 15.5. **HREI-MS**: *m/z* calculated for $C_{47}H_{77}N_2O_5$ [M+H$^+$] 749.5827; observed 749.5839.

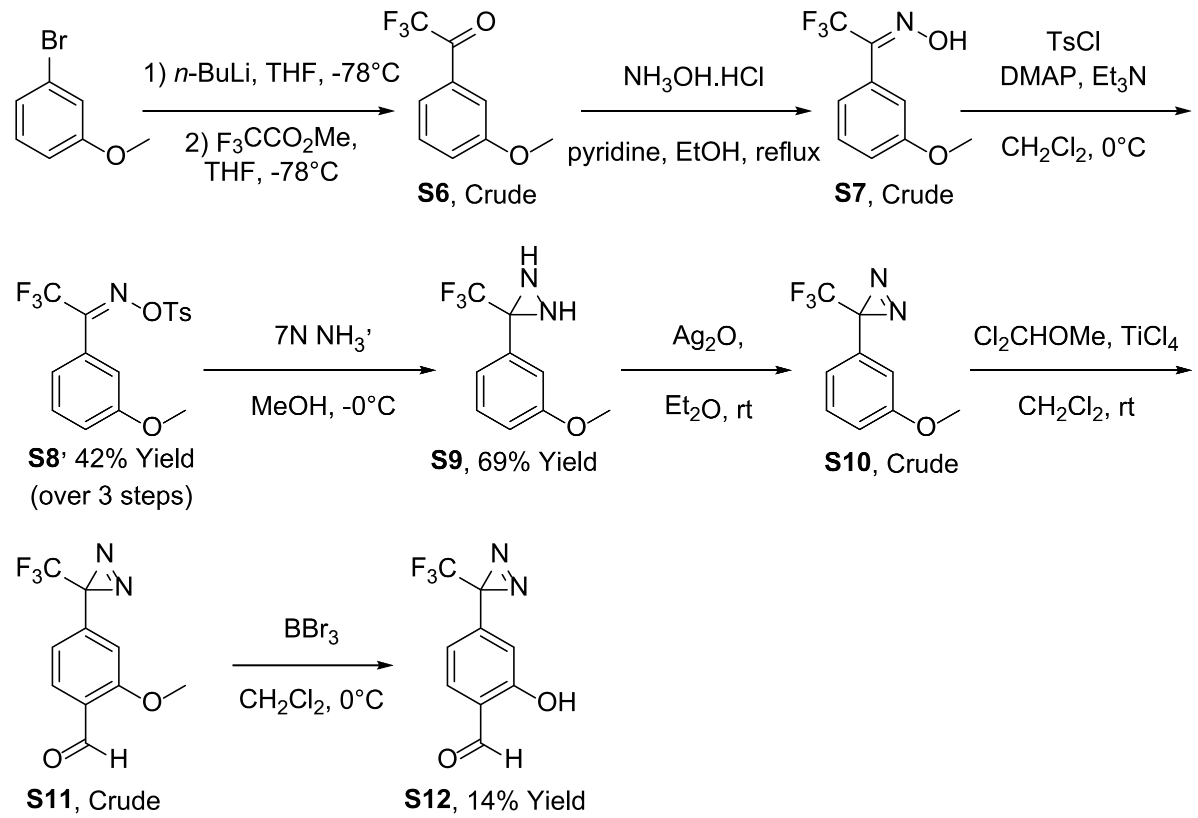


**Scheme S2.** Synthesis of **S12** from 3-bromoanisole.

**Synthesis of S6**

A solution of 3-bromoanisole (2.0 mL, 15.79 mmol) in dry THF (32 mL) was cooled to −84 °C under an argon atmosphere. Using an oven dried needle, *n*-Butyl lithium (2.5 M in hexane, 9.48 mL, 23.69 mmol) was injected into the reaction mixture over several minutes and the reaction was left stirring for 2 hours at −84 °C. In a separate reaction vessel, methyl trifluoroacetate (2.07 mL, 20.53 mmol) in dry THF (13 mL) was cooled to −84 °C under an argon atmosphere. The trifluoroacetate mixture was injected into the reaction mixture very slowly over 30 minutes. The reaction was allowed to stir at −84 °C for a further 2 hours (checking with TLC for consumption of starting material). The reaction was slowly quenched by injecting sat. $NH_4Cl_{(aq)}$ (12 mL) at −84 °C. The quenched reaction was allowed to warm to room termination before being diluted with diethyl ether (25 mL) and extracted with $H_2O$ (2 × 15 mL). Organic layer dried over anhydrous $MgSO_4$, filtered, solvent reduced under vacuum and dried on high vacuum overnight giving crude product **S6** as a colourless oil (2.492 g) and used as is for the next reaction crude.

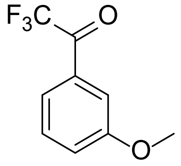


**S6**

**$^1$H NMR** (400 MHz, $CDCl_3$) δ 7.66 (ddd, *J* = 7.8, 2.5, 1.4 Hz, 1H), 7.57 (dd, *J* = 2.0 Hz, 1H), 7.45 (dd, *J* = 8.0 Hz, 1H), 7.25 (ddd, *J* = 8.3, 2.7, 0.9 Hz, 1H), 3.88 (s, 3H). **$^{13}$C NMR** (101 MHz, $CDCl_3$) 180.5 (q, *J* = 35.0 Hz), 160.1, 131.2, 130.3, 122.9 (q, *J* = 2.7 Hz), 122.4, 116.8 (q, *J* = 291.2 Hz), 114.1 (q, *J* = 1.8 Hz), 55.7.

**Synthesis of S7**

Crude **S6** (2.462 g, 12.06 mmol) was dissolved in dry pyridine (12 mL) and dry EtOH (6 mL). Hydroxylamine hydrochloride (1.060 g, 15.25 mmol) dissolved in dry pyridine (12 mL) and dry EtOH (6 mL) was added to the reaction and the reaction was refluxed at 85 °C for 9 hours. Upon consumption of the starting material, the solvent was removed under vacuum and the resulting residue partitioned between $Et_2O$ (25 mL) and $H_2O$ (25 mL). The organic layer was then washed with of 1 M $HCl_{(aq)}$ (3 × 25 mL), dried over $MgSO_4$, filtered and the solvent removed under reduced pressure. Dried on high vacuum for 1 hour to give **S7** as a clear volatile oil (2.393 g) and used for next reaction directly crude.

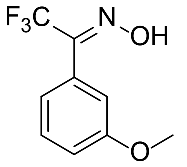


**S7**

**$^1$H NMR** (400 MHz, $CDCl_3$) δ 9.00 (s, 1H), 7.40 (ddd, *J* = 7.7, 7.7, 1.1 Hz, 1H), 7.10 – 7.06 (m, 1H), 7.06 – 7.00 (m, 2H), 3.84 (s, 3H). **$^{13}$C NMR** (101 MHz, $CDCl_3$) δ 159.6, 147.8 (q, *J* = 32.5 Hz), 129.9, 127.3, 120.9, 120.7 (q, *J* = 274.7 Hz), 116.4, 114.4, 55.5. **HREI-MS**: *m/z* calculated for $C_9H_9F_3NO_2$ [M+H$^+$] 220.0580; observed 220.0583.

**Synthesis of S8**

DMAP (0.141 g, 1.156 mmol) and TsCl (3.081 g, 16.18 mmol) were dissolved in $CH_2Cl_2$ (30 mL) and cooled to 0 °C. $Et_3N$ (1.27 mL, 17.34 mmol) was added to the solution and it was stirred for 30 mins. Crude **S7** (2.359 g) was dissolved in $CH_2Cl_2$ (30 mL) at 0 °C. The solution of DMAP, TsCl and $Et_3N$ was added to solution **S7** at 0 °C and left to slowly warm up to room temperature. The reaction was left at room temperature overnight After 20 hours, the reaction was transferred to a separating funnel and washed with 1 M $HCl_{(aq)}$ (3 × 50 mL), dried over $MgSO_4$, filtered and reduced under vacuum to give a crude white solid. Crude product was purified by isocratic column chromatography (30% $CH_2Cl_2$/ 70% hexane) followed by recrystallised from boiling ethanol and the mother liquid was recrystallised again to give additional product giving **S8** as a white solid (2.498 g, 42.4 % yield (over 3 steps)).

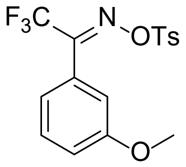

**S8**

**[1]H NMR** (400 MHz, $CDCl_3$) δ 7.93 – 7.85 (m, 2H), 7.42 – 7.35 (m, 3H), 7.05 (ddd, *J* = 8.4, 2.6, 1.0 Hz, 1H), 6.97 – 6.91 (m, 1H), 6.88 (dd, *J* = 2.1, 2.1 Hz, 1H), 3.82 (s, 3H), 2.48 (s, 3H). **[13]C NMR** (101 MHz, $CDCl_3$) δ 159.6, 154.0 (q, *J* = 33.6 Hz), 146.1, 131.2, 130.0, 129.9, 129.3, 125.6, 120.5, 119.7 (q, *J* = 278.2 Hz), 117.2, 113.9, 55.4, 21.8. **HREI-MS**: *m/z* calculated for $C_{16}H_{14}F_3NO_4SNa$ [M+Na$^+$] 396.0488; observed 396.0493.

**Synthesis of S9**

**S8** (2.270 g, 6.080 mmol) was dissolved in 7 M methanolic ammonia (25 mL, 175 mmol) at 0 °C and left in ice bath to warm to room temperature over the weekend. The reaction was monitored via mini extractions ($Et_2O$ and sat. $NaHCO_{3(aq)}$) followed by TLC. After 72 hours, the reaction was opened to the air and nitrogen gas was bubbled through the reaction mixture to remove excess ammonia. Sat. $NaHCO_{3(aq)}$ (3 mL) was added and a white precipitate formed. The precipitate was filtered off, $H_2O$ (50 mL) added, and the crude product was extracted with $CH_2Cl_2$(3 × 30 mL), dried over anhydrous $MgSO_4$, filtered and concentrated under reduced pressure to give **S9** as a crude product. The crude **S9** was purified by isocratic column chromatography (10%:90%, $Et_2O$:Hexane) to yield the desired diaziridine product **S9** (921.0 mg, 69.4%) as a white solid and the hydrazone side product **S13** (159.0 mg, 12.0%) as a white solid and the imine side product **S14** (66.0 mg, 5.3%) as a white solid.

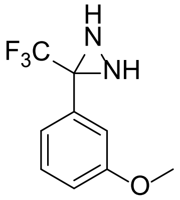

**S9**

**[1]H NMR** (400 MHz, $CDCl_3$) δ 7.33 (dd, *J* = 8.0, 8.0 Hz, 1H), 7.22 – 7.18 (m, 1H), 7.15 (dd, *J* = 2.1, 2.1 Hz, 1H), 6.97 (ddd, *J* = 8.3, 2.7, 1.0 Hz, 1H), 3.82 (s, 3H), 2.78 (s, 1H), 2.25 (s, 1H). **[13]C NMR** (101 MHz, $CDCl_3$) δ 159.8, 133.1, 130.0, 123.6 (q, *J* = 278.3 Hz), 120.4, 116.0, 113.8, 58.2 (q, *J* = 36.0 Hz), 55.5. **HREI-MS**: *m/z* calculated for $C_9H_{10}F_3N_2O$ [M+H$^+$] 219.0740; observed 219.0747.

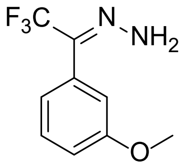

**S13**

**$^{1}$H NMR** (400 MHz, $CDCl_3$) δ 7.30 – 7.22 (m, 1H), 6.67 (ddd, *J* = 8.3, 2.5, 0.9 Hz, 1H), 6.56 – 6.47 (m, 2H), 5.00 (s, 2H), 3.79 (s, 3H). **$^{13}$C NMR** (101 MHz, $CDCl_3$) δ 161.0, 147.4 (q, *J* = 6.1 Hz), 145.6 (q, *J* = 32.4 Hz), 130.7, 118.3 (q, *J* = 278.1 Hz), 112.8, 110.5, 106.5, 55.4.

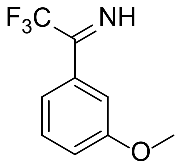


**S14**

**$^{1}$H NMR** (400 MHz, $CDCl_3$) δ 9.14 (s, 1H), 7.40 (dd, *J* = 7.9, 7.9 Hz, 1H), 7.09 (d, *J* = 7.7 Hz, 1H), 7.07 – 7.01 (m, 2H), 3.84 (s, 3H). **$^{13}$C NMR** (101 MHz, $CDCl_3$) δ 159.6, 147.7 (q, *J* = 32.5 Hz), 129.9, 127.2, 120.9, 120.7 (q, *J* = 274.7 Hz), 116.4, 114.4, 55.5.

**Synthesis of S10**

Silver(I) oxide (1696 mg, 15.58 mmol) was dissolved in $Et_2O$ (5 mL). **S9** (146.0 mg, 0.6692 mmol) was dissolved in $Et_2O$ (5 mL) and added to the silver oxide dropwise under nitrogen. Caution, addition of **S9** directly to silver(I) oxide in the absence of solvent results in a violent reaction. After 96 hours, the reaction was filtered, dried over anhydrous $MgSO_4$ and filtered again, all whilst avoiding contact with light. The crude product **S10** (144.6 mg) was taken forward directly to the following reaction.

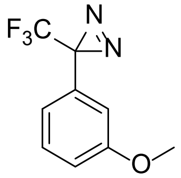


**S10**

**$^{1}$H NMR** (400 MHz, $CDCl_3$) δ 7.30 (ddd, *J* = 8.3, 7.8, 0.4 Hz, 1H), 6.94 (ddd, *J* = 8.4, 2.5, 0.9 Hz, 1H), 6.77 (ddd, *J* = 7.9, 1.8, 0.9 Hz, 1H), 6.71 – 6.65 (m, 1H), 3.80 (s, 3H). **$^{13}$C NMR** (101 MHz, $CDCl_3$) δ 159.9, 130.7, 130.1, 122.2 (q, *J* = 274.6 Hz), 118.8 (q, *J* = 1.6 Hz), 115.3, 112.3 (q, *J* = 1.3 Hz), 55.4, 28.5 (q, *J* = 40.4 Hz).

**Synthesis of S12 via S11**

Crude **S10** (144.6 mg, 0.6689 mmol) was dissolved in dry $CH_2Cl_2$ (6 mL). $TiCl_4$ (0.110 mL, 1.00 mmol) was added to the reaction at 0 °C. Dichloromethyl methyl ether (0.091 mL, 1.00 mmol) was then slowly added to the reaction at 0 °C. The reaction was allowed to warm to room temperature for 2 hours before cooling back to 0 °C and quenching with $H_2O$ (1 mL). The organic phase was diluted with $CH_2Cl_2$ (15 mL) and extracted with $H_2O$ (2 × 10 mL), sat. $NaHCO_{3(aq)}$ (2 × 10 mL) and $H_2O$ again (2 × 10 mL). Organic layer collected, dried over anhydrous $MgSO_4$, filtered and reduced under vacuum yielding **S11** as a crude product (163.4 mg), which was taken forward imminently to the next reaction. Crude **S11**

(163.4 mg, 0.6692 mmol) was dissolved in dry $CH_2Cl_2$ (3 mL) at 0 °C under nitrogen. A solution of 1 M boron tribromide in $CH_2Cl_2$ (0.74 mL, 0.736 mmol) was slowly injected and the reaction stirred at 0 °C under nitrogen. After 1.5 hours, the reaction was quenched with $H_2O$ (5 mL), $CH_2Cl_2$ (15 mL) added, and the organic layer washed with brine (3 × 20 mL). The organic layer was then collected and dried over anhydrous $MgSO_4$, filtered and reduced under vacuum. The crude product was purified by isocratic column chromatography (10%:90%, $Et_2O$:Hexane) to yield **S12** (21.4 mg, 13.9% yield (over 3 steps)), mixed with $Et_2O$ (22.6 mg). As **S12** was volatile and $Et_2O$ presented no complications in the following reaction, product was carried forward as is.

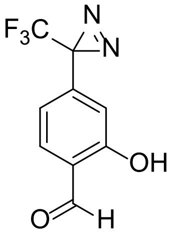


**S12**

**$^1$H NMR** (400 MHz, $CDCl_3$) δ 11.04 (s, 1H), 9.92 (d, $J$ = 0.5 Hz, 1H), 7.59 (dd, $J$ = 7.9, 0.7 Hz, 1H), 6.82 – 6.75 (m, 2H). **$^{13}$C NMR** (101 MHz, $CDCl_3$) δ 196.0, 161.5, 138.2, 134.1, 121.8 (q, $J$ = 274.8 Hz), 121.0, 117.4 (q, $J$ = 1.5 Hz), 116.0 (d, $J$ = 1.7 Hz), 29.7 (q, $J$ = 29.5 Hz).

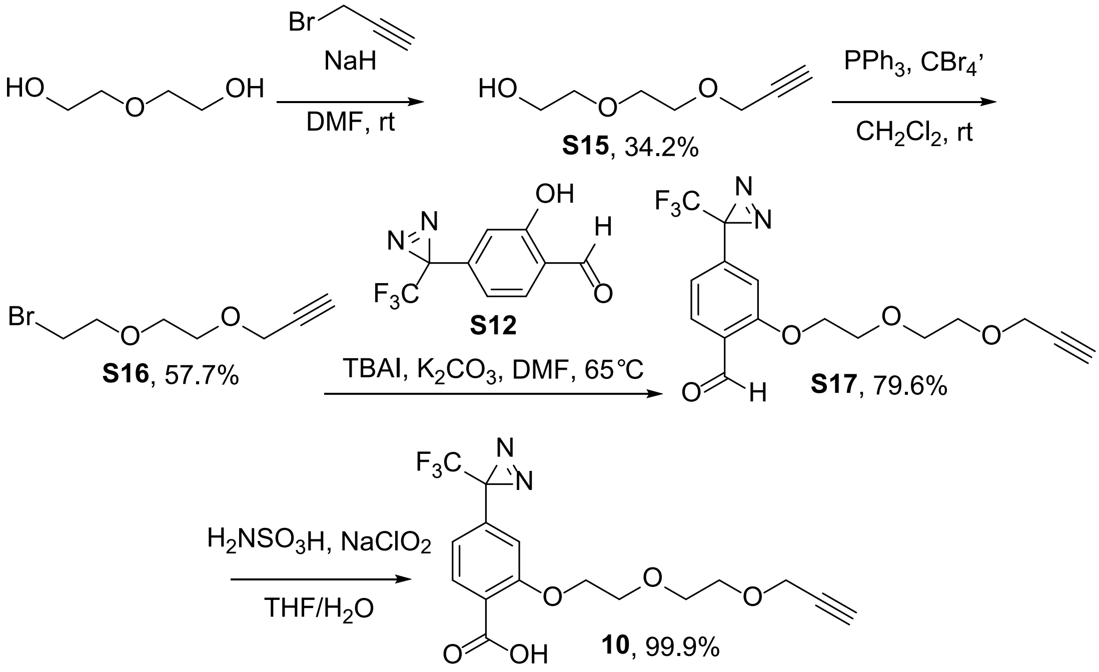


**Scheme S3.** Synthesis of **10** from diethylene glycol.

**Synthesis of S15**

Sodium hydride (60% wt in oil, 0.63 g, 15.80 mmol) was dissolved in anhydrous DMF (80 mL) and cooled to 0 °C. Diethylene glycol (1.0 mL, 10.53 mmol) was injected into the reaction and the mixture stirred for 30 min at 0 °C. Then, propargyl bromide (80% in toluene, 1.4 mL, 13.17 mmol) was added dropwise and the resulting mixture was stirred for overnight at rt. After 20 hours, MeOH (10 mL) was added. The mixture was concentrated under reduced pressure and the residue was dissolved in EtOAc (40 mL). The organic layer was washed with saturated $NH_4Cl_{(aq)}$ (40 mL), the aqueous layer was extracted with EtOAc (2 × 40 mL). The organic layers were combined, dried over anhydrous $MgSO_4$, filtered and reduced under vacuum. The crude residue was purified by isocratic column chromatography (50%:50%, EtOAc:Hexane) to give the corresponding **S15** (519.2 mg, 34.2% yield).

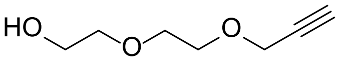


**S15**

**$^1$H NMR** (400 MHz, $CDCl_3$) δ 4.20 (d, *J* = 2.4 Hz, 2H), 3.75 – 3.66 (m, 6H), 3.62 – 3.58 (m, 2H), 2.48 (s, 1H), 2.44 (t, *J* = 2.4 Hz, 1H). **$^{13}$C NMR** (101 MHz, $CDCl_3$) δ 79.5, 74.9, 72.6, 70.3, 69.2, 61.8, 58.5.

**Synthesis of S16**

Starting material **S15** (179.7 mg, 1.246 mmol) and $CBr_4$ (655.0 mg, 1.957 mmol) were dissolved in $CH_2Cl_2$ (5 mL) and $PPh_3$ (492.3 mg, 1.870 mmol) in $CH_2Cl_2$ (5 mL) was added slowly while the mixture was stirred at 0 °C. The resulting mixture was stirred at rt overnight. After 19 hours, reaction had not gone to completion so additional $CBr_4$ (628.6 mg, 1.89 mmol) and $PPh_3$ (490.2 mg, 1.87 mmol) in dry $CH_2Cl_2$ (5 mL) was added. The reaction was left for an additional 24 hours. After 43 hours (total), the mixture was concentrated under reduced pressure onto silica for column loading. The crude product purified by isocratic column chromatography (25:75%, $Et_2O$:Hexane). Product was further purified by a second attempt at isocratic column chromatography (10%:90%, $Et_2O$:Hexane) to give pure **S16** (148.9 mg, 57.7% yield).

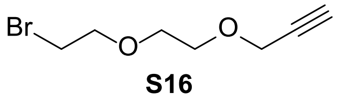


**S16**

**$^1$H NMR** (400 MHz, $CDCl_3$) δ 4.21 (d, *J* = 2.4 Hz, 2H), 3.80 (t, *J* = 6.3 Hz, 2H), 3.73 – 3.67 (m, 4H), 3.47 (t, *J* = 6.3 Hz, 2H), 2.44 (t, *J* = 2.4 Hz, 1H). **$^{13}$C NMR** (101 MHz, $CDCl_3$) δ 79.7, 74.8, 71.4, 70.5, 69.2, 58.6, 30.3.

**Synthesis of S17**

Alkyne **S16** (102.8 mg, 0.478 mmol), potassium carbonate (134.9 mg, 0.869 mmol), Diazirine **S12** (98.4 mg, 0.434 mmol) and TBAI (21.2 mg, 0.0574 mmol) were weighed into a round bottom flask and dissolved in dry DMF (10 mL). The mixture was heated at 65 °C overnight under nitrogen. After 20 hours, the reaction mixture was transferred to a separating funnel, diluted with EtOAc (10 mL) and washed with $H_2O$ (3 × 20 mL) and brine (1 × 20 mL). Organic layer was collected, dried over anhydrous magnesium sulphate, filtered and reduced under vacuum to give product **S17** as a pure yellow oil (121.3 mg, 79.6%).

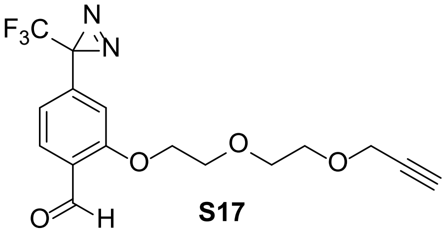


**S17**

**$^1$H NMR** (400 MHz, $CDCl_3$) δ 10.47 (d, *J* = 0.8 Hz, 1H), 7.83 (d, *J* = 8.1 Hz, 1H), 6.83 (ddt, *J* = 8.2, 1.7, 0.9 Hz, 1H), 6.72 (d, *J* = 1.6 Hz, 1H), 4.27 – 4.23 (m, 2H), 4.20 (d, *J* = 2.4 Hz, 2H), 3.96 – 3.88 (m, 2H), 3.78 – 3.68 (m, 4H), 2.44 (t, *J* = 2.4 Hz, 1H). **$^{13}$C NMR** (101 MHz, $CDCl_3$) δ 189.0, 161.1, 136.7, 128.9, 125.7, 121.9 (q, *J* = 274.8 Hz), 119.0 (q, *J* = 1.7 Hz), 111.0 (q, *J* = 1.7 Hz), 79.5, 74.9, 70.9, 69.5, 69.2, 68.6, 58.6, 28.7 (q, *J* = 40.7 Hz).

**Synthesis of 10**

**S17** (120.0 mg, 0.3368 mmol) was dissolved in THF (8 mL) under nitrogen and sulfamic acid (75.3 mg, 0.623 mmol) dissolved in $H_2O$ (2 mL) was injected into the reaction. After stirring for 5 mins, sodium chlorite solution (25% w/v, 0.183 mL, 0.606 mmol) was injected into the reaction and it was left to stir at room temperature under nitrogen. After 4 hours, the reaction was transferred to a separating funnel, diluted with $H_2O$ (25 mL) and extracted with $CH_2Cl_2$ (3 × 25 mL). The organic phases were combined, extracted with brine (50 mL), dried over anhydrous magnesium sulphate, filtered and solvent removed under reduced pressure yielding pure **10** (125.3 mg, 99.9%).

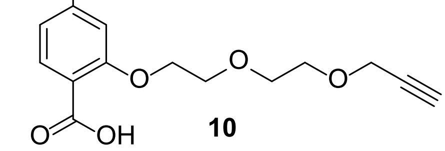


**10**

**$^1$H NMR** (400 MHz, $CDCl_3$) δ 9.33 (s, 1H), 8.13 (d, *J* = 8.3 Hz, 1H), 6.91 (ddd, *J* = 8.3, 1.8, 0.9 Hz, 1H), 6.75 (d, *J* = 1.8 Hz, 1H), 4.39 – 4.32 (m, 2H), 4.16 (d, *J* = 2.4 Hz, 2H), 3.96 – 3.89 (m, 2H), 3.78 – 3.65 (m, 4H), 2.41 (t, *J* = 2.4 Hz, 1H). **$^{13}$C NMR** (101 MHz, $CDCl_3$) δ 164.8, 157.6, 135.9, 134.2, 121.8 (q, *J* = 274.9 Hz), 120.3 (q, *J* = 1.7 Hz), 119.8, 111.6 (q, *J* = 1.5 Hz), 79.4, 74.9, 70.6, 69.6, 69.2, 68.6, 58.5, 28.4 (q, *J* = 40.8 Hz).

**Synthesis of S18**

Madecassic acid **1** (1201.8 mg, 2.38 mmol) was dissolved in dry acetone (40 mL) and stirred with 4 Å sieves. Concentrated HCl 38% (5 drops) was added and the reaction left to stir at room temperature. After 24 hours the crude reaction mixture was filtered, filter washed with additional dry acetone (10 mL) and the solvent reduced under vacuum to give **S18** (1.268 g) as a white solid that was used directly in the following reaction.

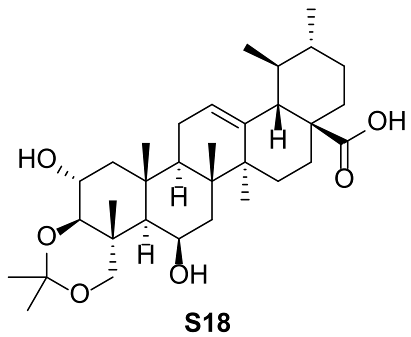


**S18**

**$^1$H NMR** (400 MHz, Methanol-*d*$_4$) δ 5.27 (t, *J* = 3.6 Hz, 1H), 4.14 – 4.07 (m, 1H), 3.75 (td, *J* = 10.4, 4.5 Hz, 1H), 3.65 (d, *J* = 10.5 Hz, 1H), 3.58 (d, *J* = 10.3 Hz, 1H), 3.33 (d, *J* = 8.7 Hz, 1H), 2.26 – 0.86 (m, 43H). **$^{13}$C NMR** (101 MHz, Methanol-*d*$_4$) δ 181.6, 139.0, 126.8, 100.6, 83.2, 73.6, 67.5, 66.5, 54.4, 53.1, 50.9, 49.3, 43.7, 41.3, 40.5, 40.4, 40.1, 38.9, 38.4, 38.1, 31.8, 30.1, 29.2, 25.3, 24.2, 24.2, 21.6, 19.7, 19.6, 18.9, 17.6, 15.4. **HREI-MS**: *m/z* calculated for $C_{33}H_{52}O_6Na$ [M+Na$^+$] 567.3656; observed 567.3661.

**Synthesis of 11**

**S18** (1268 mg, 2.38 mmol) and $K_2CO_3$ (997 mg, 7.14 mmol) were dissolved in dry DMF (25 mL). Allyl Bromide (0.412 mL, 4.76 mmol) was added under a nitrogen atmosphere. The mixture was stirred at room temperature overnight under nitrogen. After 22 hours, the mixture was diluted with EtOAc (100 mL) and extracted with $H_2O$ (3 × 100 mL) and brine (1 × 100 mL) The organic layer was dried over anhydrous magnesium sulphate. Filtration and evaporation of solvent at reduced pressure gave a light-yellow solid. Crude product was purified by column chromatography, (25%:75%, EtOAc:Hexane) to yield the product **11** (769 mg, 55.2%, over 2 steps)) as a white solid.

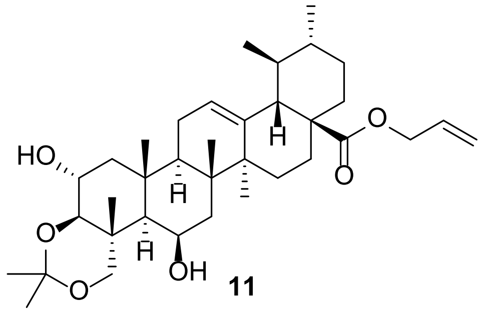


**11**

**$^1$H NMR** (400 MHz, Methanol-*d*$_4$) δ 5.92 (ddt, *J* = 17.2, 10.8, 5.6 Hz, 1H), 5.36 – 5.27 (m, 2H), 5.21 (dq, *J* = 10.5, 1.4 Hz, 1H), 4.49 (dt, *J* = 5.7, 1.5 Hz, 2H), 4.10 (dt, *J* = 4.9, 2.6 Hz,

1H), 3.76 (td, *J* = 10.5, 4.6 Hz, 1H), 3.65 (d, *J* = 10.4 Hz, 1H), 3.57 (d, *J* = 10.5 Hz, 1H), 3.32 (d, *J* = 8.1 Hz, 1H), 2.27 (dd, *J* = 11.3, 1.8 Hz, 1H), 2.14 – 0.84 (m, 42H). **$^{13}$C NMR** (101 MHz, Methanol-$d_4$) δ 178.7, 138.8, 133.7, 127.1, 118.3, 100.5, 83.2, 73.6, 67.4, 66.4, 66.1, 54.4, 53.1, 50.9, 49.5, 43.7, 41.3, 40.4, 40.3, 40.2, 38.8, 38.4, 38.0, 31.7, 30.1, 29.1, 25.3, 24.3, 24.2, 21.5, 19.7, 19.6, 18.9, 17.6, 15.4. **HREI-MS**: *m/z* calculated for $C_{36}H_{56}O_6Na$ [M+Na$^+$] 607.3969; observed 607.3977.

**Synthesis of 13**

Potassium *tert*-Butoxide (167.2 mg, 1.50 mmol), **11** (440.0 mg, 0.752 mmol), *tert*-Butyl 4-(bromomethyl)benzylcarbamate (331.9 mg, 1.13 mmol) and tetrabutylammonium iodide (26.9 mg, 0.0752 mmol) were dissolved in dry Toluene (25 mL) and were left to stir overnight at room temperature under nitrogen. After 24 hours, the reaction was transferred to a separating funnel, diluted with EtOAc (50 mL) and extracted with $H_2O$ (3 × 50 mL) and brine (1 × 50 mL). The organic phase was dried over anhydrous magnesium sulphate, filtered and solvent removed under reduced pressure. Crude product was purified by isocratic column chromatography (20%:80%, EtOAc:Hexane) giving crude product (520 mg) mixed with starting material **11**, The mixture was used crude directly in the following reaction with no further purification by dissolving it in MeOH (15 mL) under nitrogen before 5 M $HCl_{(aq)}$ (5.0 mL, 25 mmol) was injected into the reaction. It was left to stir overnight at room temperature under nitrogen. After 23 hours, the reaction was diluted with EtOAc (50 mL) and washed with 1 M $NaOH_{(aq)}$ (3 × 30 mL). The organic phase was dried over anhydrous magnesium sulphate, filtered and solvent removed under reduced pressure, yielding **12**, (242.1 mg, 48% (over 2 steps)). **12** was not found to be stable and was taken directly into the following reaction without NMR characterisation.

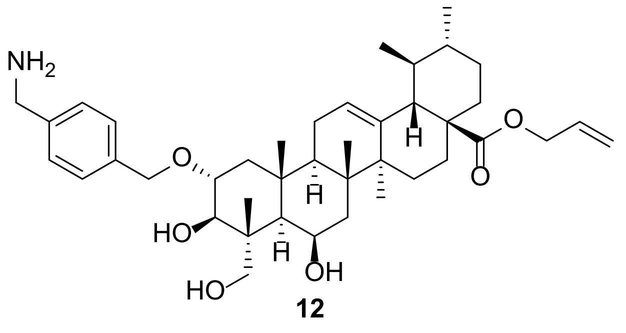


**12**

**HREI-MS**: *m/z* calculated for $C_{41}H_{62}NO_6$ [M+H$^+$] 664.4572; observed 664.4597.

**Synthesis of 13**

A solution of **12** (225.2 mg, 0.3395 mmol), **10** (63.2 mg, 0.1698 mmol), HOBt monohydrate (39.9 mg, 0.2546 mmol) and EDC.HCl (69.7 mg, 0.3395 mmol) in DMF (25 mL) was treated with $Et_3N$ (0.095 mL, 0.679 mmol) and stirred at room temperature overnight. After 18 hours the reaction was transferred to a separating funnel. 1 M $HCl_{(aq)}$ (40 mL) and brine (40 mL) were added. Aqueous layer was extracted with EtOAC (3 × 40 mL), organic layers combined, dried over anhydrous $MgSO_4$, filtered and reduced under vacuum. Crude product was purified by isocratic column chromatography (50%:50%, EtOAc:Hexane), followed by second isocratic column chromatography purification (100%, $Et_2O$) yielding **13** (57.8 mg, 34.8%).

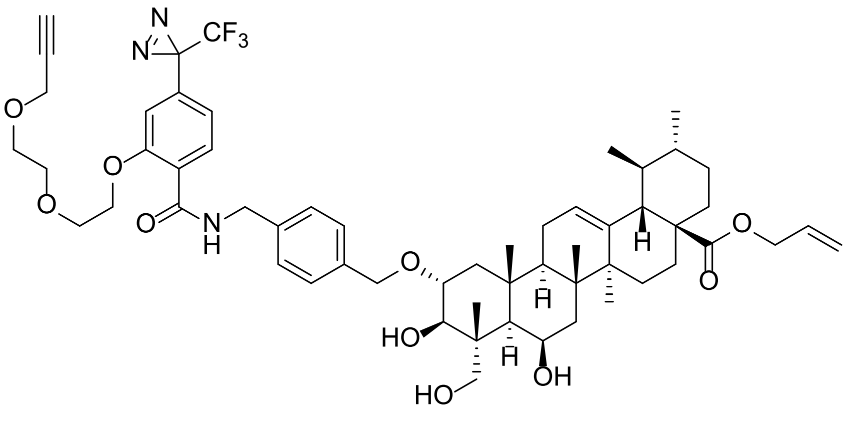

**13**

[1]**H NMR** (400 MHz, $CDCl_3$) δ 8.42 (t, *J* = 5.7 Hz, 1H), 8.20 (d, *J* = 8.3 Hz, 1H), 7.35 – 7.27 (m, 4H), 6.88 (ddd, *J* = 8.3, 1.8, 0.9 Hz, 1H), 6.66 (d, *J* = 1.8 Hz, 1H), 5.88 (ddt, *J* = 17.2, 10.4, 5.6 Hz, 1H), 5.33 – 5.25 (m, 2H), 5.18 (dq, *J* = 10.5, 1.3 Hz, 1H), 4.68 – 4.60 (m, 3H), 4.49 (ddt, *J* = 5.5, 2.7, 1.4 Hz, 2H), 4.45 – 4.35 (m, 2H), 4.24 – 4.18 (m, 2H), 4.05 (d, *J* = 2.4 Hz, 2H), 3.78 – 3.71 (m, 2H), 3.67 – 3.55 (m, 2H), 3.52 – 3.37 (m, 6H), 2.41 (t, *J* = 2.4 Hz, 1H), 2.25 (d, *J* = 11.3 Hz, 1H), 2.16 – 0.78 (m, 36H). [13]**C NMR** (101 MHz, $CDCl_3$) δ 177.2, 164.4, 156.8, 138.4, 137.6, 137.6, 133.6, 133.1, 132.6, 128.2, 127.8, 125.6, 123.3, 121.9 (q, *J* = 275.1 Hz), 119.6, 117.9, 110.7, 79.5, 77.1, 75.0, 70.8, 70.3, 69.0, 68.9, 68.4, 68.1, 67.7, 65.0, 58.4, 53.0, 48.5, 48.2, 47.7, 45.0, 43.7, 43.2, 42.8, 40.7, 39.2, 38.9, 38.8, 37.6, 36.8, 30.8, 28.5, 28.4 (q, *J* = 40.7 Hz), 28.0, 24.3, 23.8, 23.5, 21.2, 18.9, 18.6, 17.1, 14.6.

**Synthesis of 14**

To a solution of **13** (55.8 mg, 0.0638 mmol), $Pd(PPh_3)_4$ (23.6 mg, 0.0191 mmol) in THF (1 mL) was treated with pyrrolidine (0.026 mL, 0.319 mmol) and the reactions stirred at room temperature overnight. After 18 hours the reaction was transferred to a separating funnel, diluted with EtOAc (40 mL) and extracted with 1 M $HCl_{(aq)}$ (3 × 40 mL). Organic layer was collected, dried over anhydrous $MgSO_4$, filtered and reduced under vacuum. Crude product was purified by gradient column chromatography (50-100%, EtOAc:Hexane). Semi-Pure product was then further purified by isocratic column chromatography (5%:95%, MeOH:$CH_2Cl_2$) to give a single pure fraction of product **14** (13.4 mg, 25.0% yield).

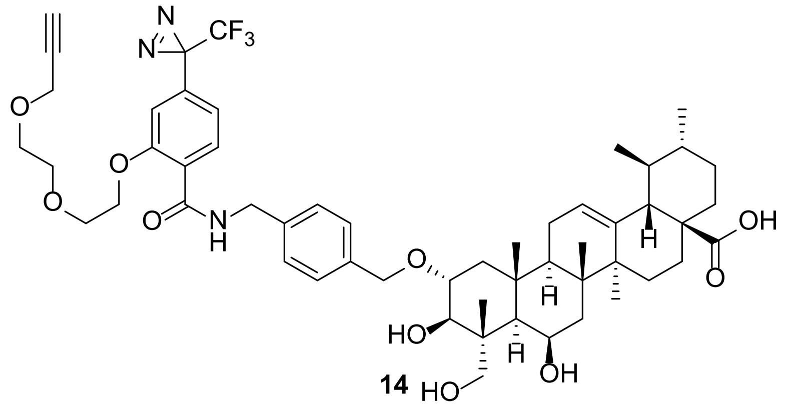


**14**

**$^{1}$H NMR** (400 MHz, $CDCl_3$) δ 8.46 (t, *J* = 5.8 Hz, 1H), 8.22 (d, *J* = 8.3 Hz, 1H), 7.35 (d, *J* = 8.2 Hz, 2H), 7.30 (d, *J* = 8.1 Hz, 2H), 6.90 (dd, *J* = 8.3, 1.5 Hz, 1H), 6.67 (d, *J* = 1.8 Hz, 1H), 5.24 (t, *J* = 3.7 Hz, 1H), 4.73 – 4.59 (m, 3H), 4.48 – 4.36 (m, 2H), 4.26 – 4.18 (m, 2H), 4.07 (d, *J* = 2.4 Hz, 2H), 3.80 – 3.68 (m, 3H), 3.65 – 3.56 (m, 1H), 3.55 – 3.37 (m, 6H), 2.42 (t, *J* = 2.4 Hz, 1H), 2.18 (d, *J* = 11.1 Hz, 1H), 2.13 – 0.74 (m, 36H). **$^{13}$C NMR** (101 MHz, $CDCl_3$) δ 183.0, 164.5, 156.9, 138.5, 137.5, 137.5, 133.7, 133.2, 128.4, 127.9, 125.6, 123.2, 122.0 (q, *J* = 274.9 Hz), 119.7, 110.7, 79.5, 77.4, 75.0, 71.0, 70.4, 69.1, 69.0, 68.4, 68.2, 68.1, 58.5, 52.6, 48.8, 48.0, 47.7, 44.9, 43.7, 43.2, 42.7, 40.4, 39.2, 38.9, 38.7, 37.6, 36.8, 30.8, 29.8, 28.5 (q, *J* = 40.8 Hz), 28.0, 24.3, 23.8, 23.4, 21.3, 19.0, 18.5, 17.1, 14.6.

**Synthesis of 15**

A solution of *p*-tolylmethanamine (0.034 mL, 0.264 mmol), **10** (65.5 mg, 0.176 mmol), HOBt monohydrate (40.8 mg, 0.264 mmol) and EDC.HCl (74.3 mg, 0.352 mmol) in DMF (10 mL) was treated with $Et_3N$ (0.049 mL, 0.352 mmol) and stirred at room temperature overnight. After 18 hours the reaction was transferred to a separating funnel. 1 M $HCl_{(aq)}$ (20 mL) and brine (20 mL) were added. The aqueous layer was extracted with EtOAC (3 × 20 mL), organic layers combined, dried over anhydrous $MgSO_4$, filtered and reduced under vacuum. Crude product was purified by isocratic column chromatography (25%:75%, EtOAc:hexane) to give **15** (70.0 mg, 83.7%).

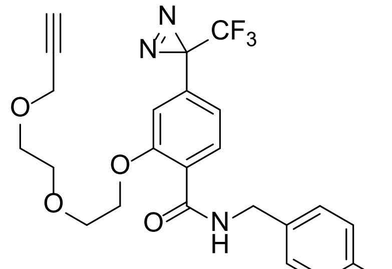


**15**

**$^{1}$H NMR** (400 MHz, $CDCl_3$) δ 8.34 (t, *J* = 5.6 Hz, 1H), 8.24 (d, *J* = 8.3 Hz, 1H), 7.25 (d, *J* = 7.9 Hz, 2H), 7.14 (d, *J* = 7.9 Hz, 2H), 6.91 – 6.87 (m, 1H), 6.66 (d, *J* = 1.7 Hz, 1H), 4.62 (d, *J* = 5.6 Hz, 2H), 4.24 – 4.17 (m, 2H), 4.07 (d, *J* = 2.4 Hz, 2H), 3.78 – 3.70 (m, 2H), 3.49 – 3.44 (m, 2H), 3.40 – 3.35 (m, 2H), 2.41 (t, *J* = 2.4 Hz, 1H), 2.33 (s, 3H). **$^{13}$C NMR** (101 MHz, $CDCl_3$) δ

164.2, 156.8, 137.0, 135.7, 133.6, 133.2, 129.4, 127.8, 123.3, 122.0 (q, *J* = 275.0 Hz), 119.6 (q, *J* = 0.7 Hz), 110.6 (q, *J* = 1.7 Hz), 79.5, 74.8, 70.4, 69.1, 69.0, 68.4, 58.5, 43.8, 28.5 (q, *J* = 40.6 Hz), 21.2. **HREI-MS**: *m/z* calculated for $C_{24}H_{24}F_3N_3O_4Na$ [M+Na$^+$] 498.1611; observed 498.1623.

#### [1]H NMR Spectra of Compound 2

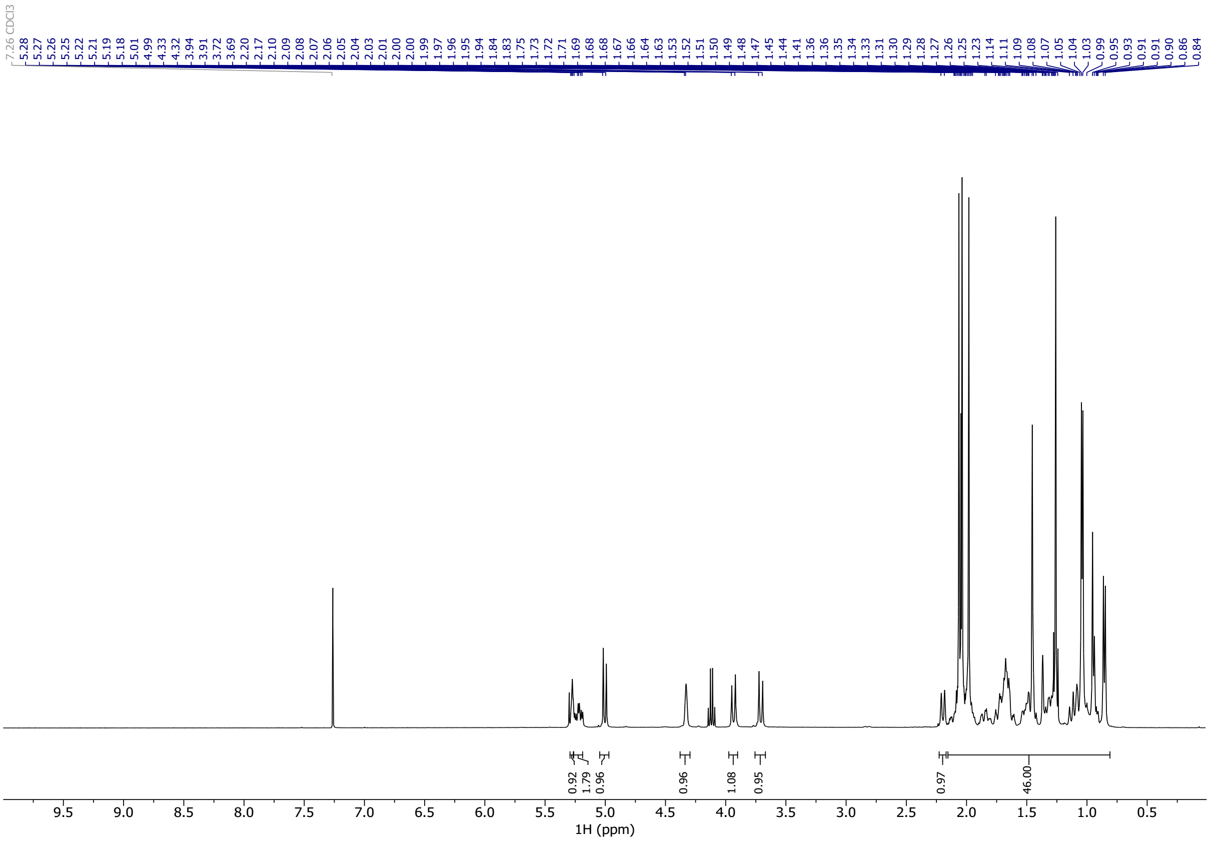

#### [13]C NMR Spectra of Compound 2

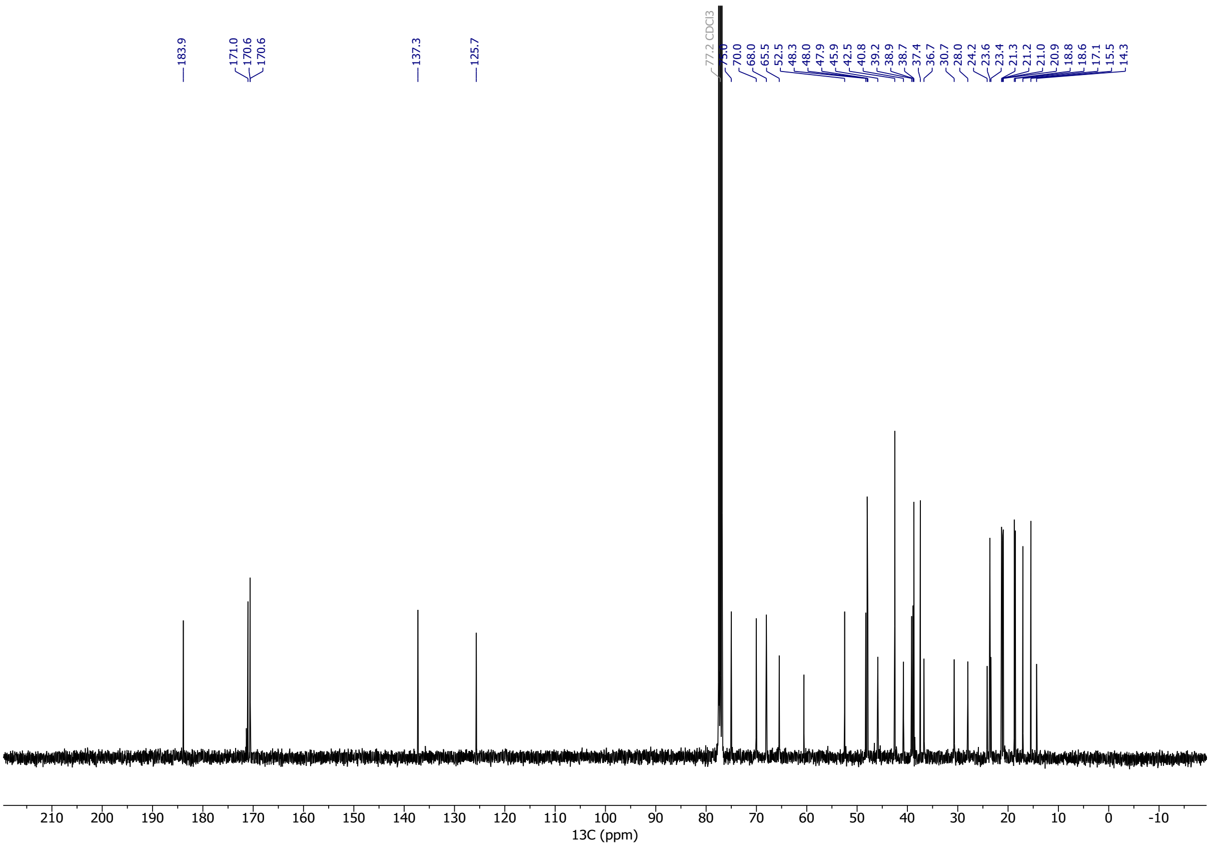

#### $^{1}$H NMR Spectra of Compound S1

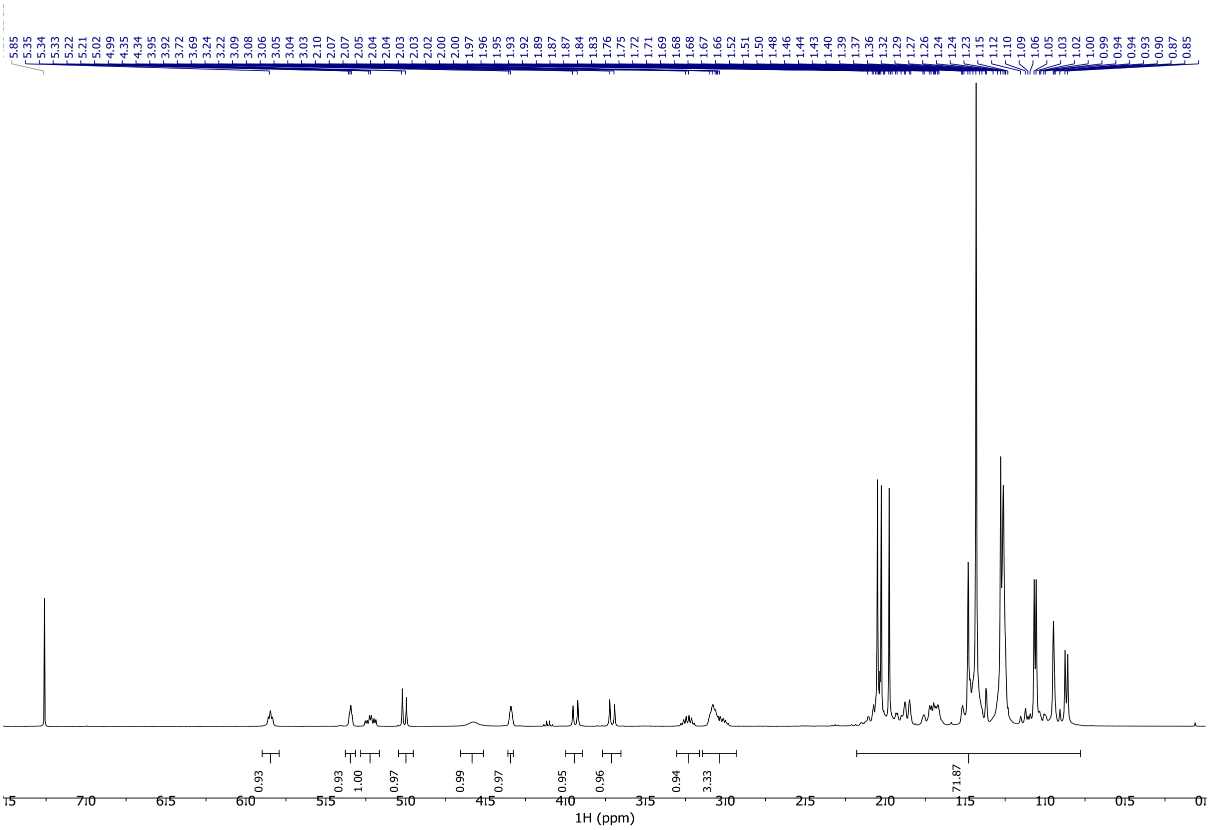


#### $^{13}$C NMR Spectra of Compound S1

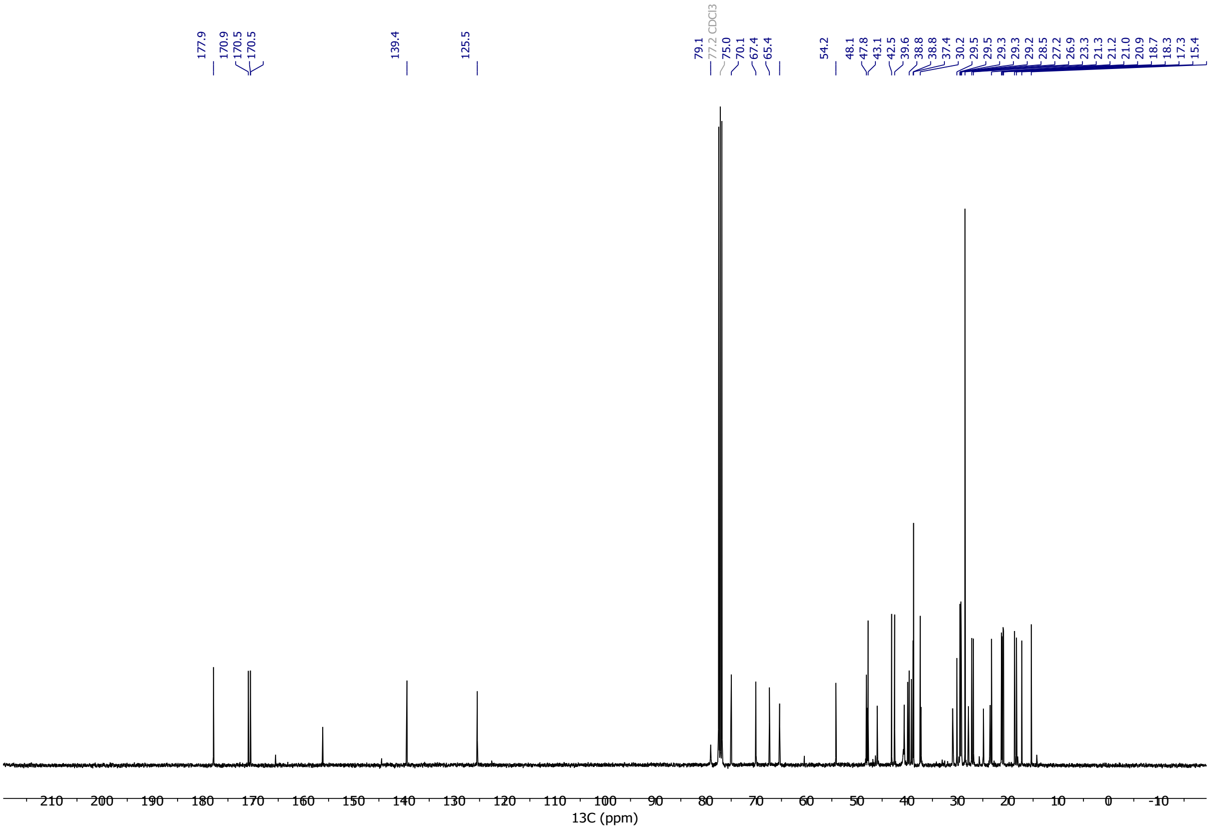

#### [1]H NMR Spectra of Compound 3

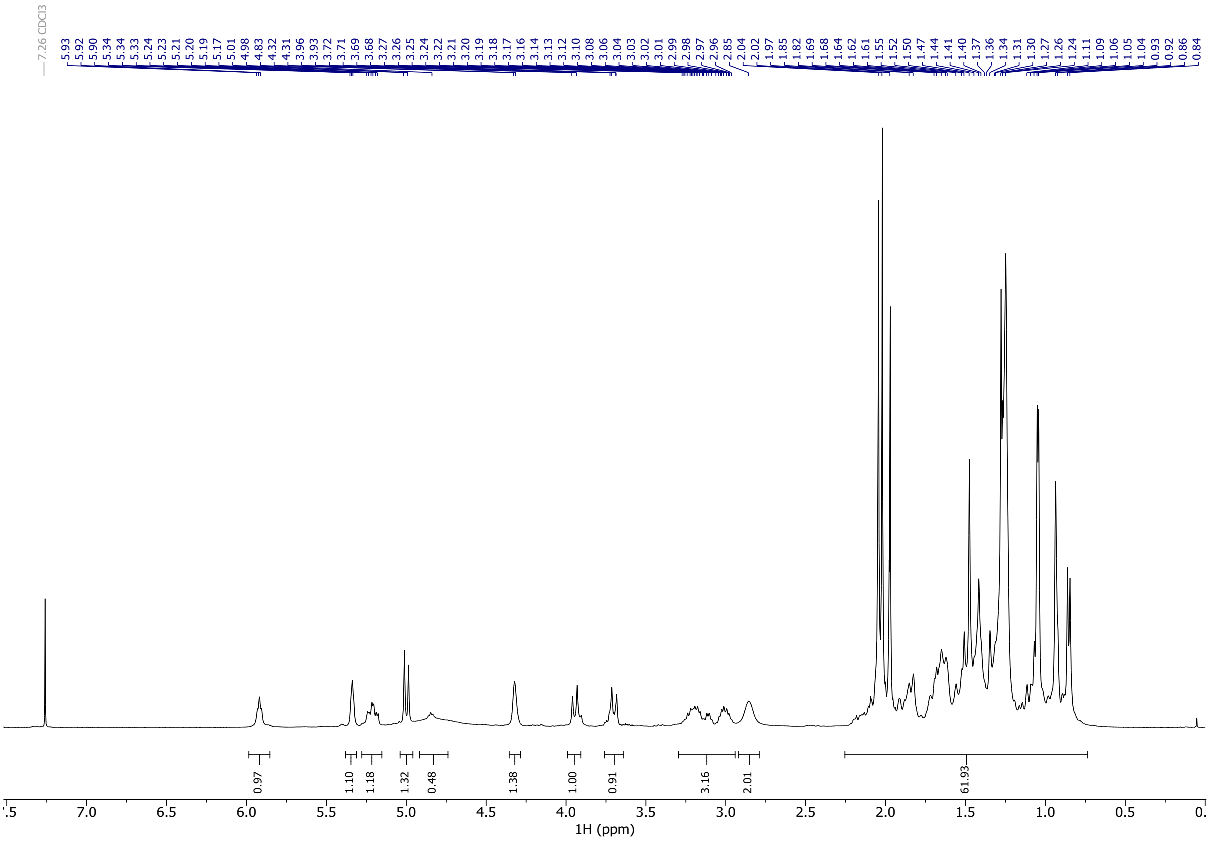

#### [13]C NMR Spectra of Compound 3

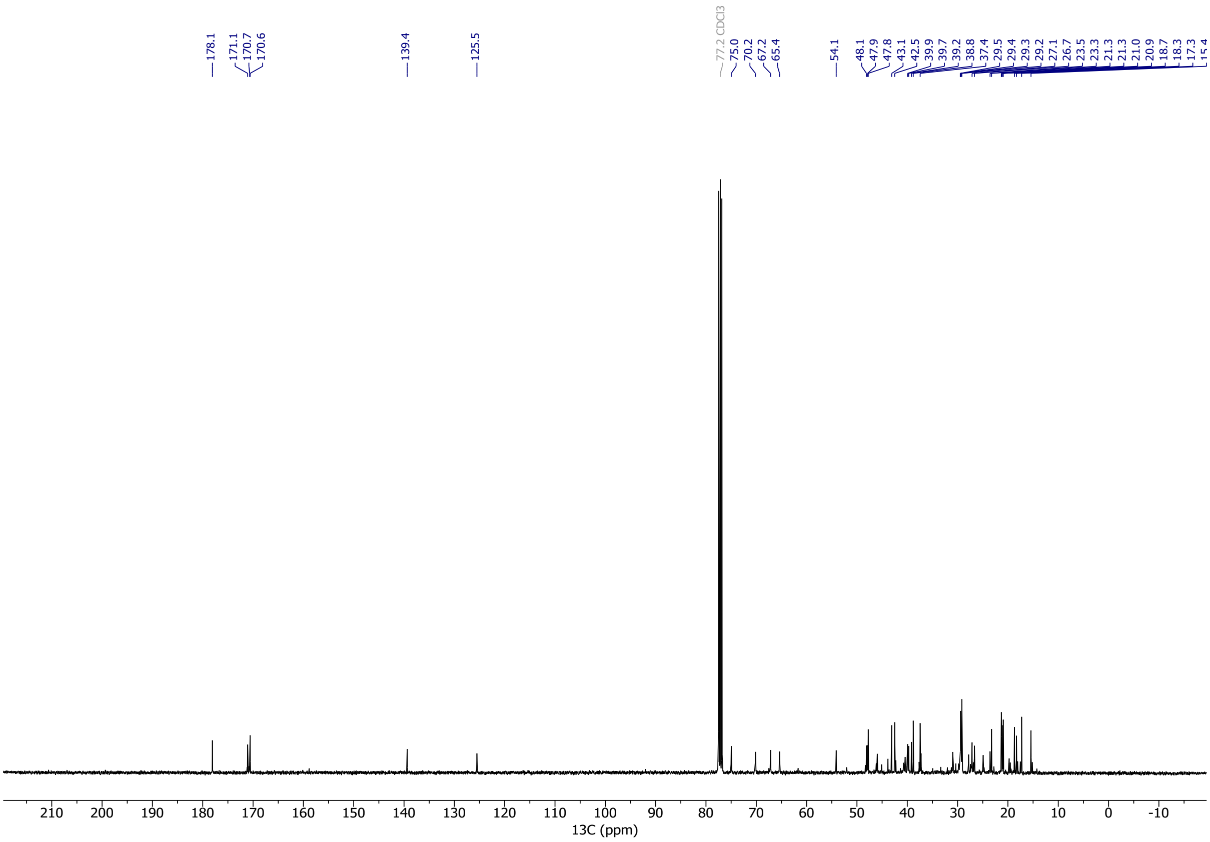

#### $^1$H NMR Spectra of Compound 4

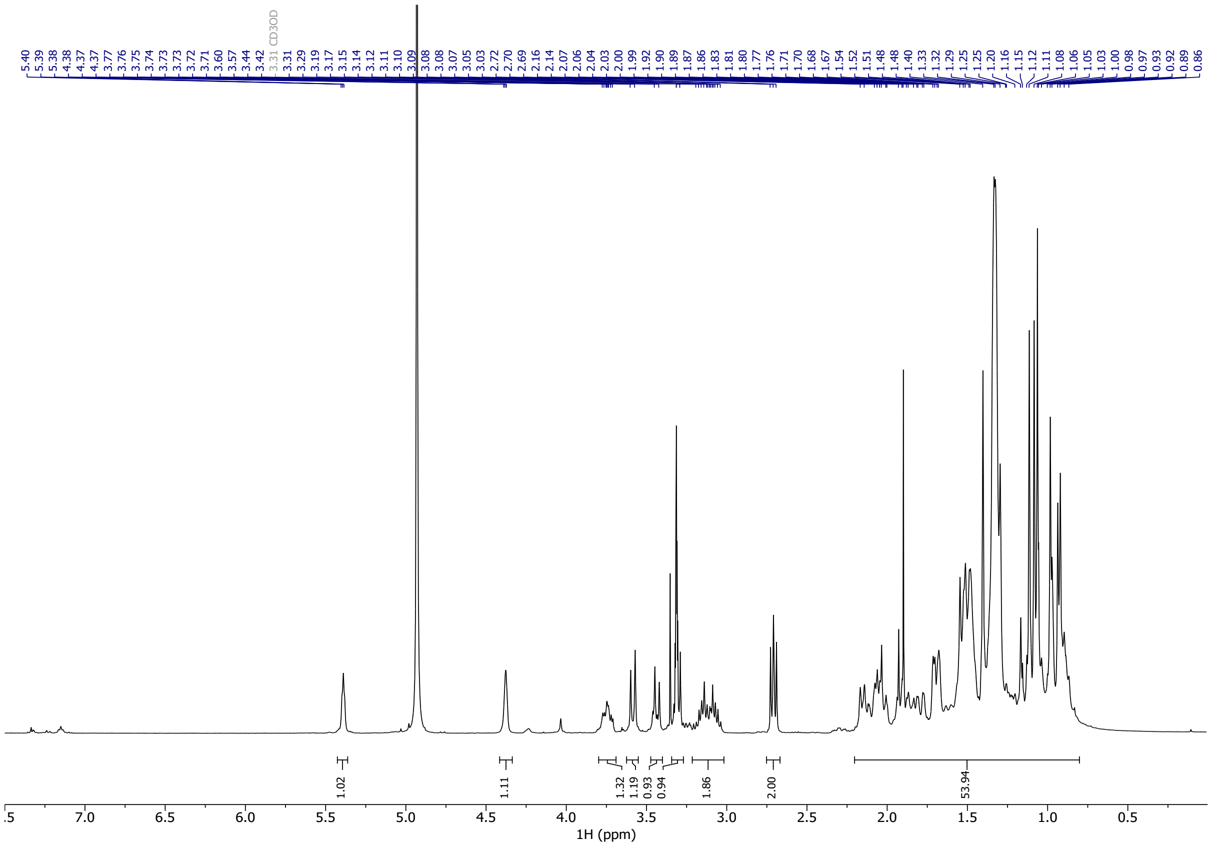

#### $^{13}$C NMR Spectra of Compound 4

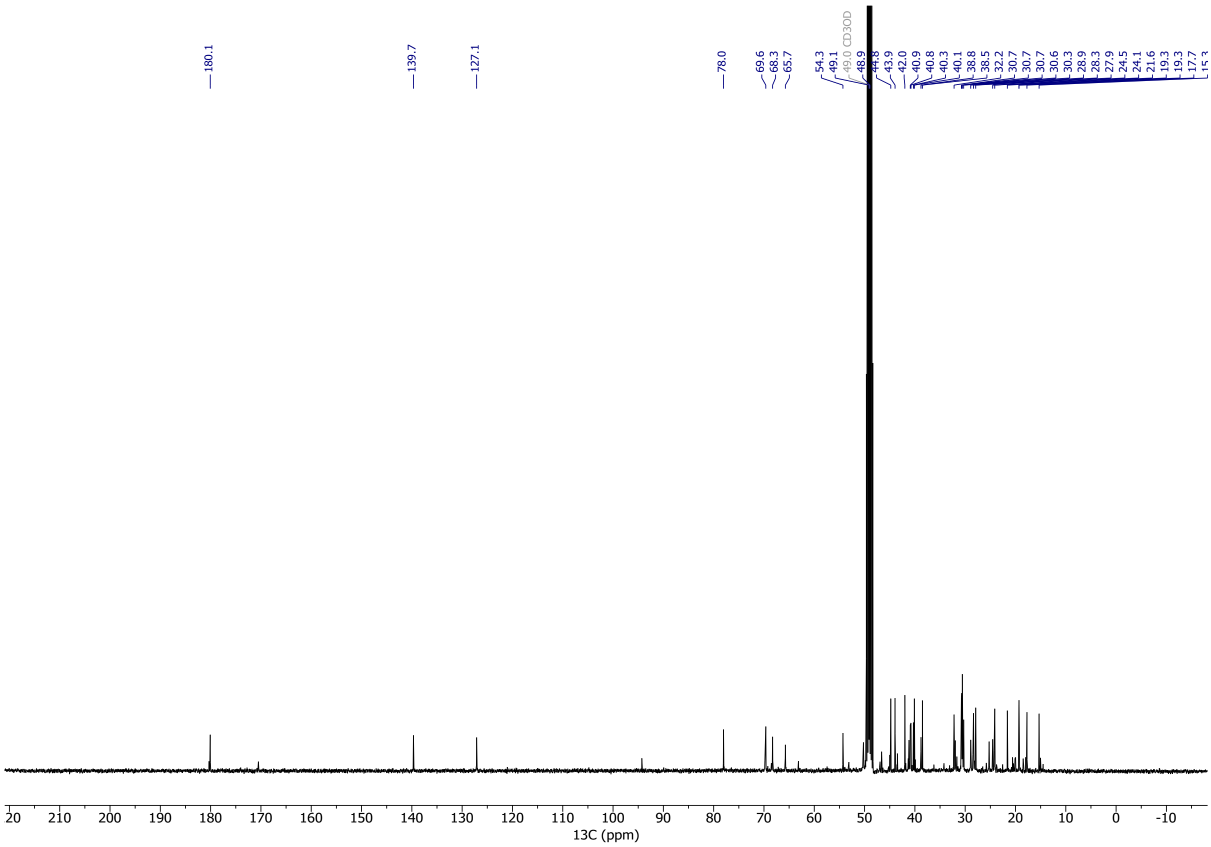

#### [1]H NMR Spectra of Compound 5

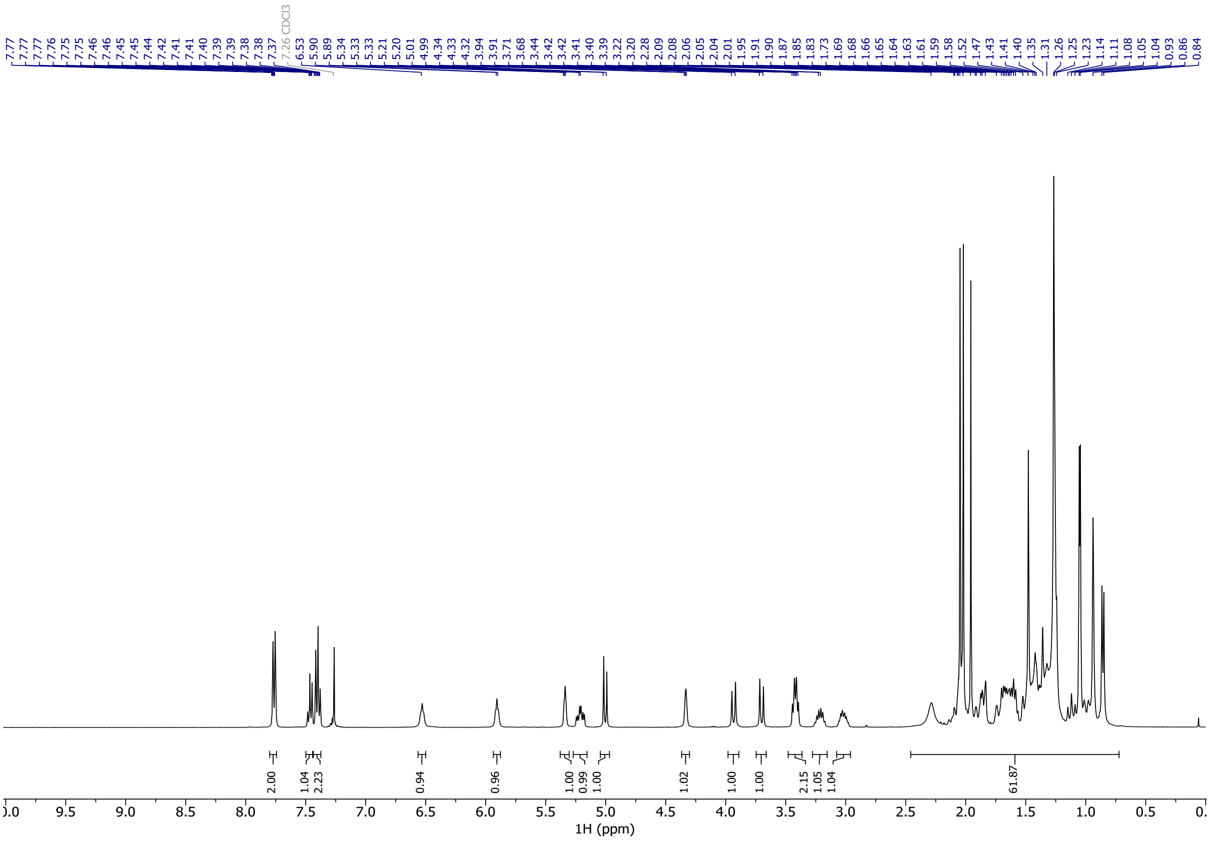


#### [13]C NMR Spectra of Compound 5

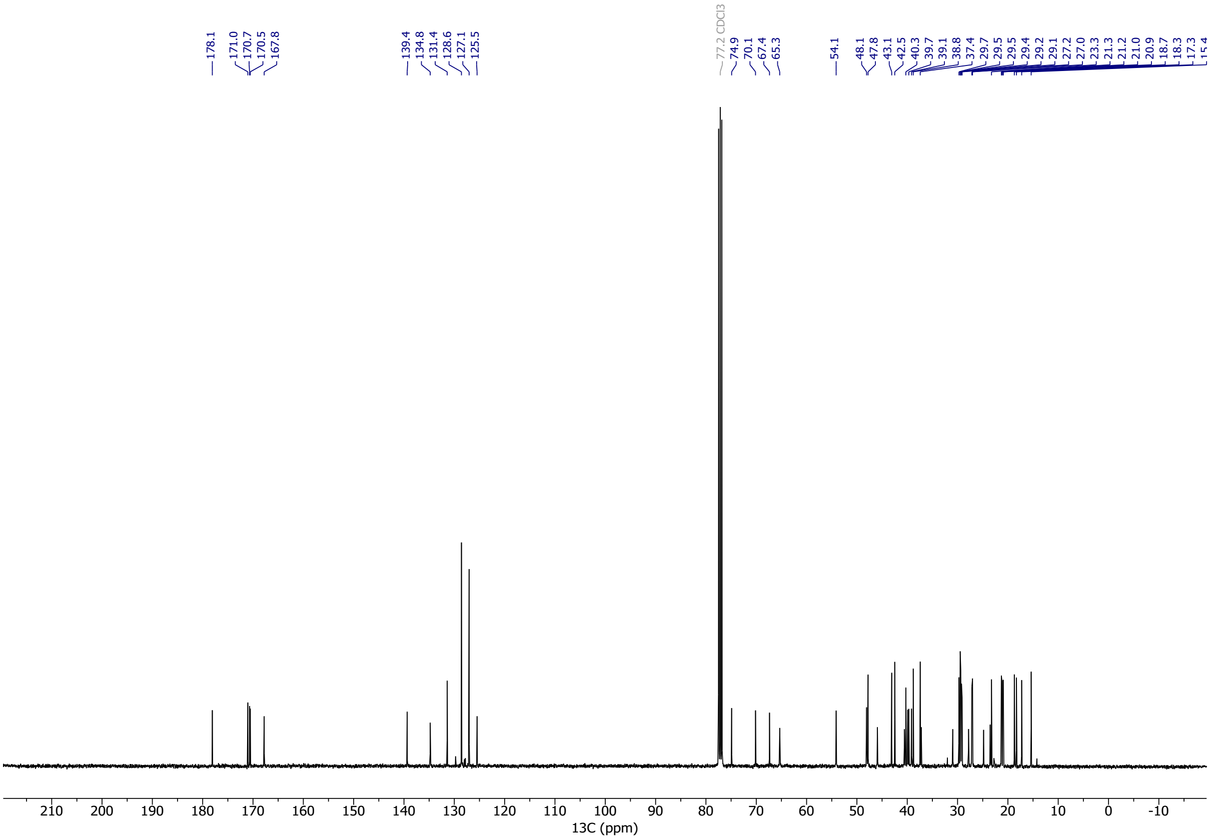

#### $^{1}$H NMR Spectra of Compound 6

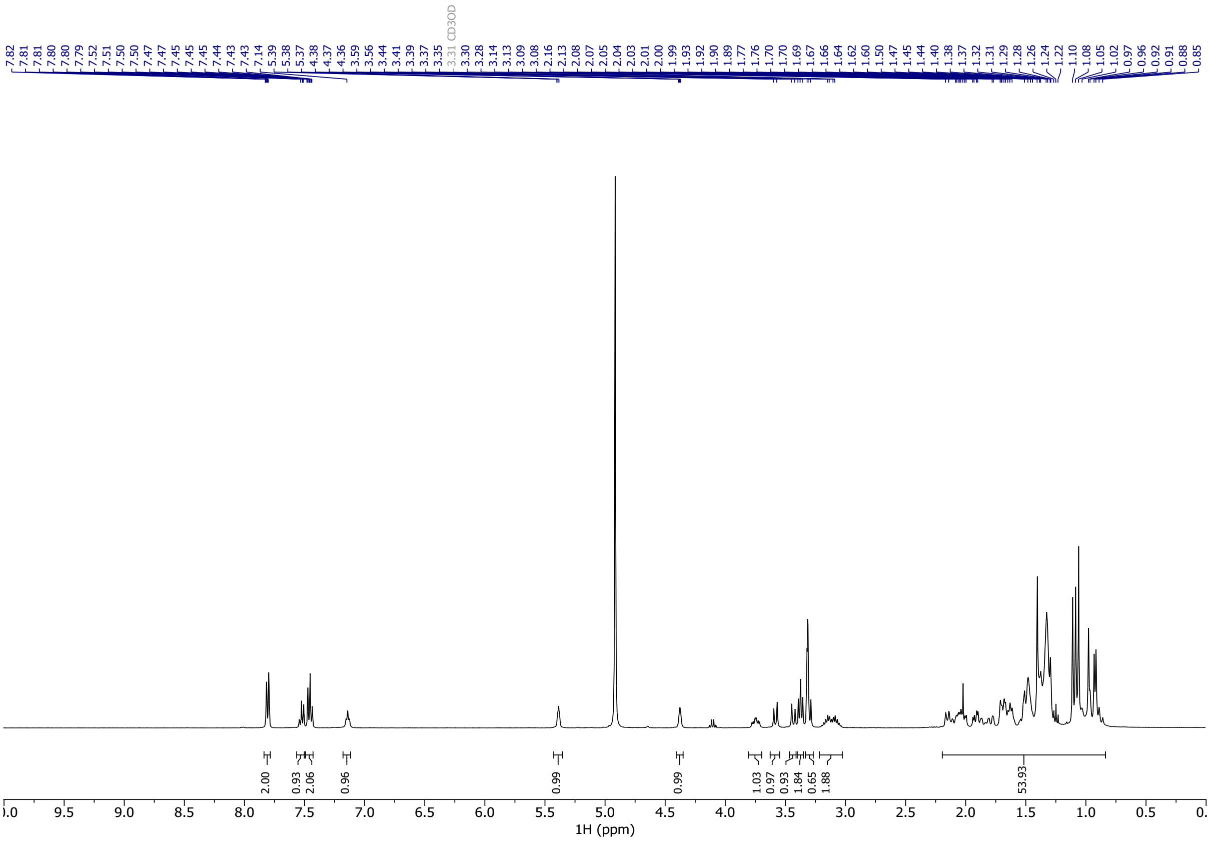


#### $^{13}$C NMR Spectra of Compound 6

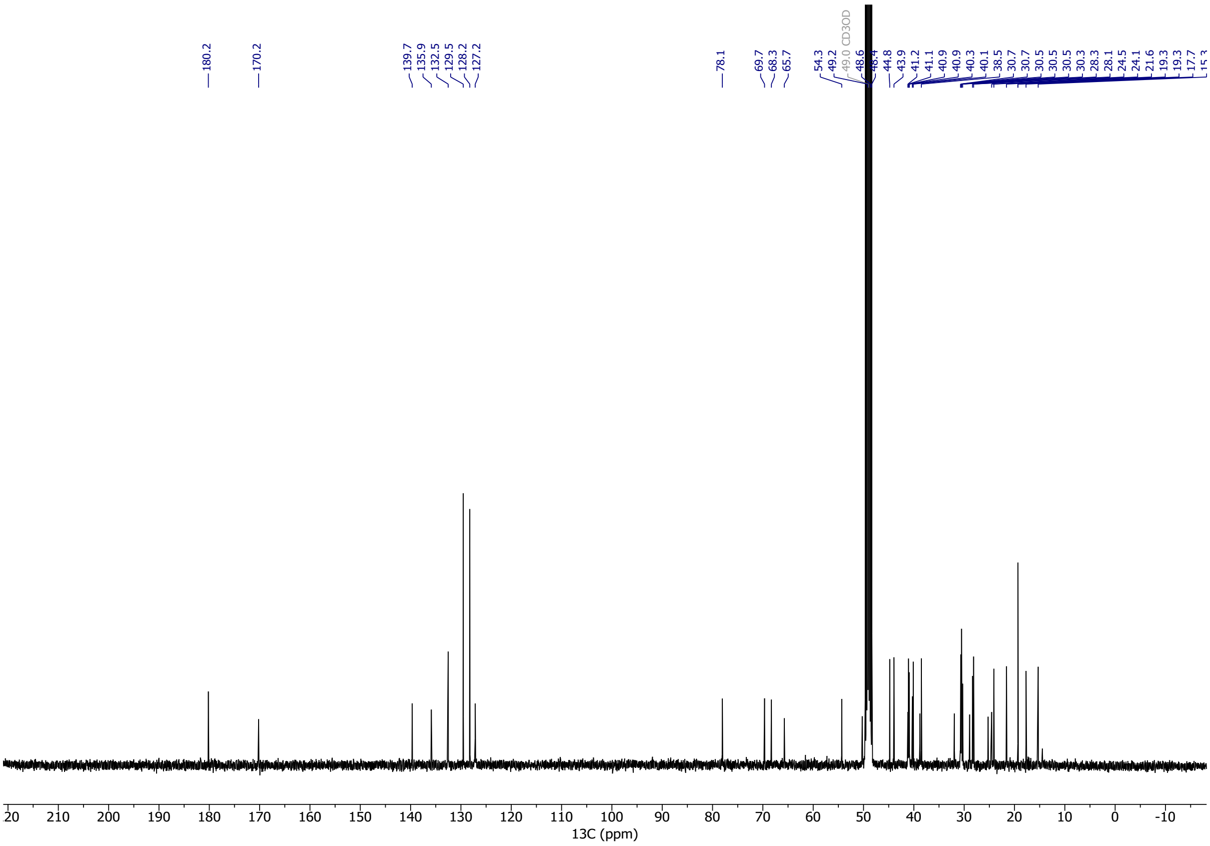

#### ¹H NMR Spectra of Compound S2

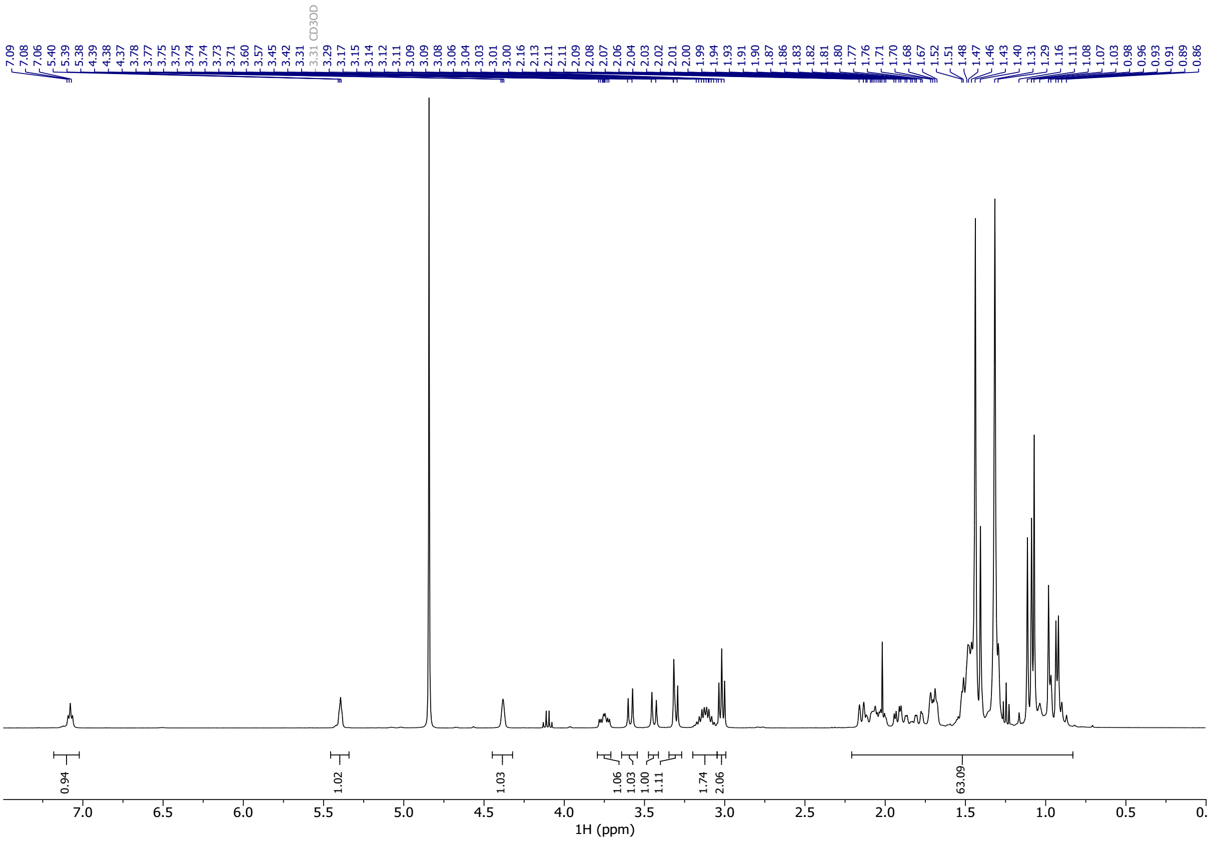


#### ¹³C NMR Spectra of Compound S2

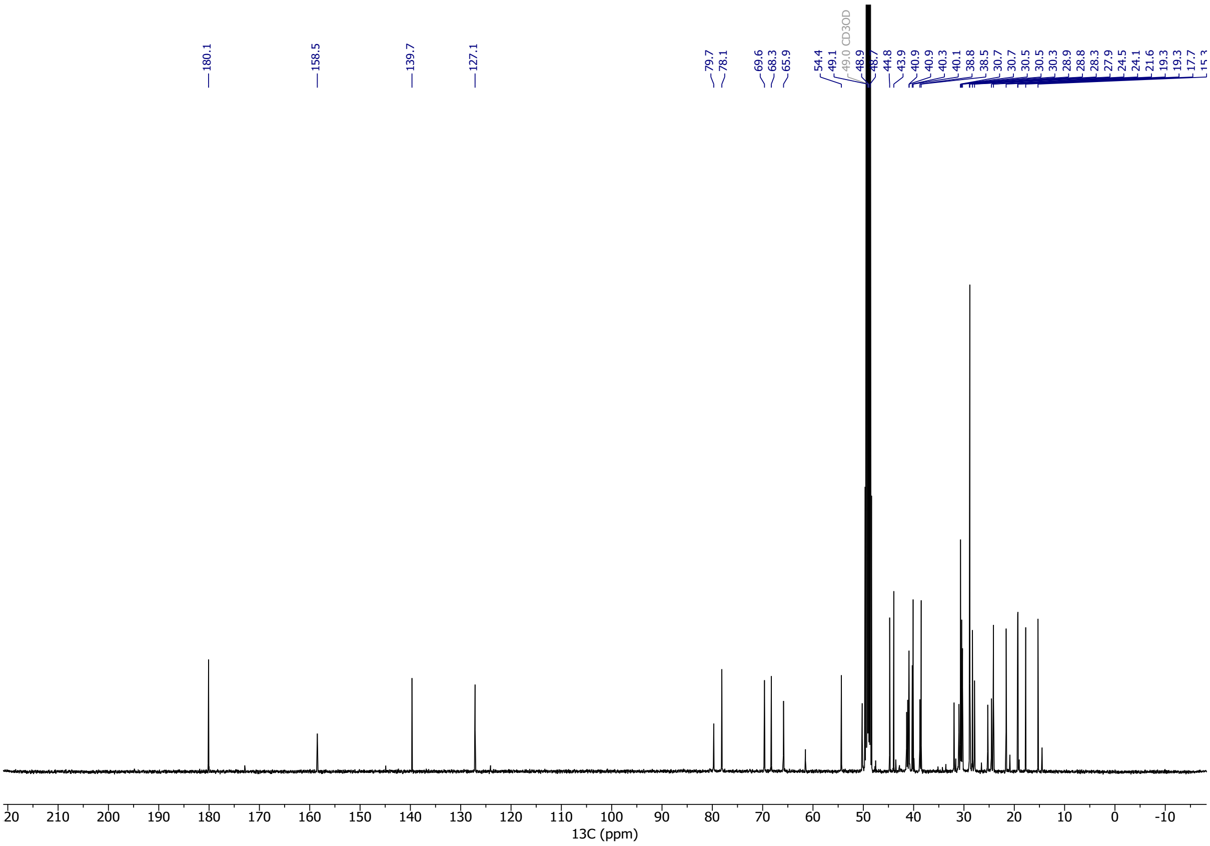

#### $^{1}$H NMR Spectra of Compound S3

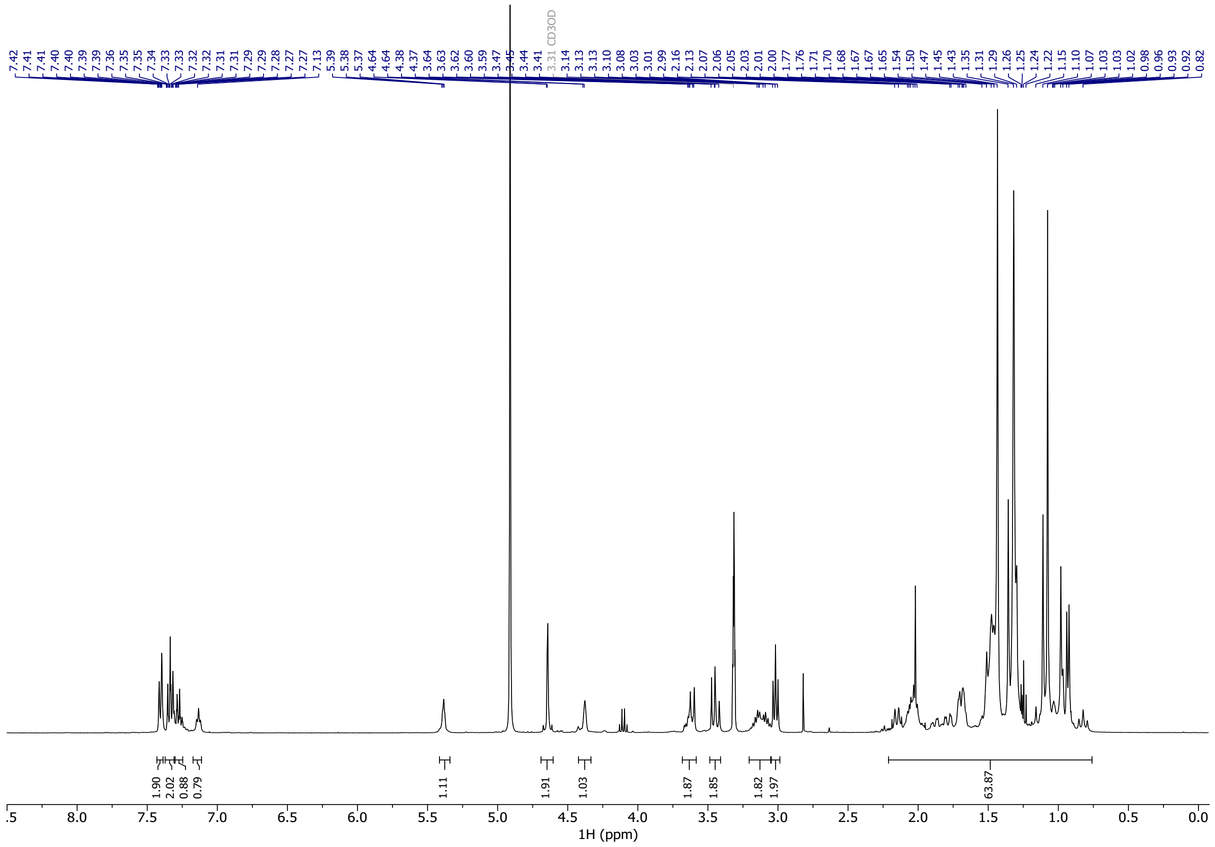

#### $^{13}$C NMR Spectra of Compound S3

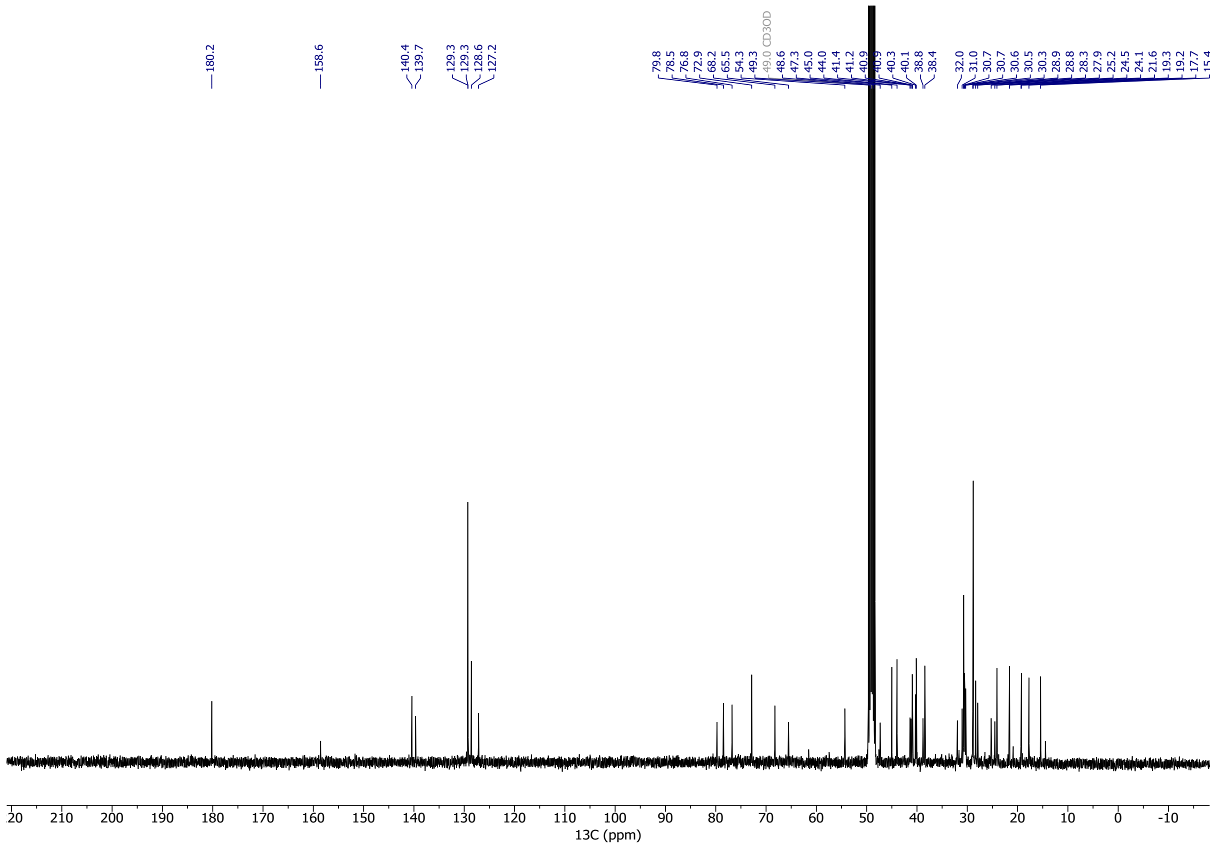

### $^{1}$H NMR Spectra of Compound S4

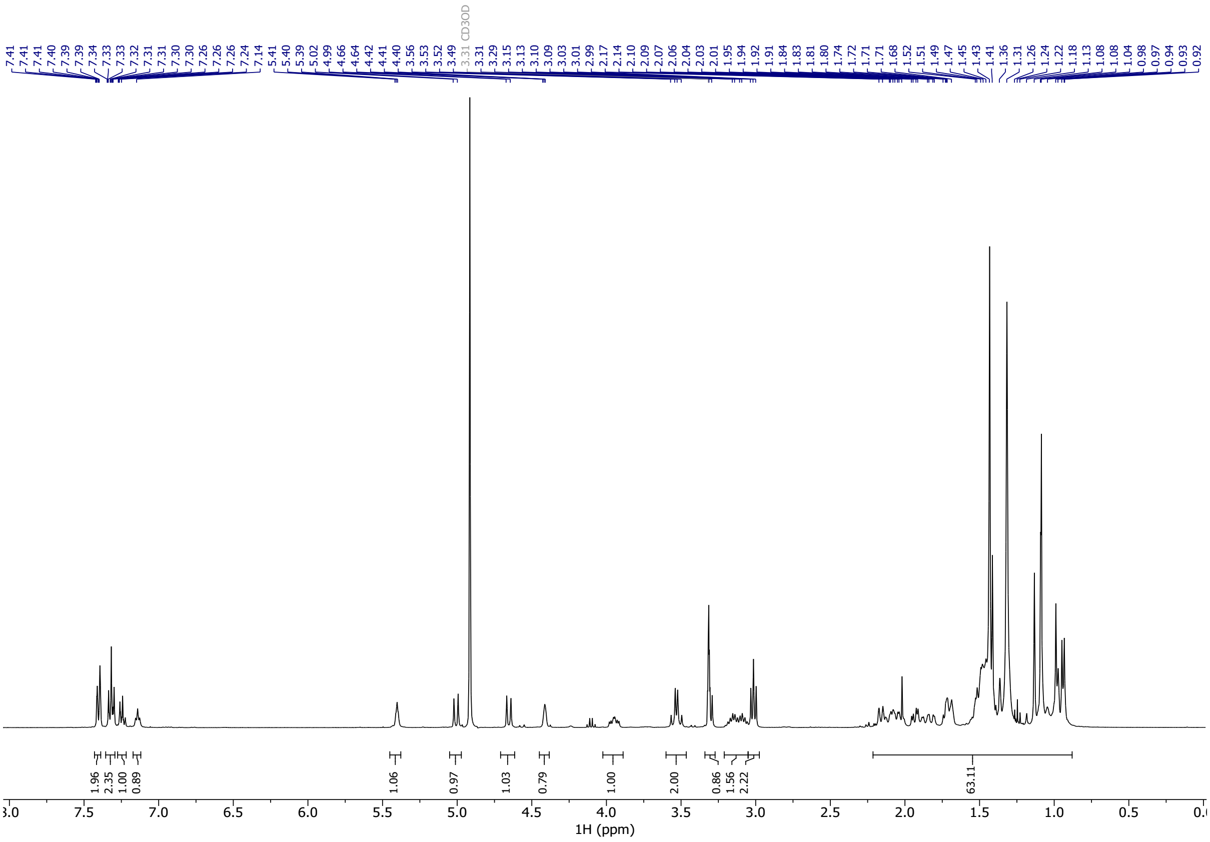

### $^{13}$C NMR Spectra of Compound S4

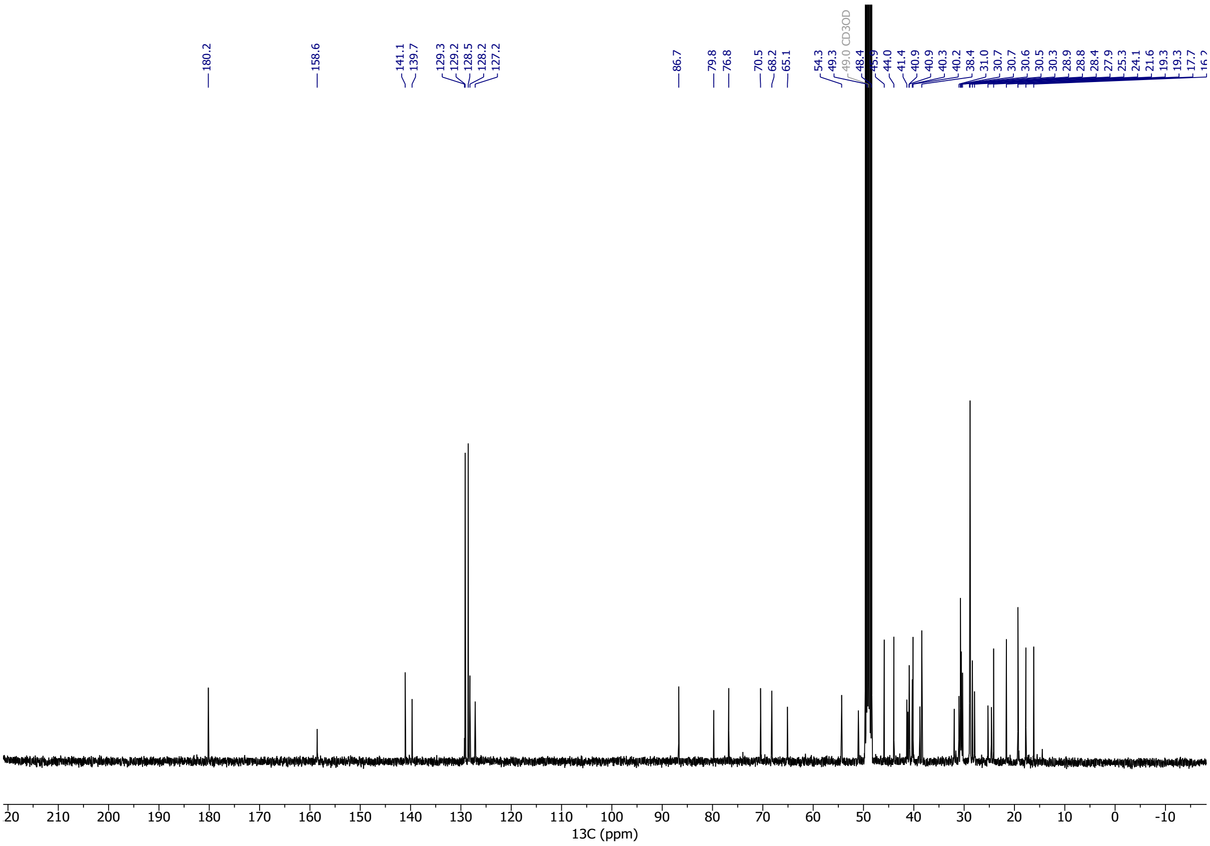

#### $^{1}$H NMR Spectra of Compound S5

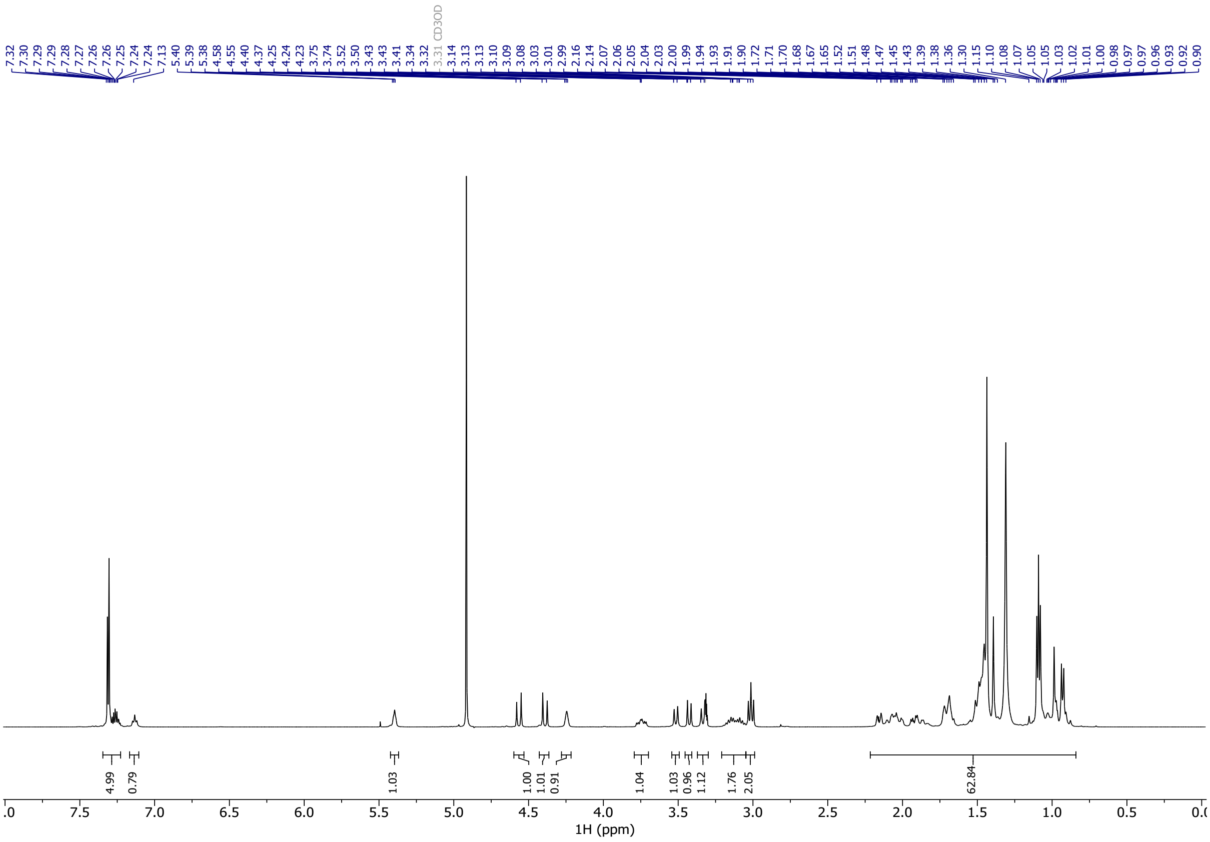

#### $^{13}$C NMR Spectra of Compound S5

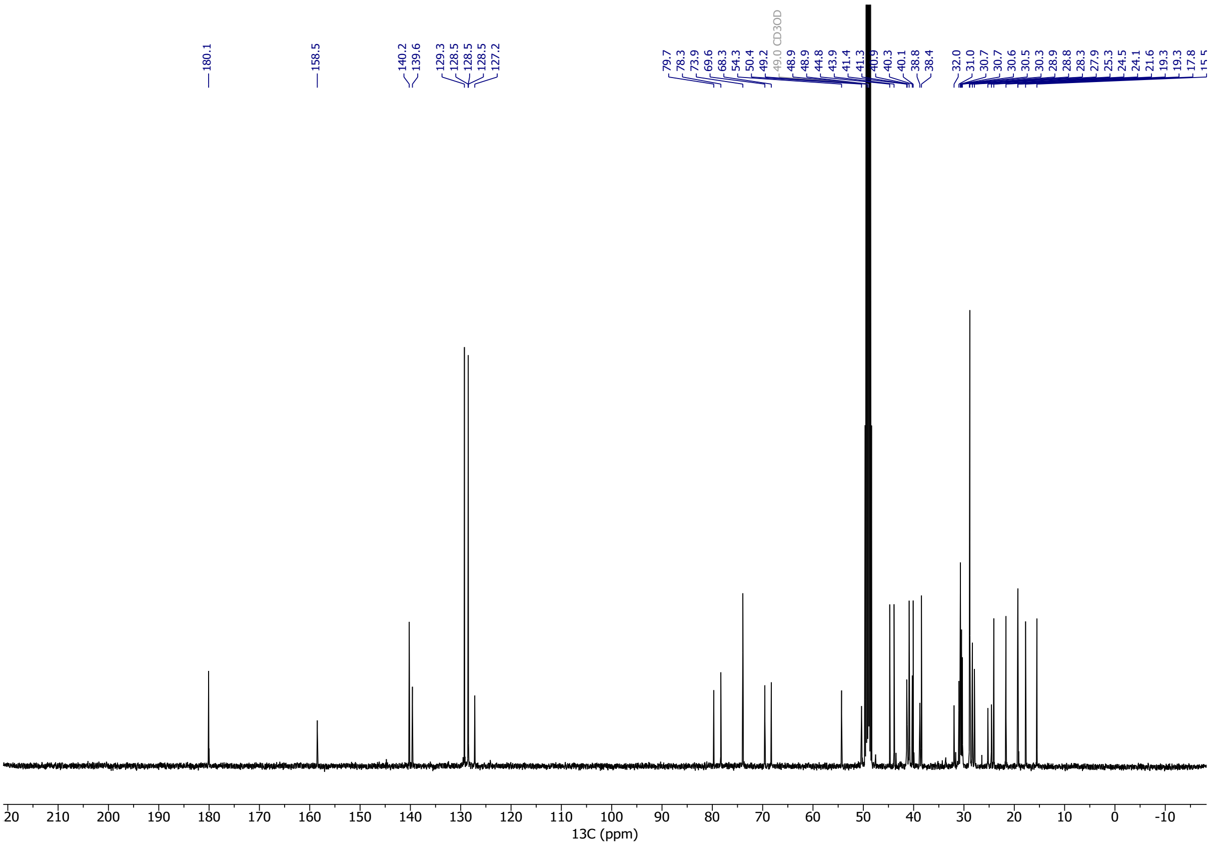

#### $^{1}$H NMR Spectra of Compound 7

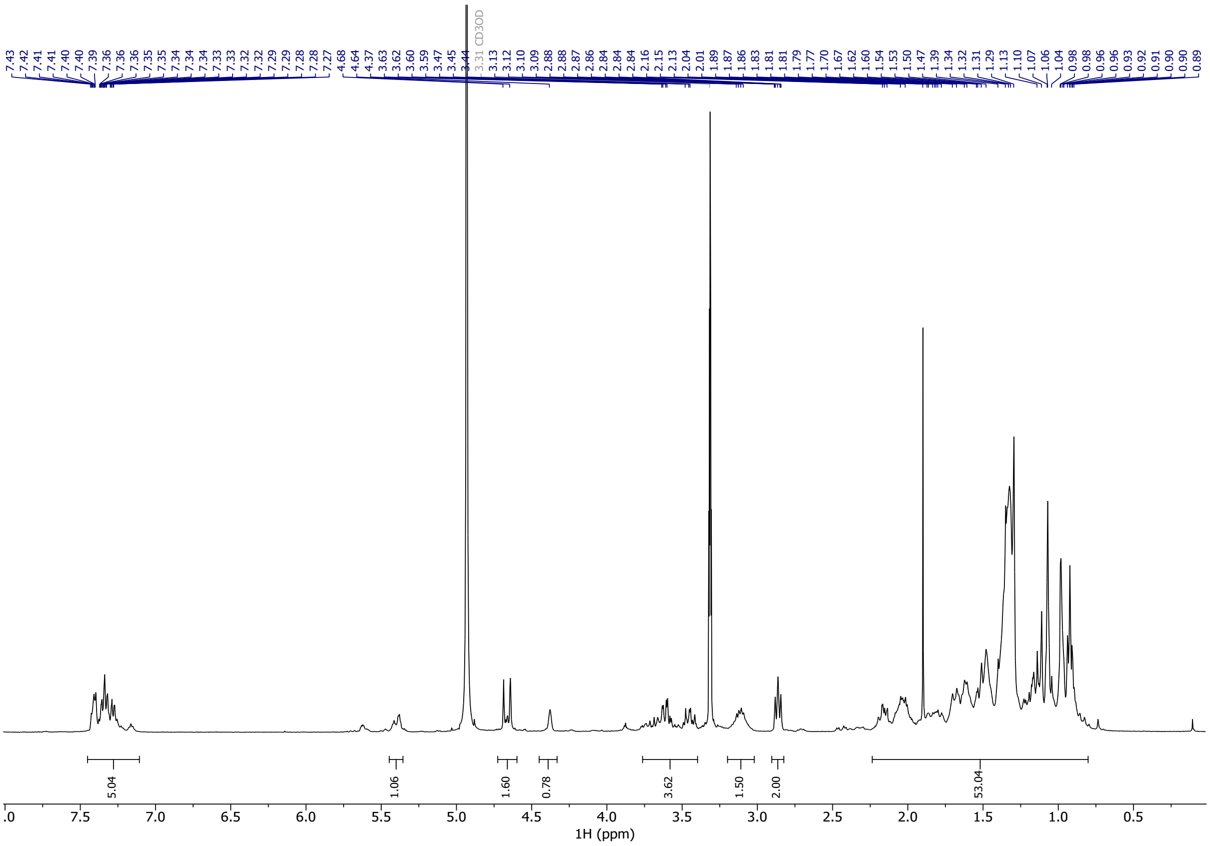

#### $^{13}$C NMR Spectra of Compound 7

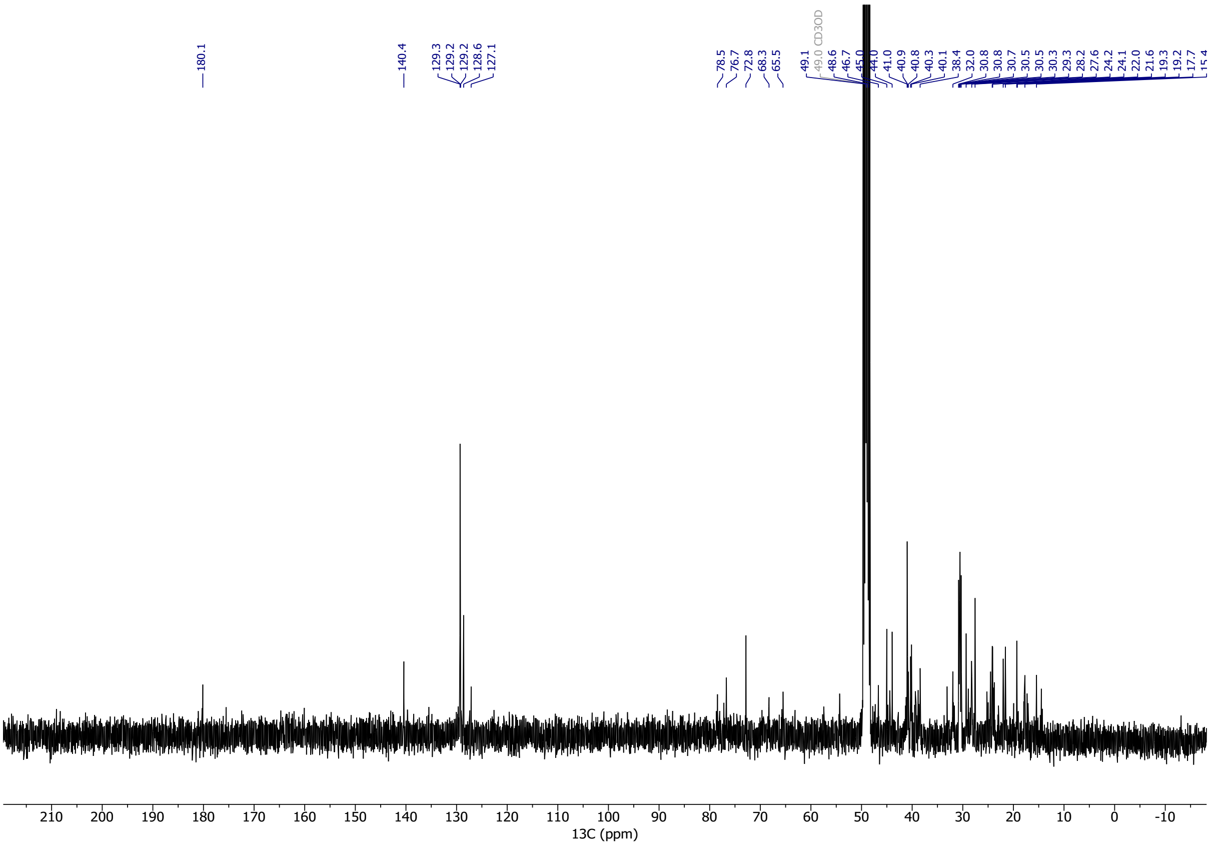

#### ¹H NMR Spectra of Compound 8

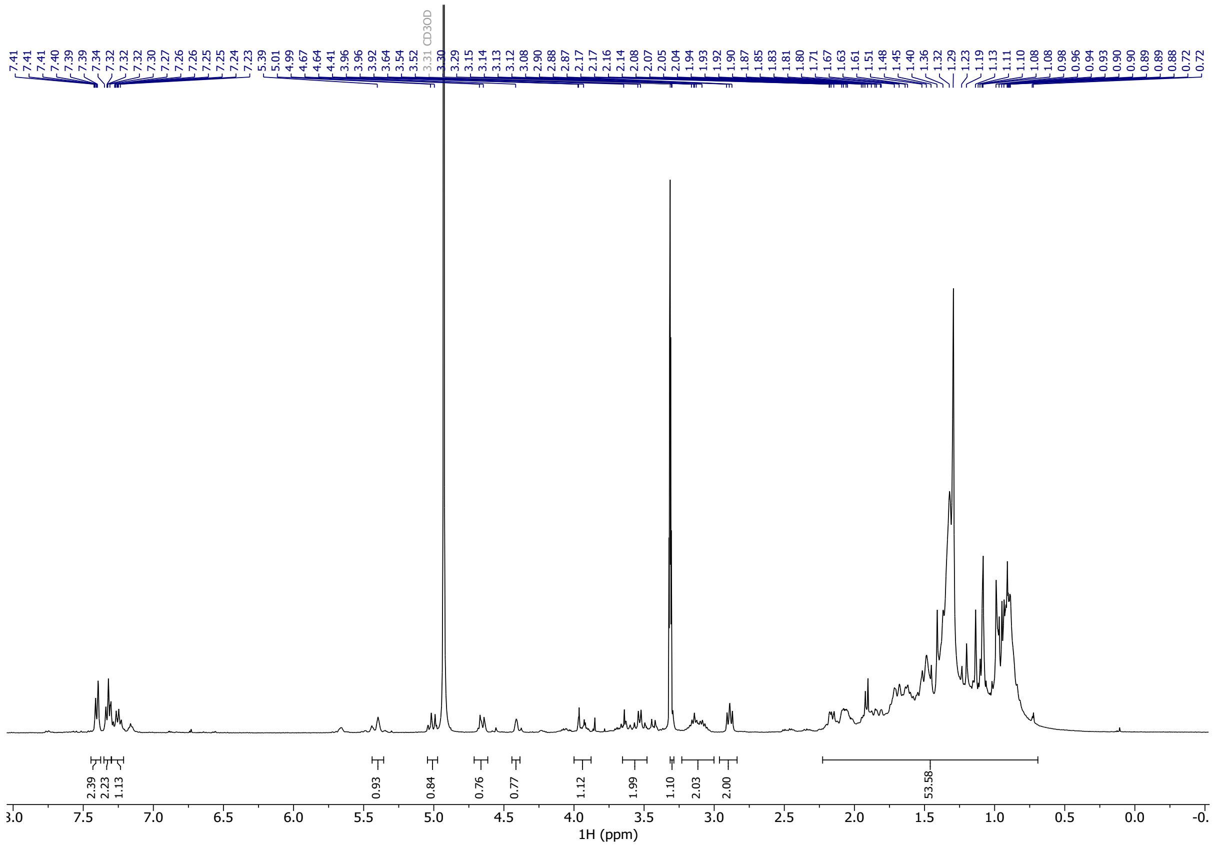

#### ¹³C NMR Spectra of Compound 8

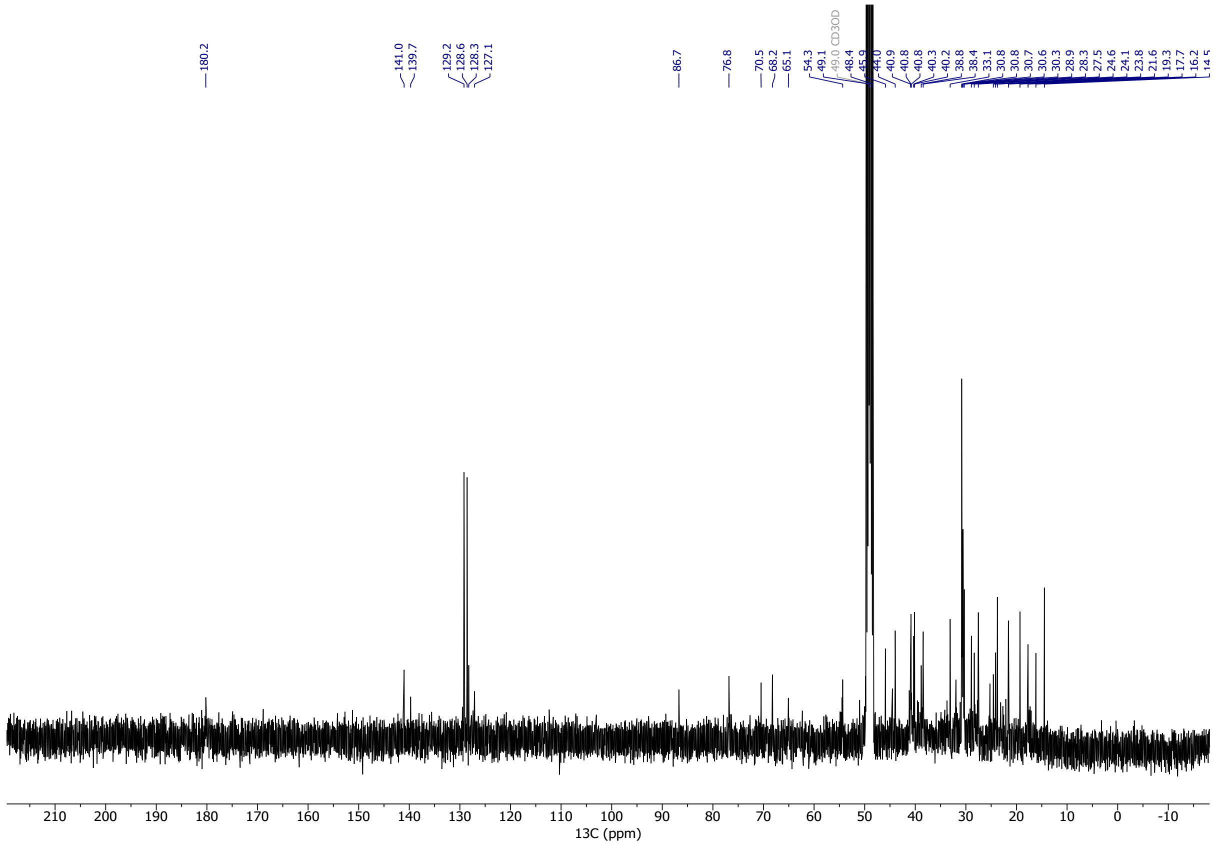

#### $^1$H NMR Spectra of Compound 9

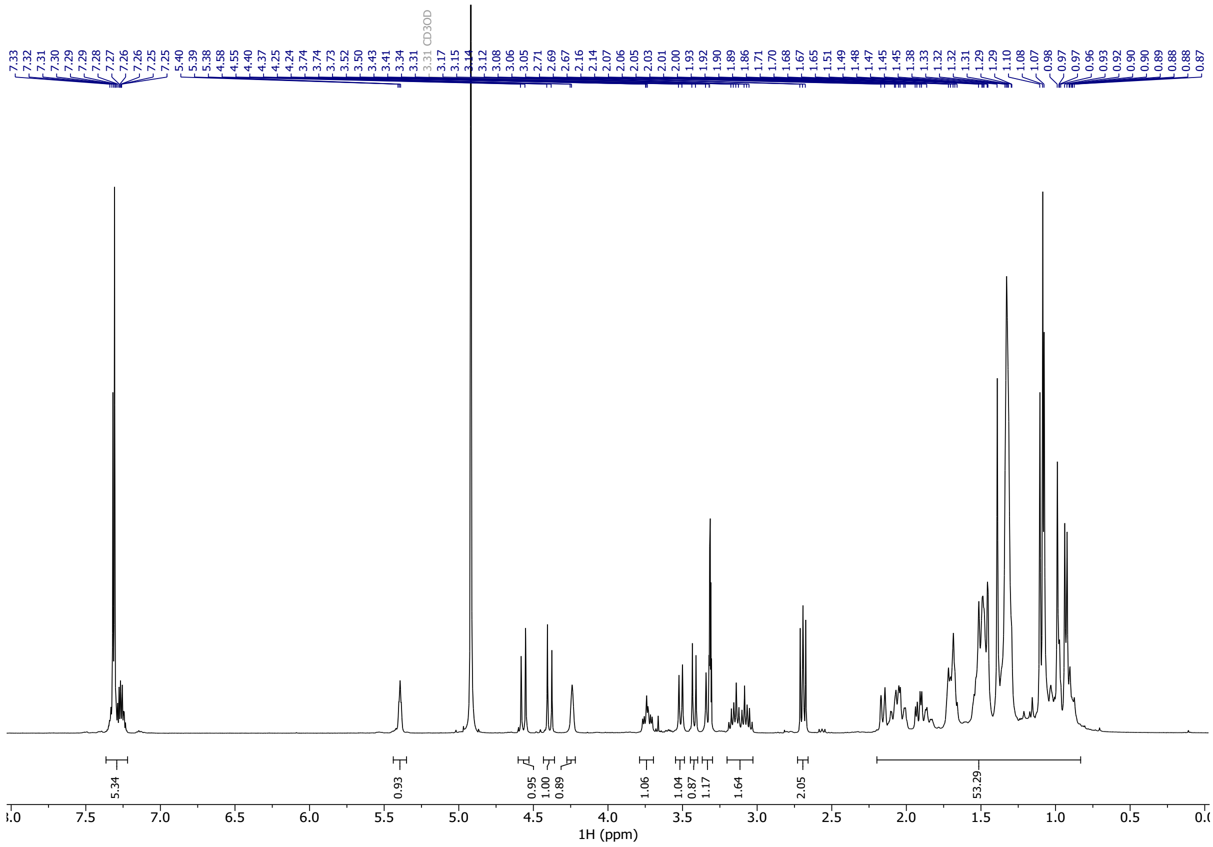

#### $^{13}$C NMR Spectra of Compound 9

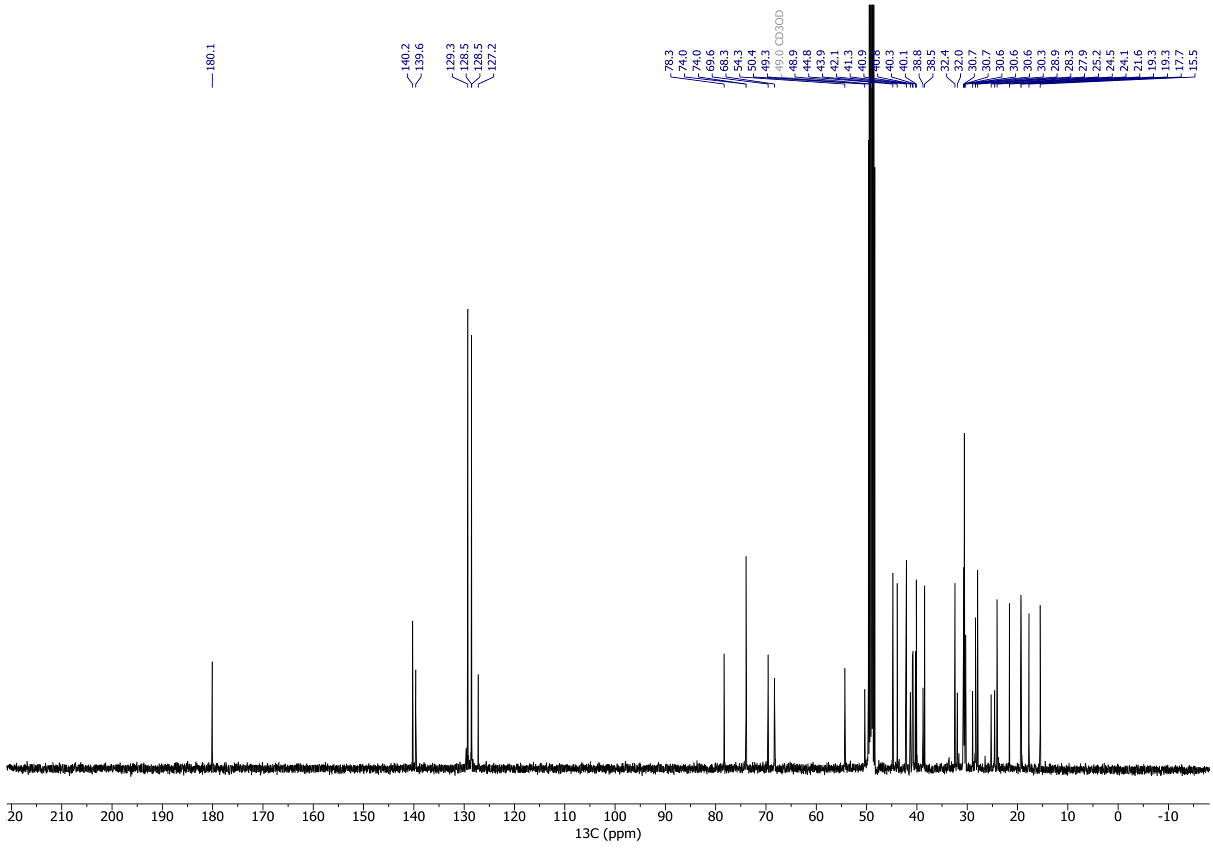

#### [1]H NMR Spectra of Compound S6

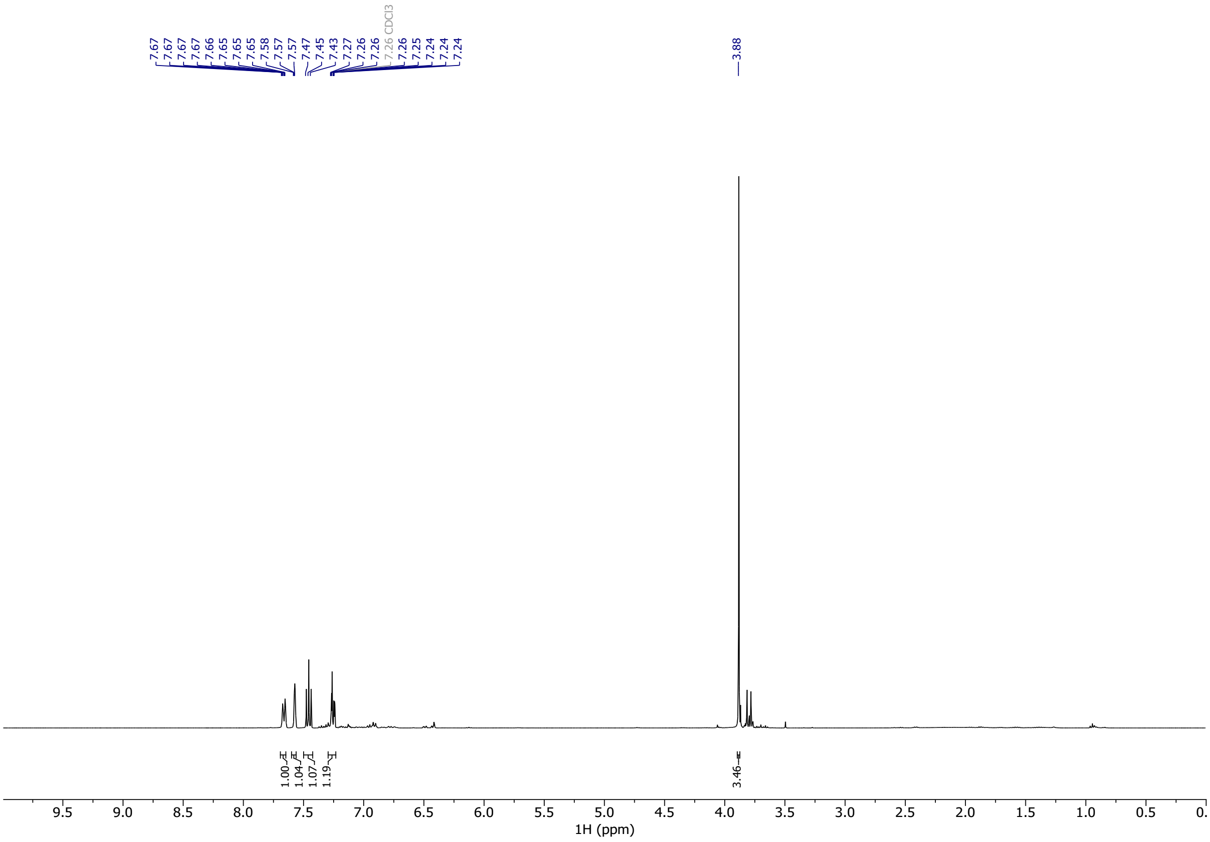


#### [13]C NMR Spectra of Compound S6

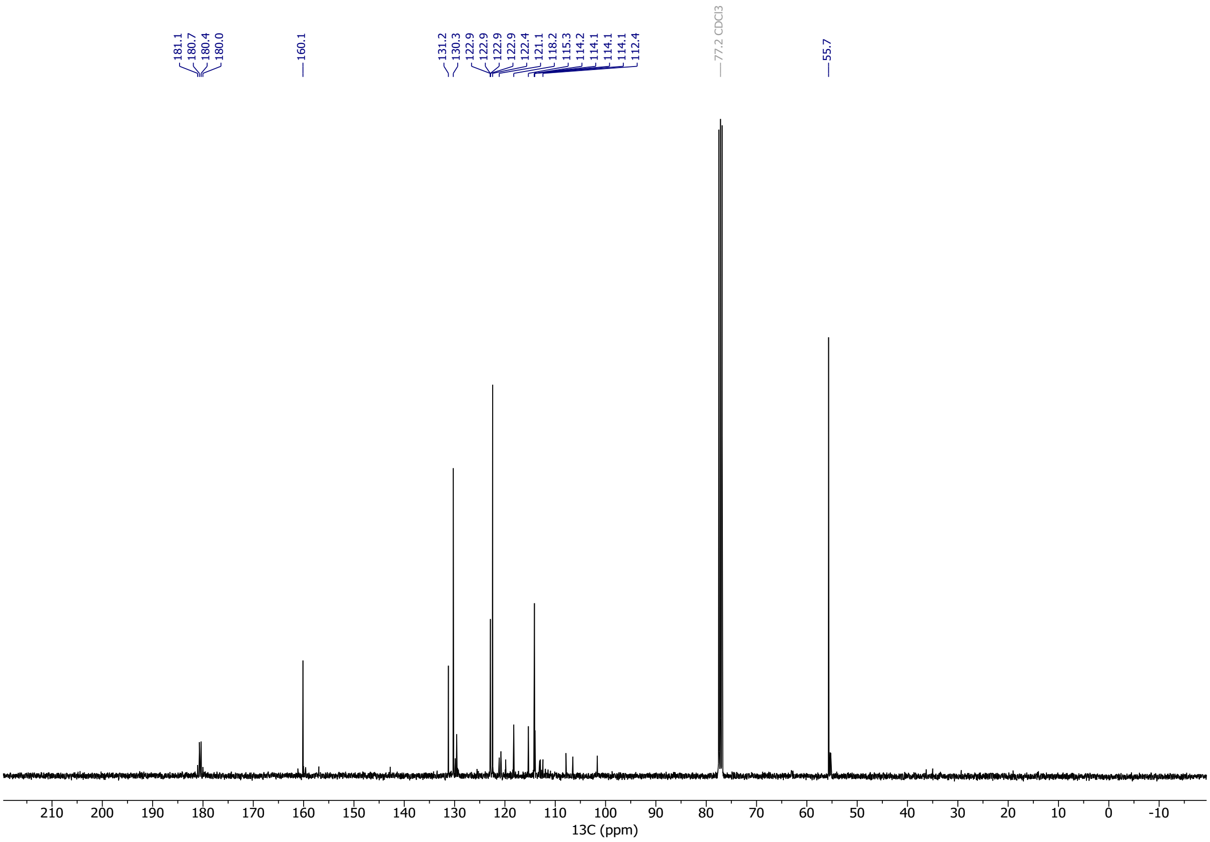

### $^{1}H$ NMR Spectra of Compound S7

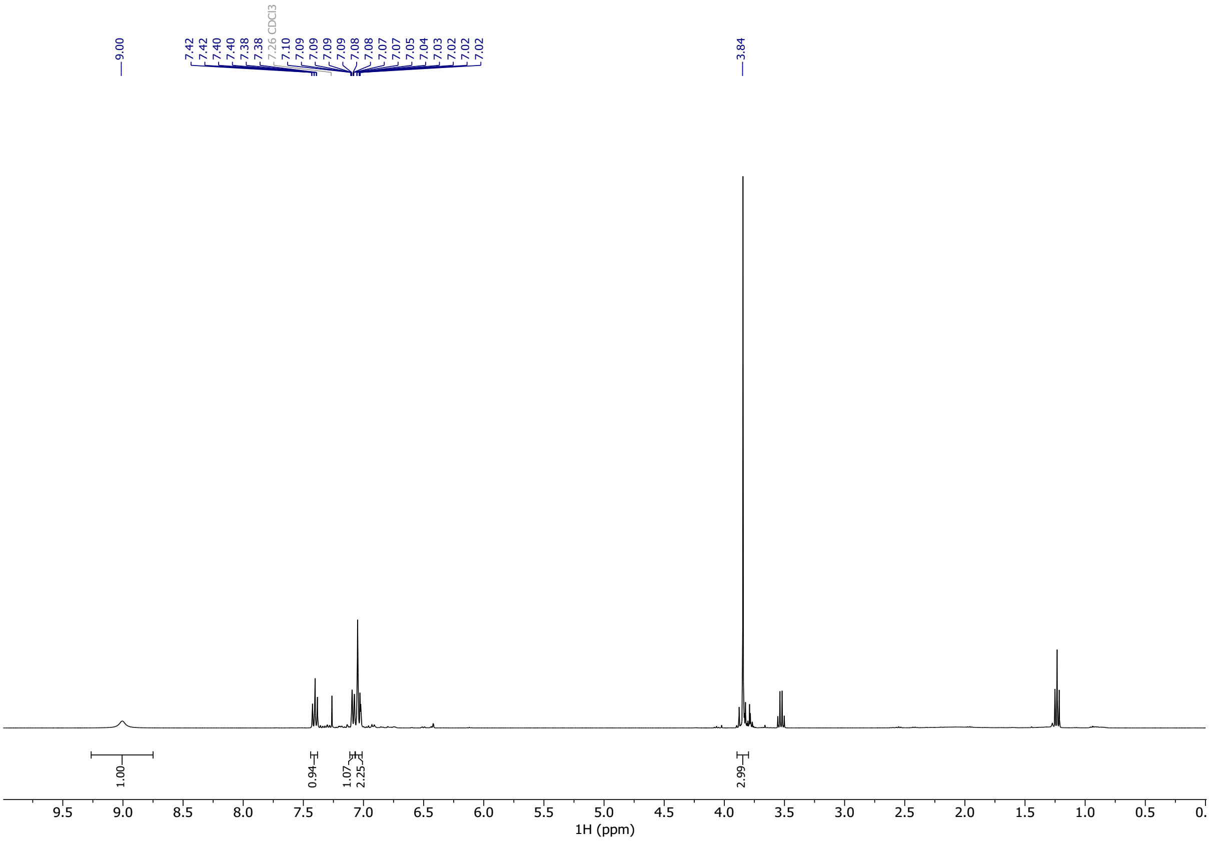


### $^{13}C$ NMR Spectra of Compound S7

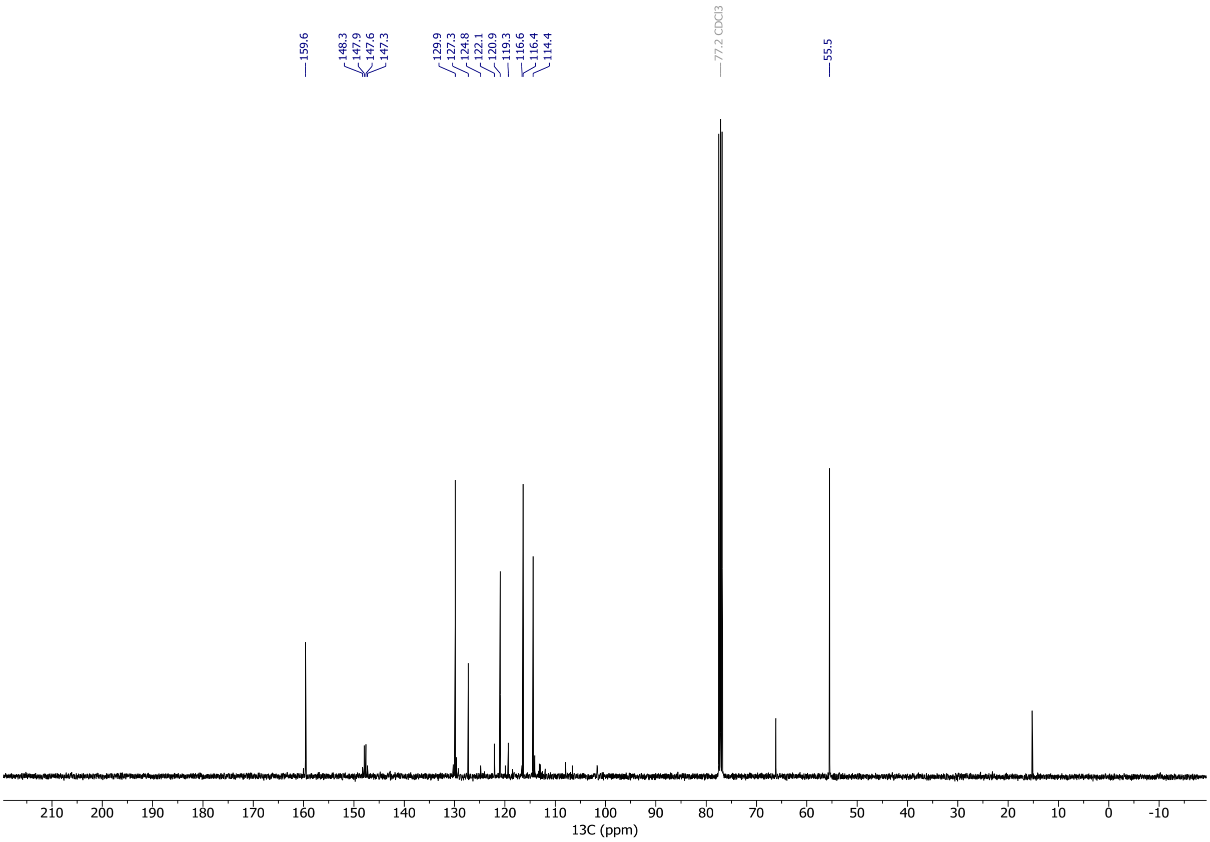

#### $^1$H NMR Spectra of Compound S8

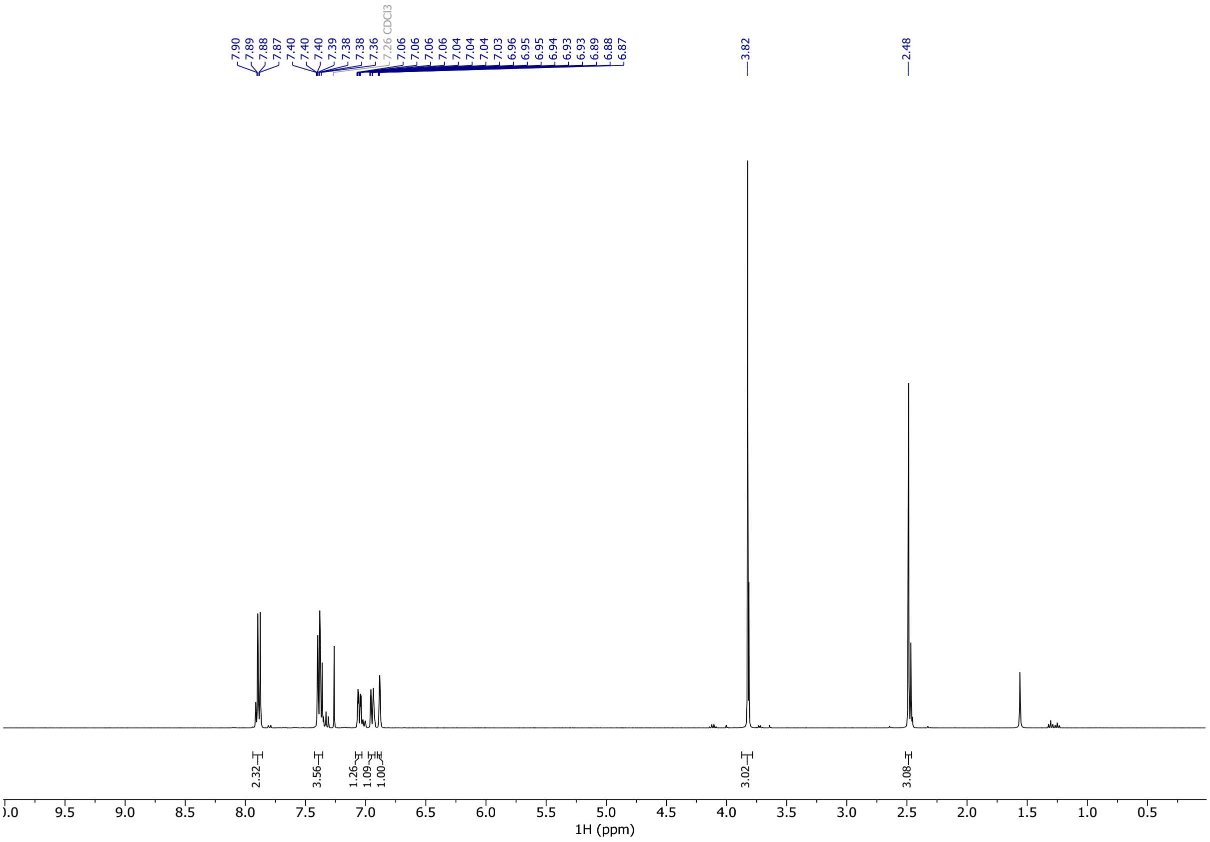

#### $^{13}$C NMR Spectra of Compound S8

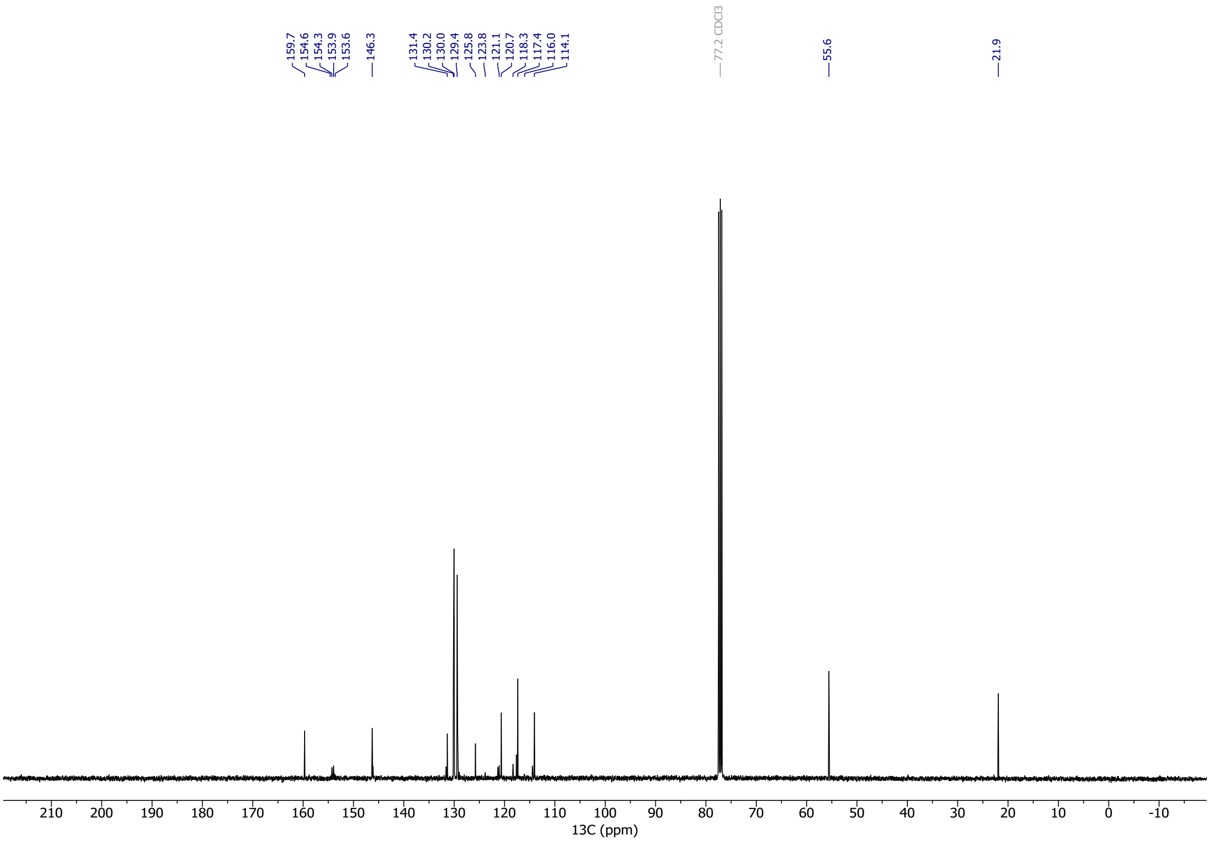

### $^1$H NMR Spectra of Compound S9

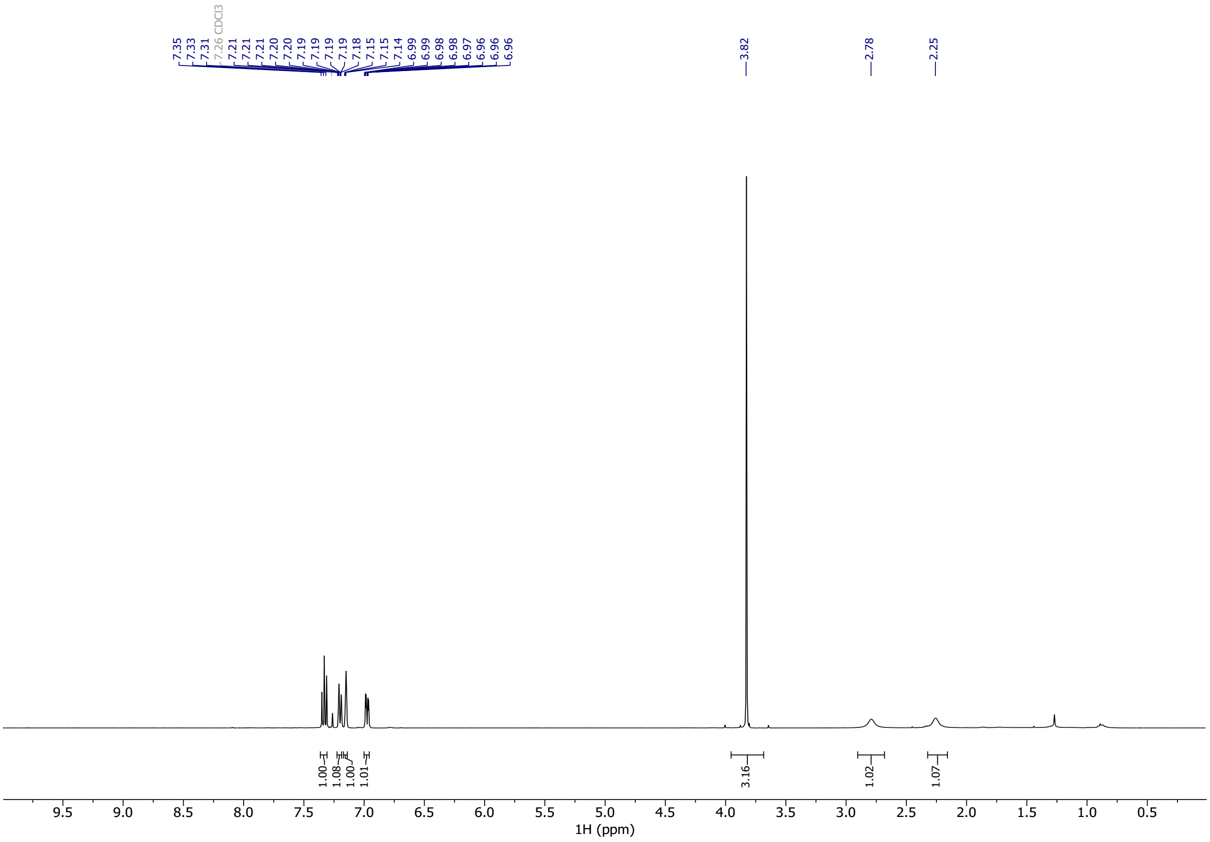

### $^{13}$C NMR Spectra of Compound S9

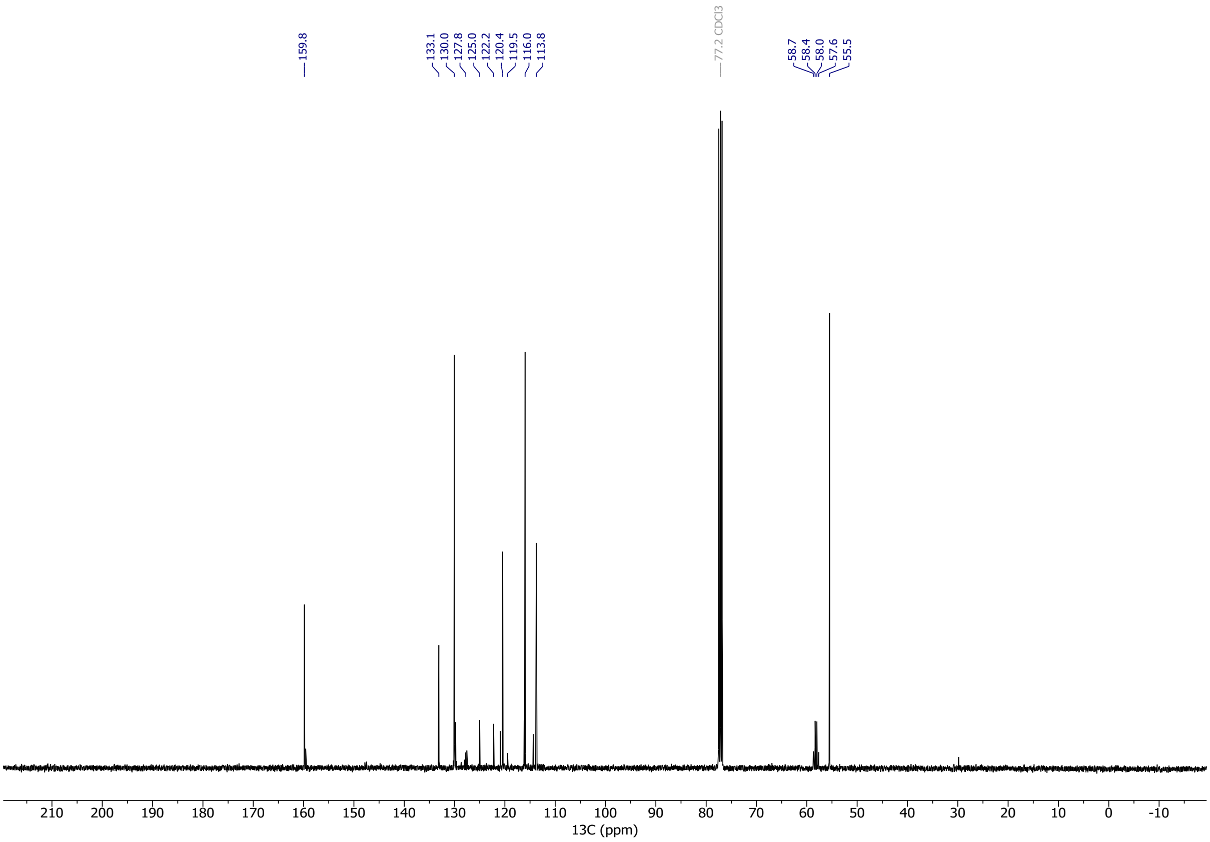

#### $^{1}$H NMR Spectra of Compound S13

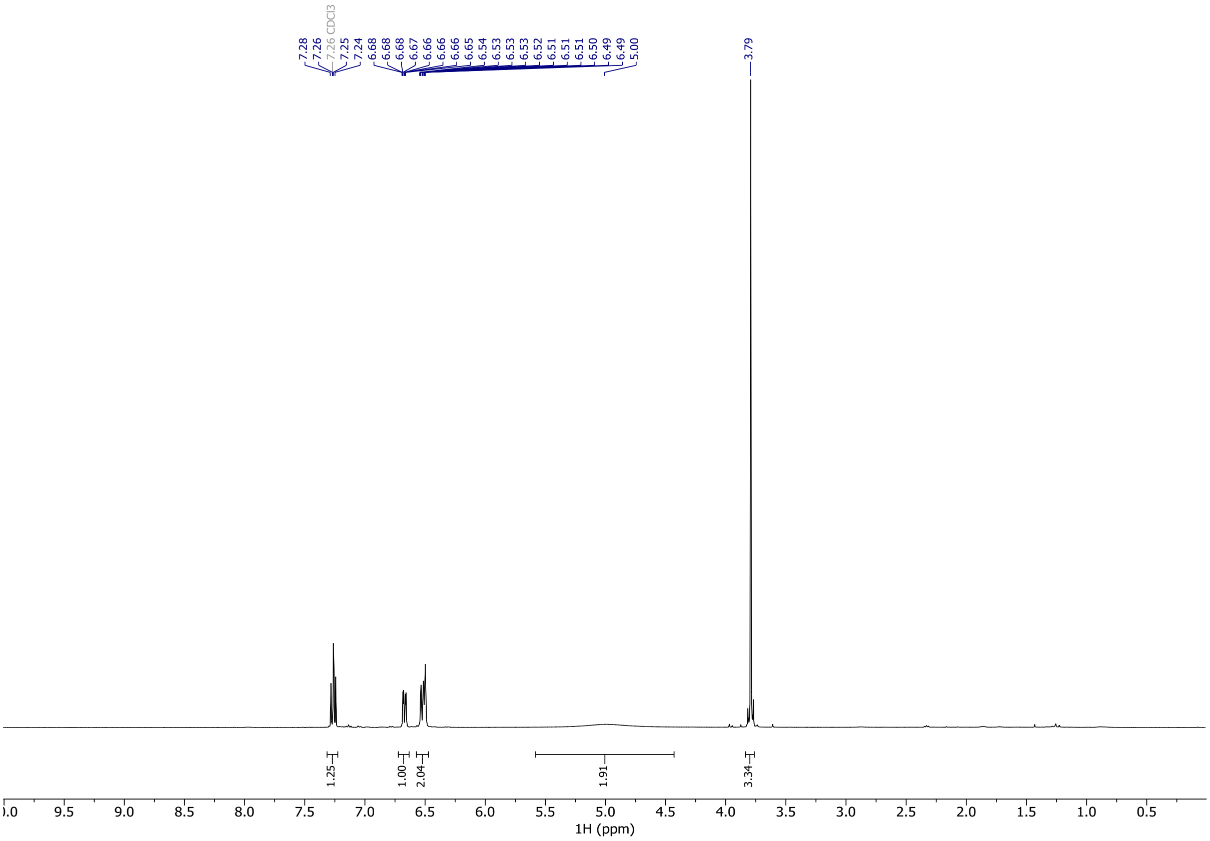


#### $^{13}$C NMR Spectra of Compound S13

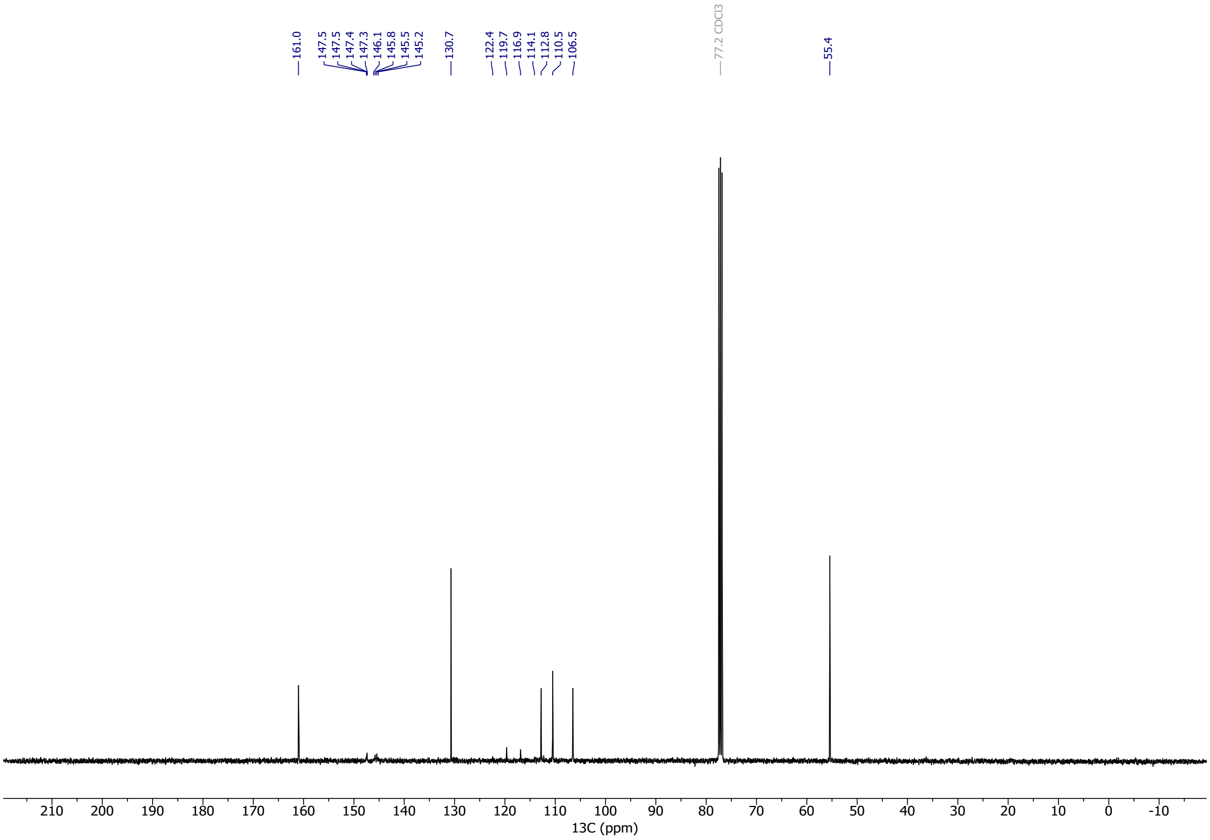

#### [1]H NMR Spectra of Compound S14

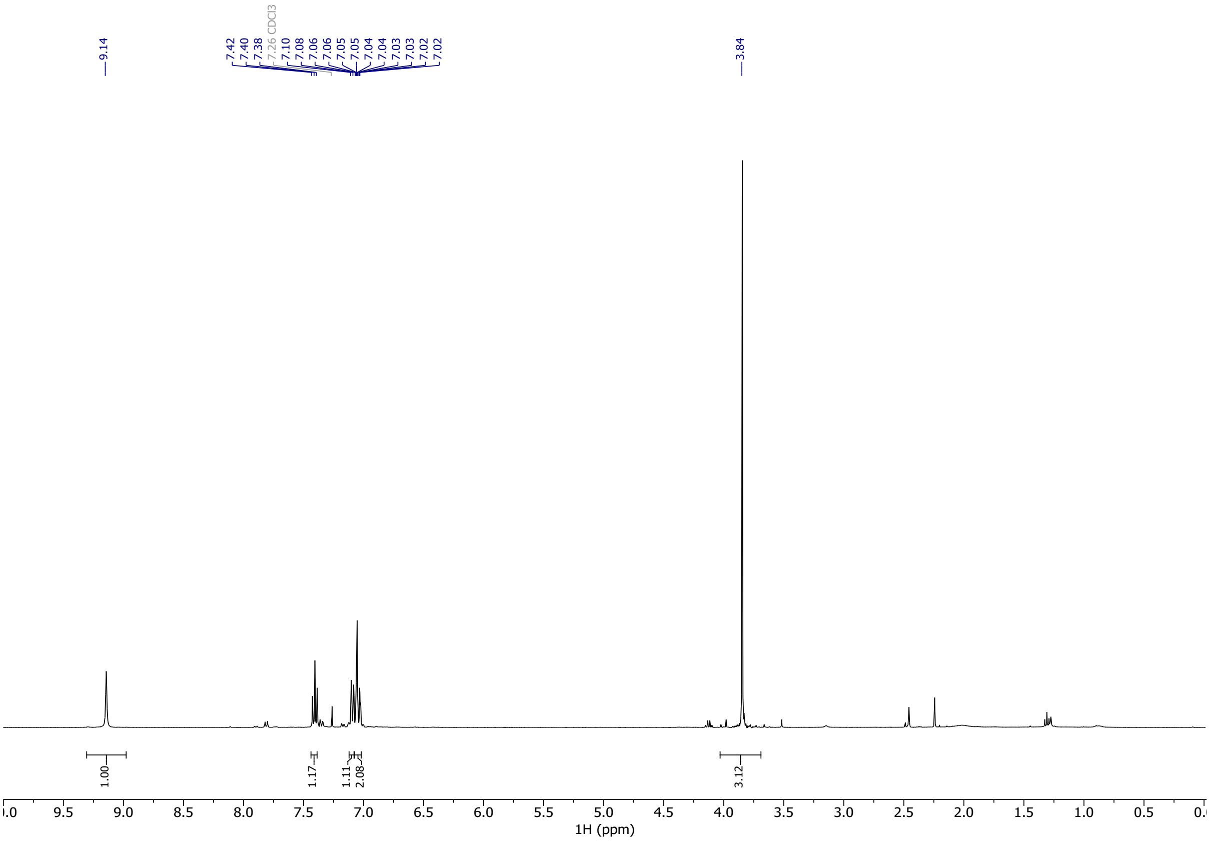


#### [13]C NMR Spectra of Compound S14

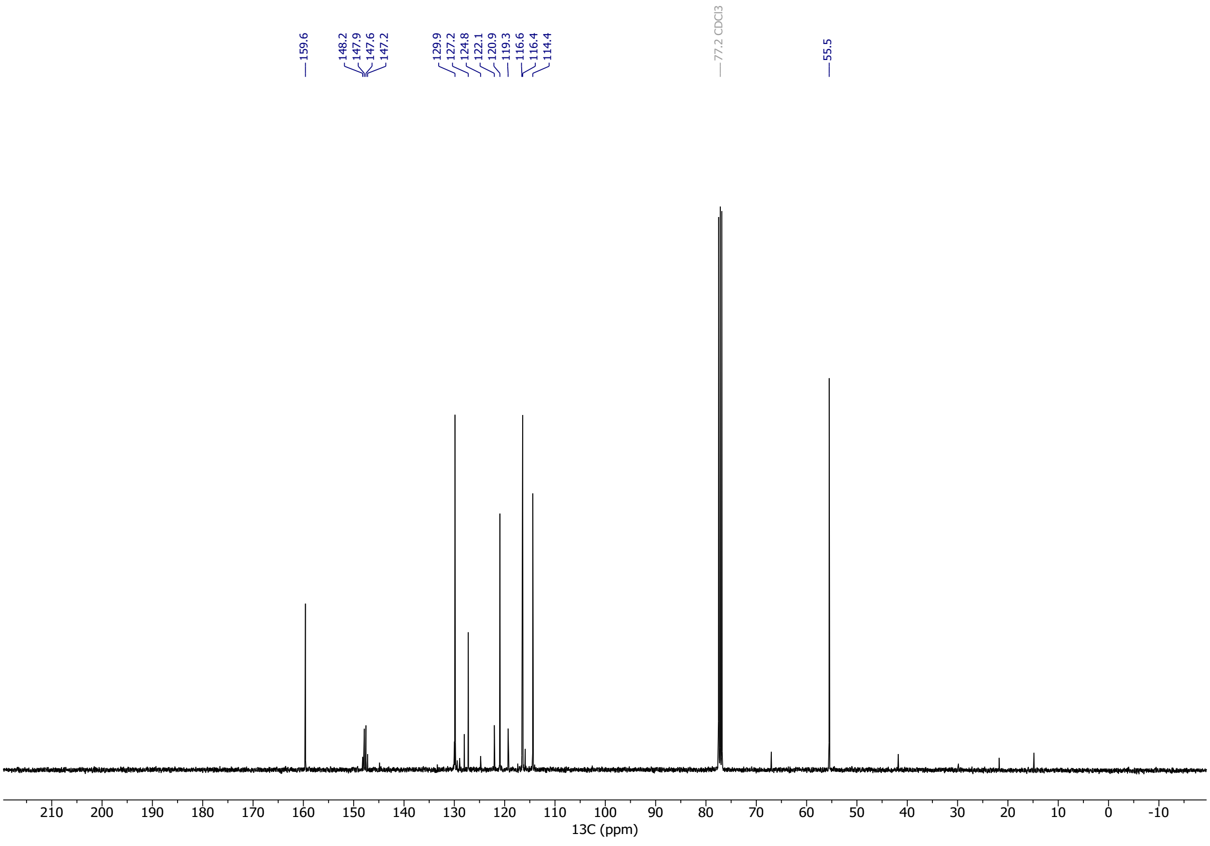

#### [1]H NMR Spectra of Compound S10

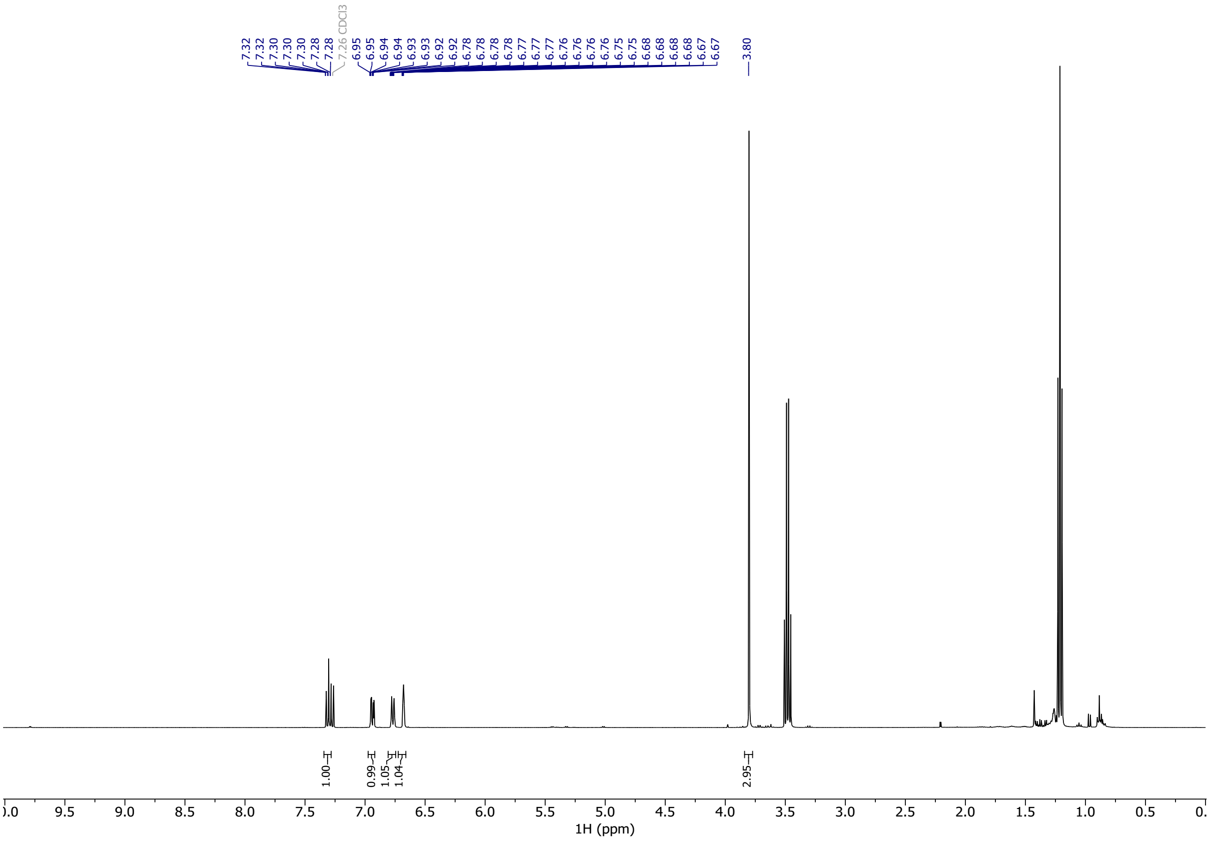


#### [13]C NMR Spectra of Compound S10

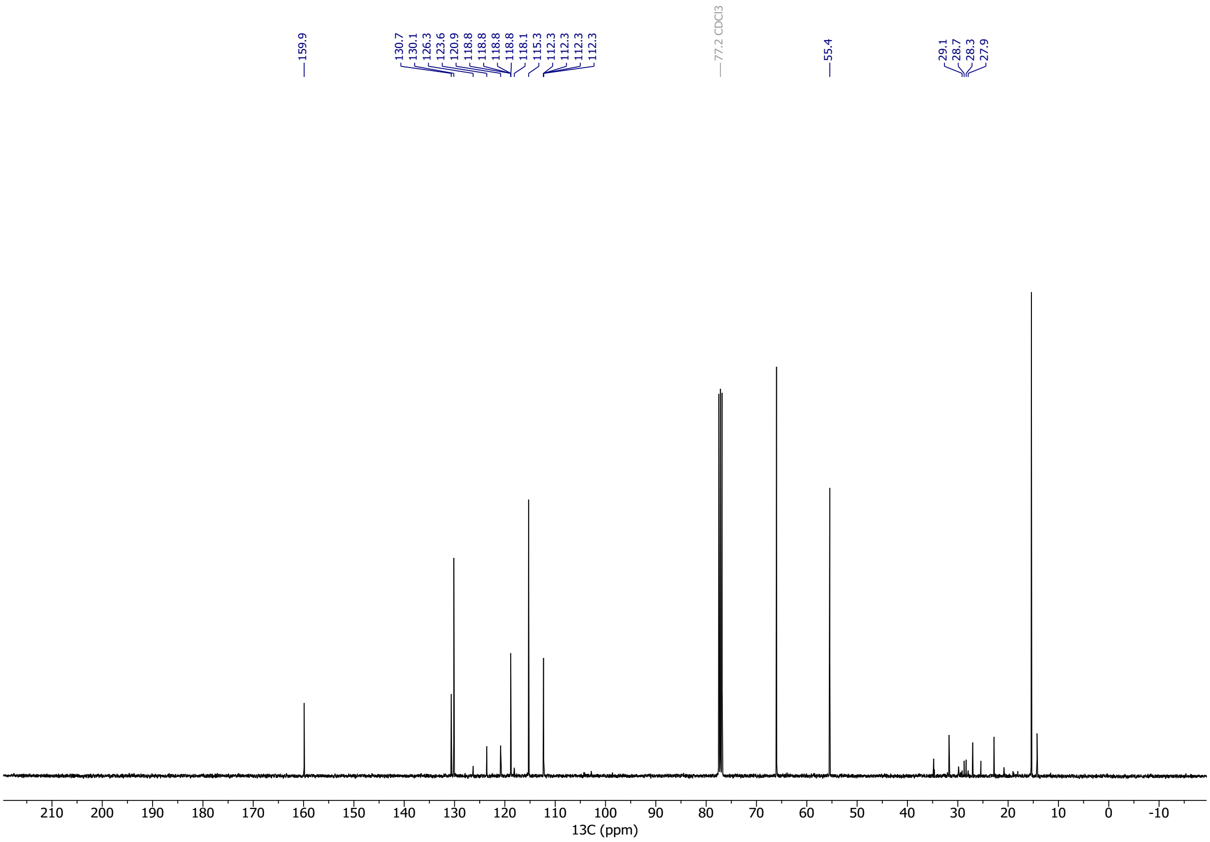

#### $^{1}$H NMR Spectra of Compound S12

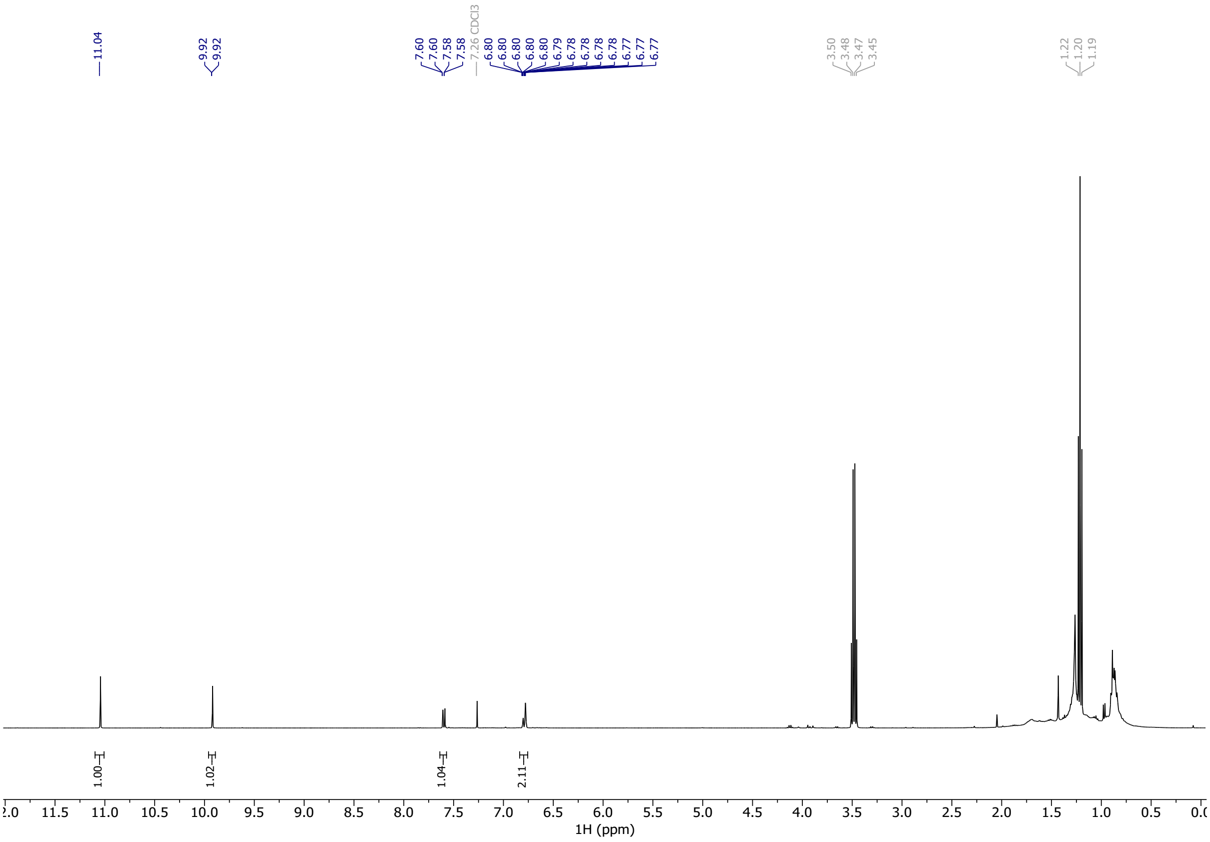


#### $^{13}$C NMR Spectra of Compound S12

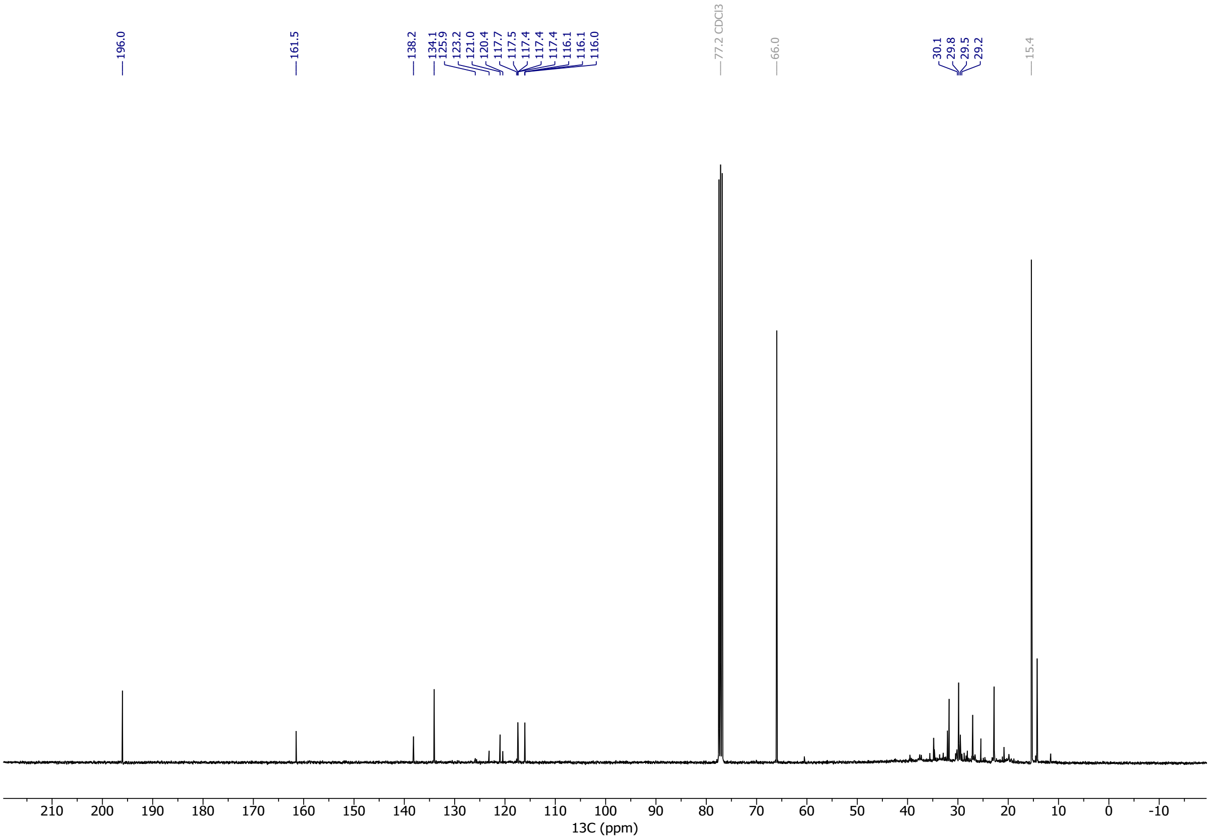

#### [1]H NMR Spectra of Compound S15

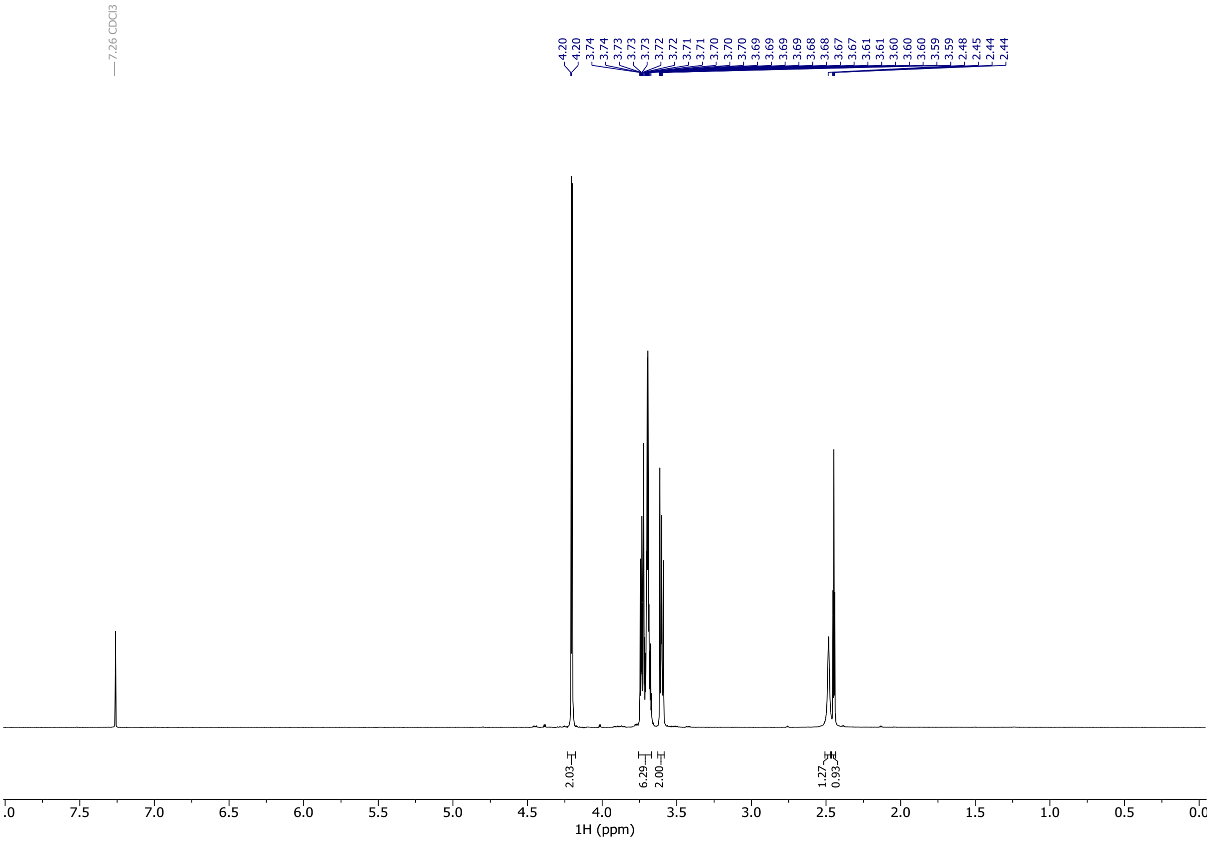


#### [13]C NMR Spectra of Compound S15

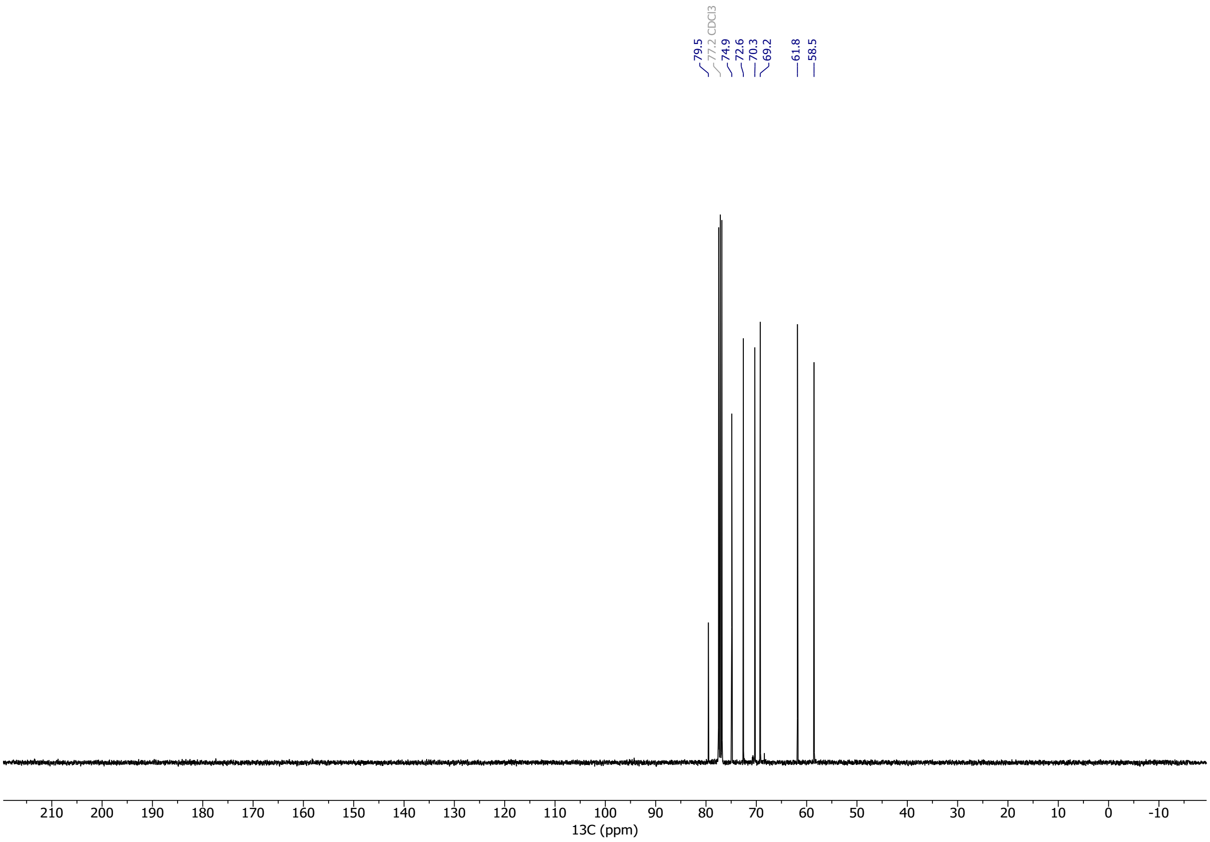

#### [1]H NMR Spectra of Compound S16

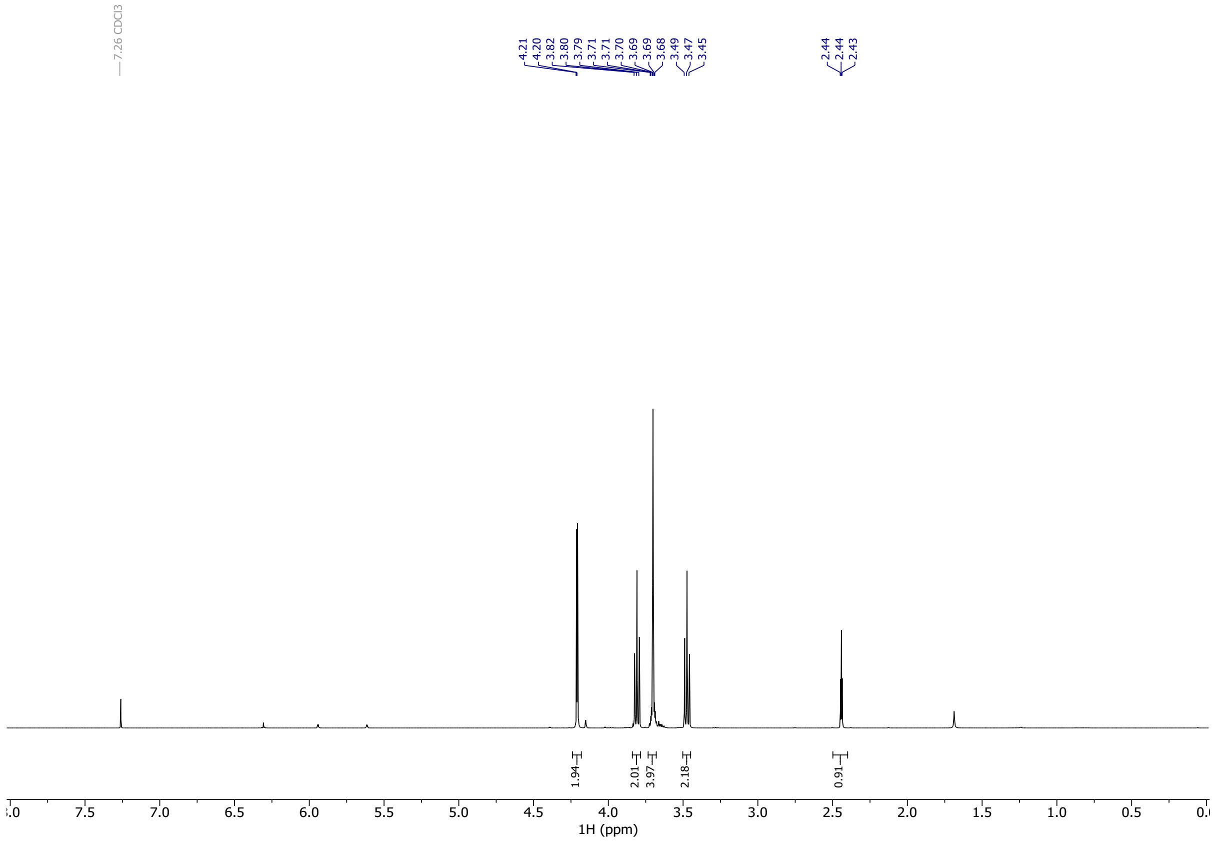


#### [13]C NMR Spectra of Compound S16

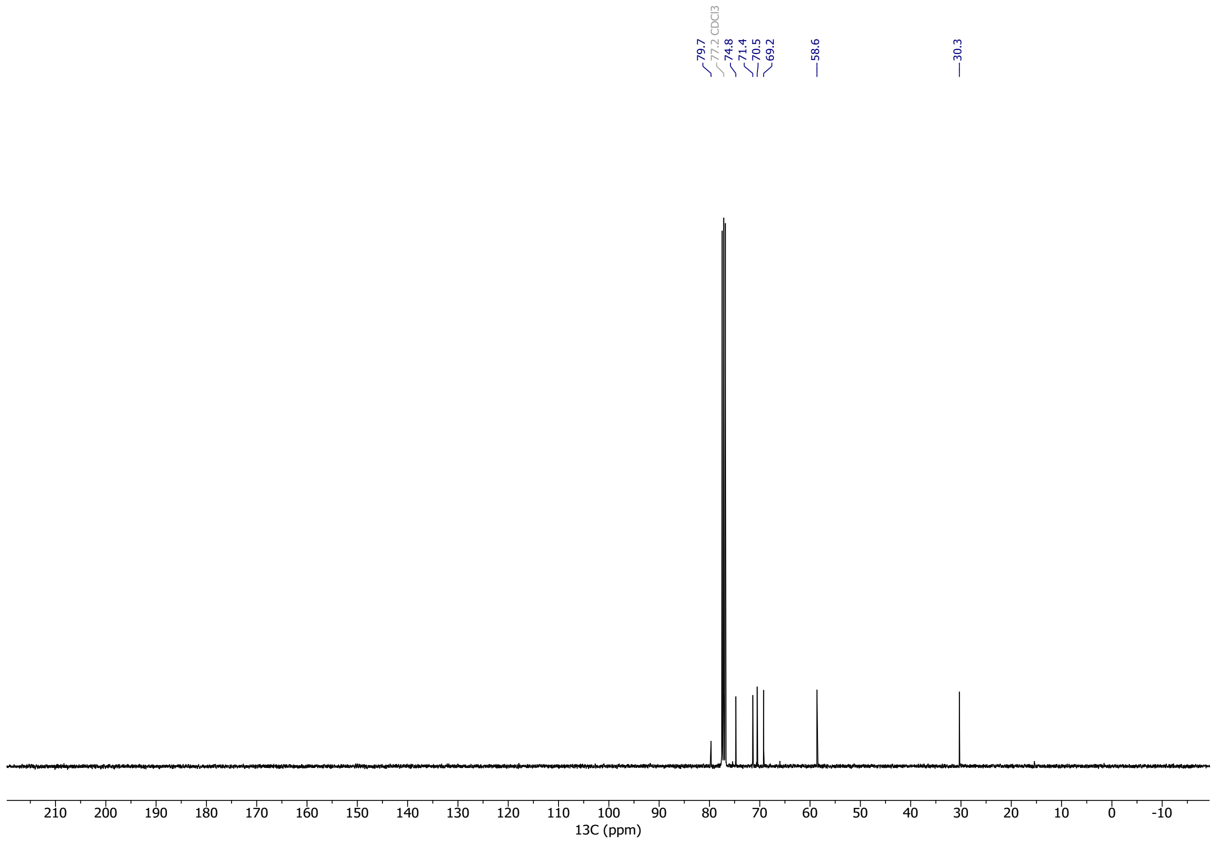

### [1]H NMR Spectra of Compound S17

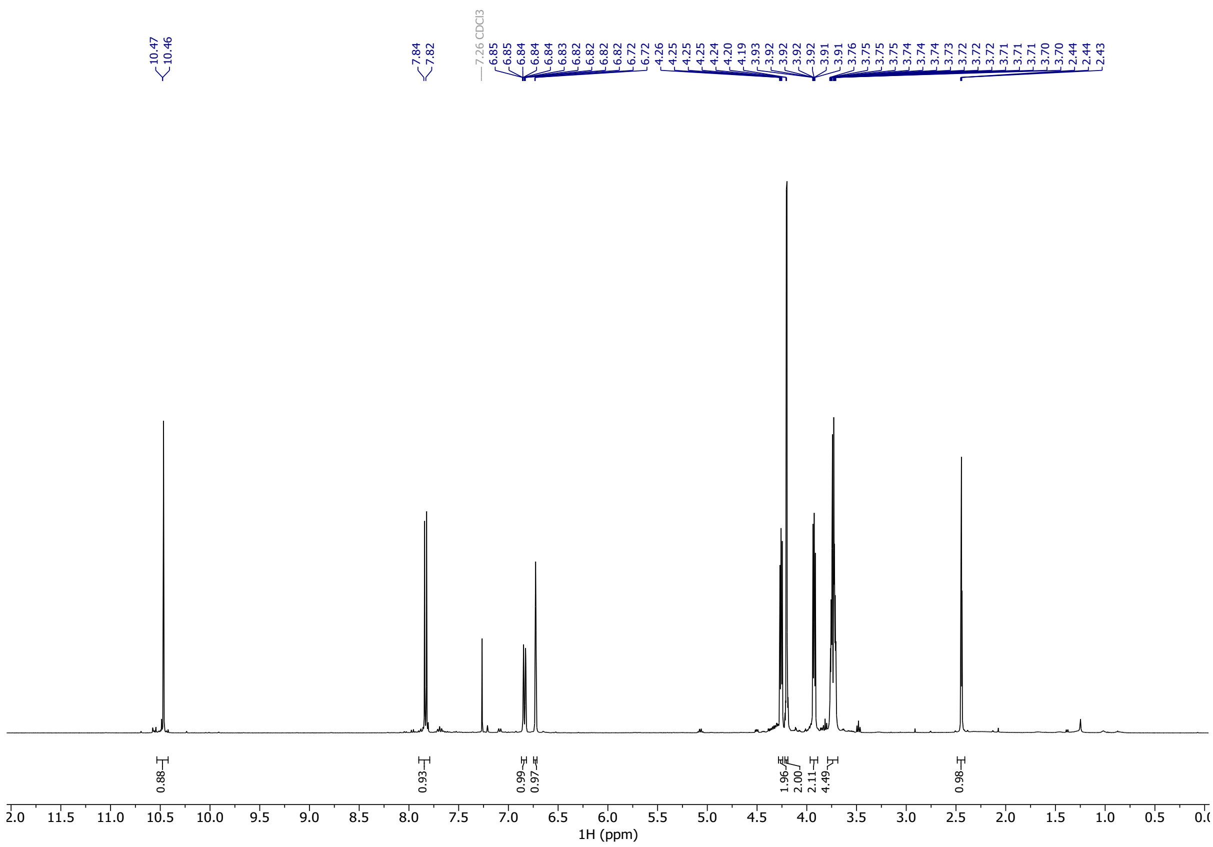

### [13]C NMR Spectra of Compound S17

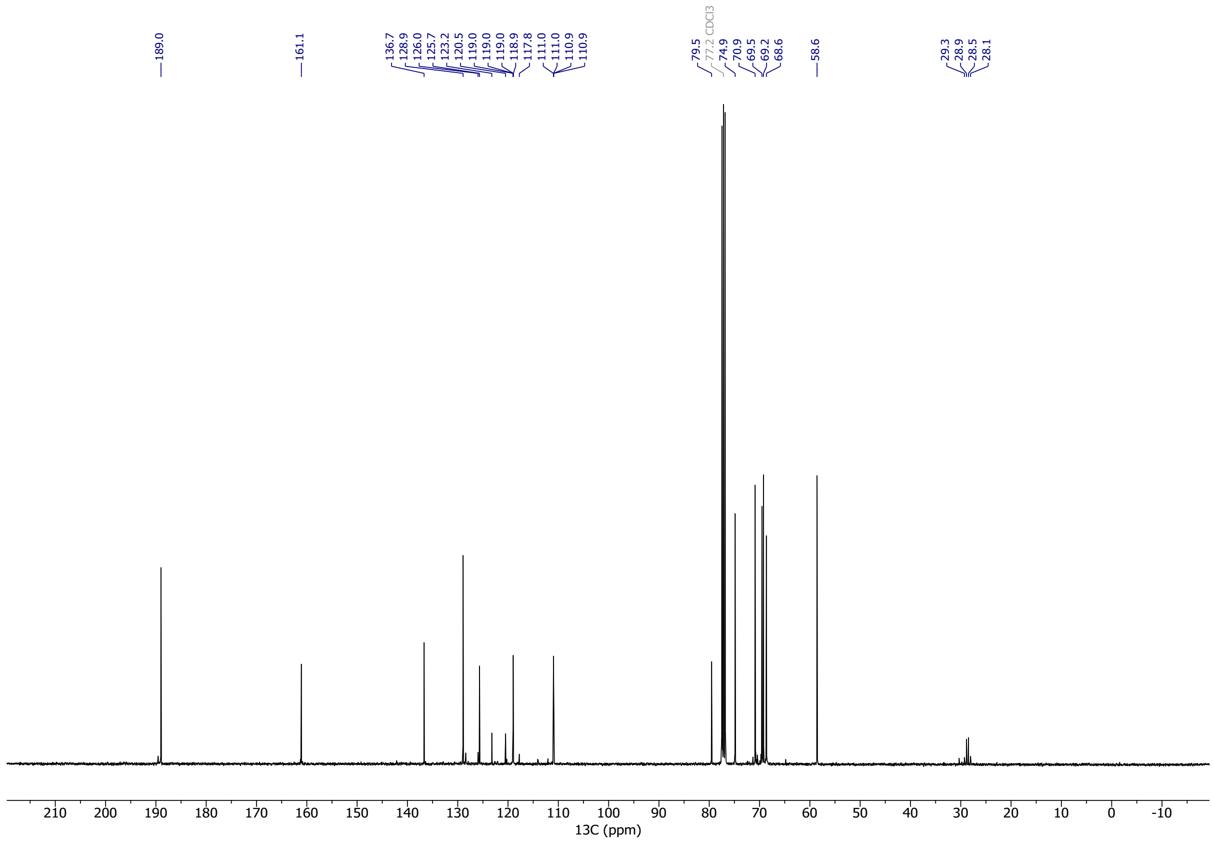

#### $^{1}$H NMR Spectra of Compound 10

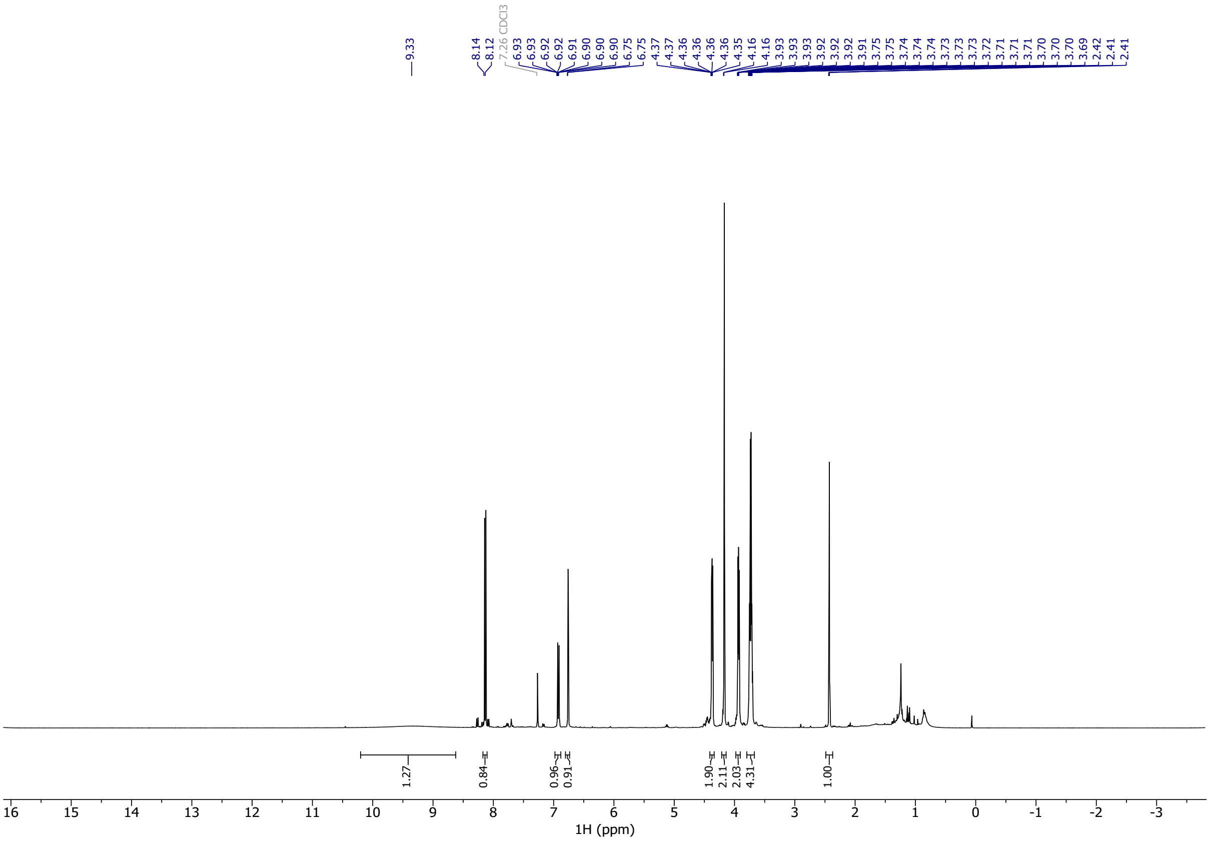


#### $^{13}$C NMR Spectra of Compound 10

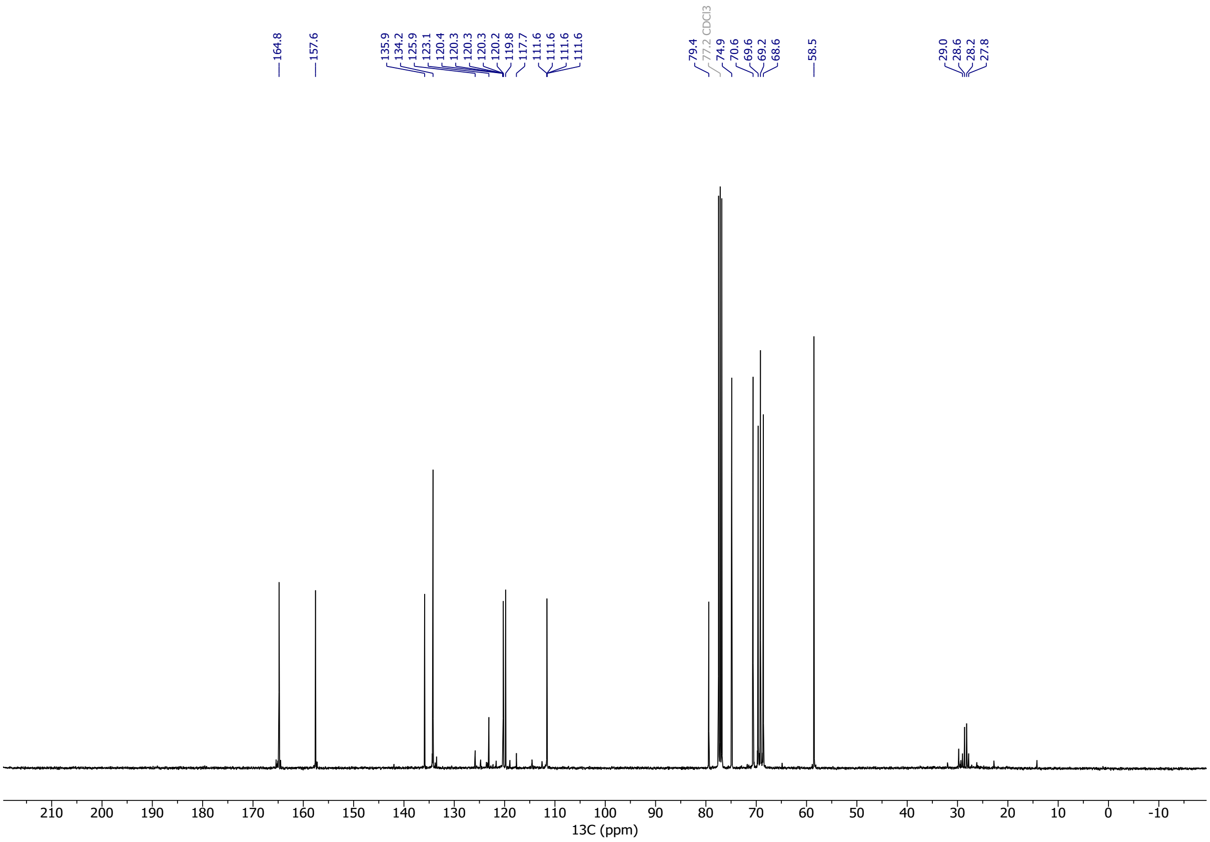

#### $^{1}$H NMR Spectra of Compound S18

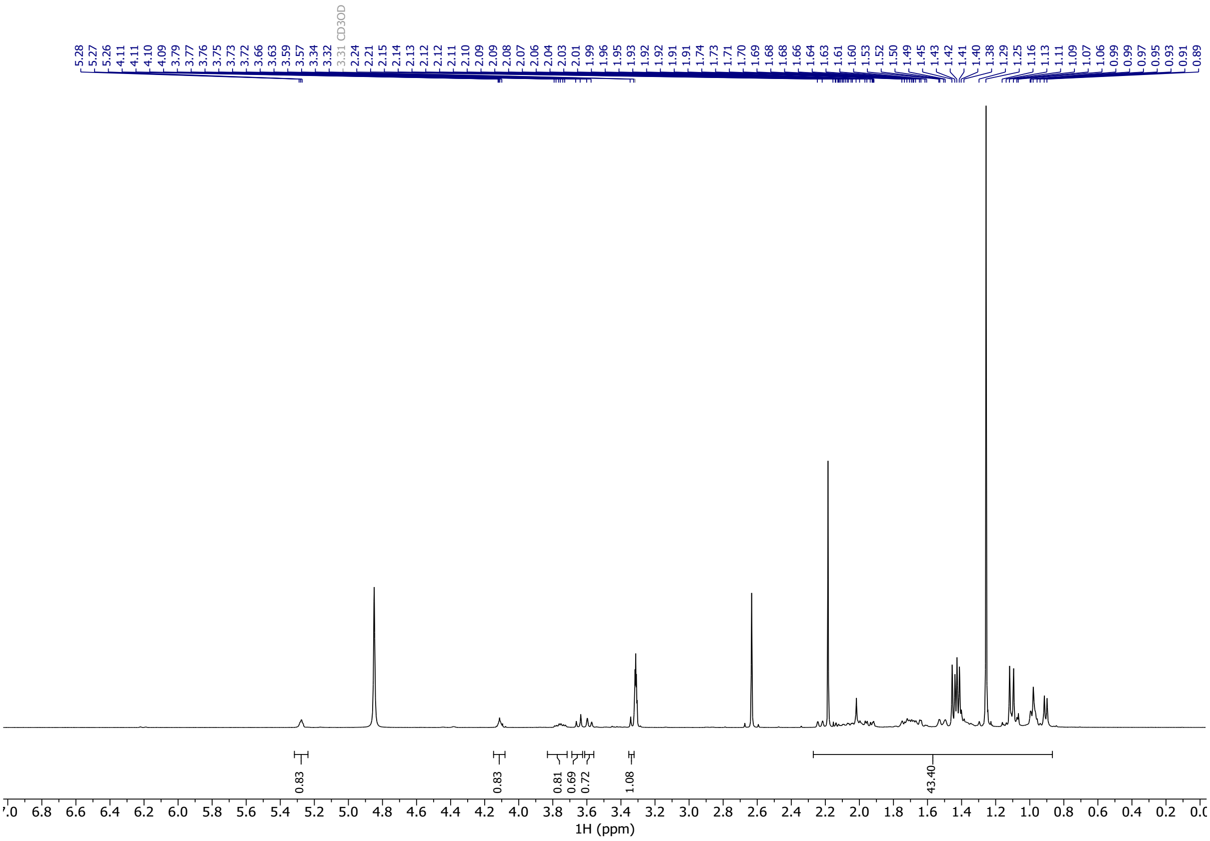


#### $^{13}$C NMR Spectra of Compound S18

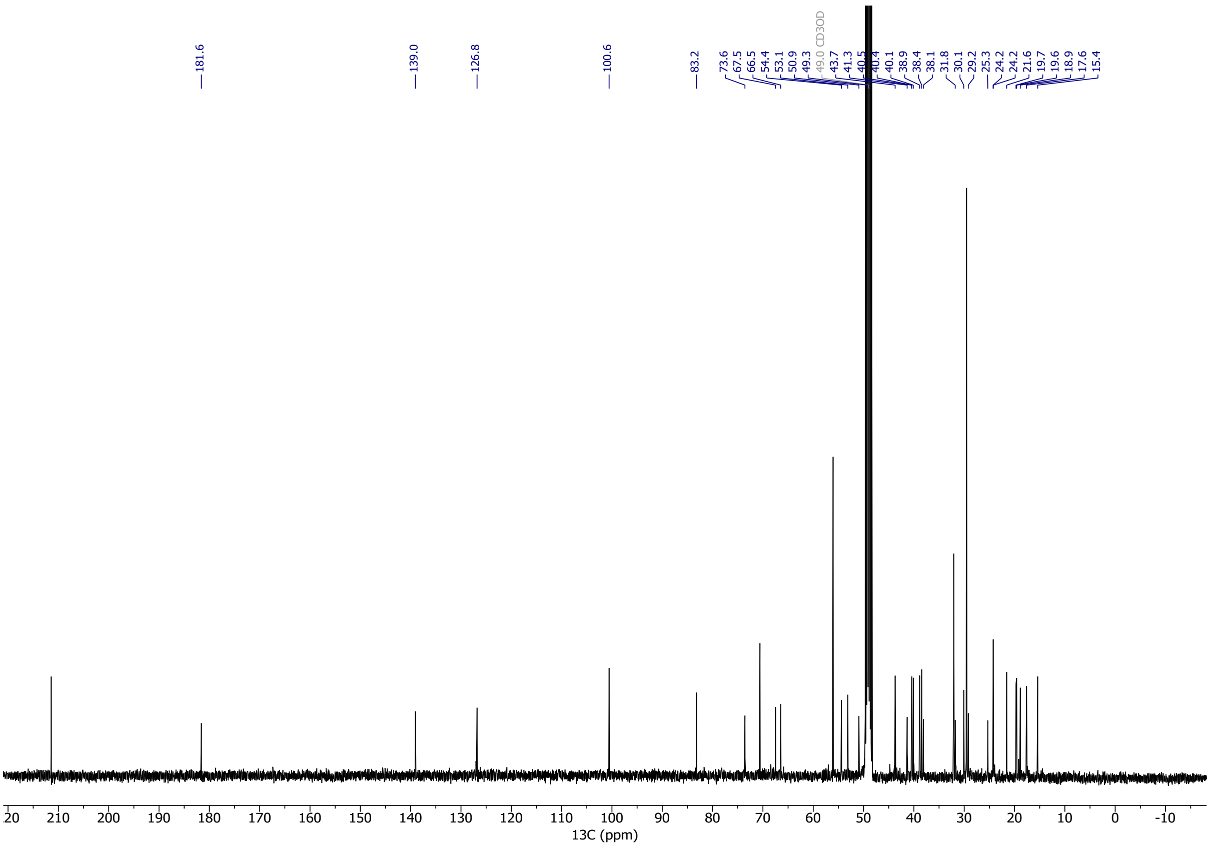

#### $^{1}H$ NMR Spectra of Compound 11

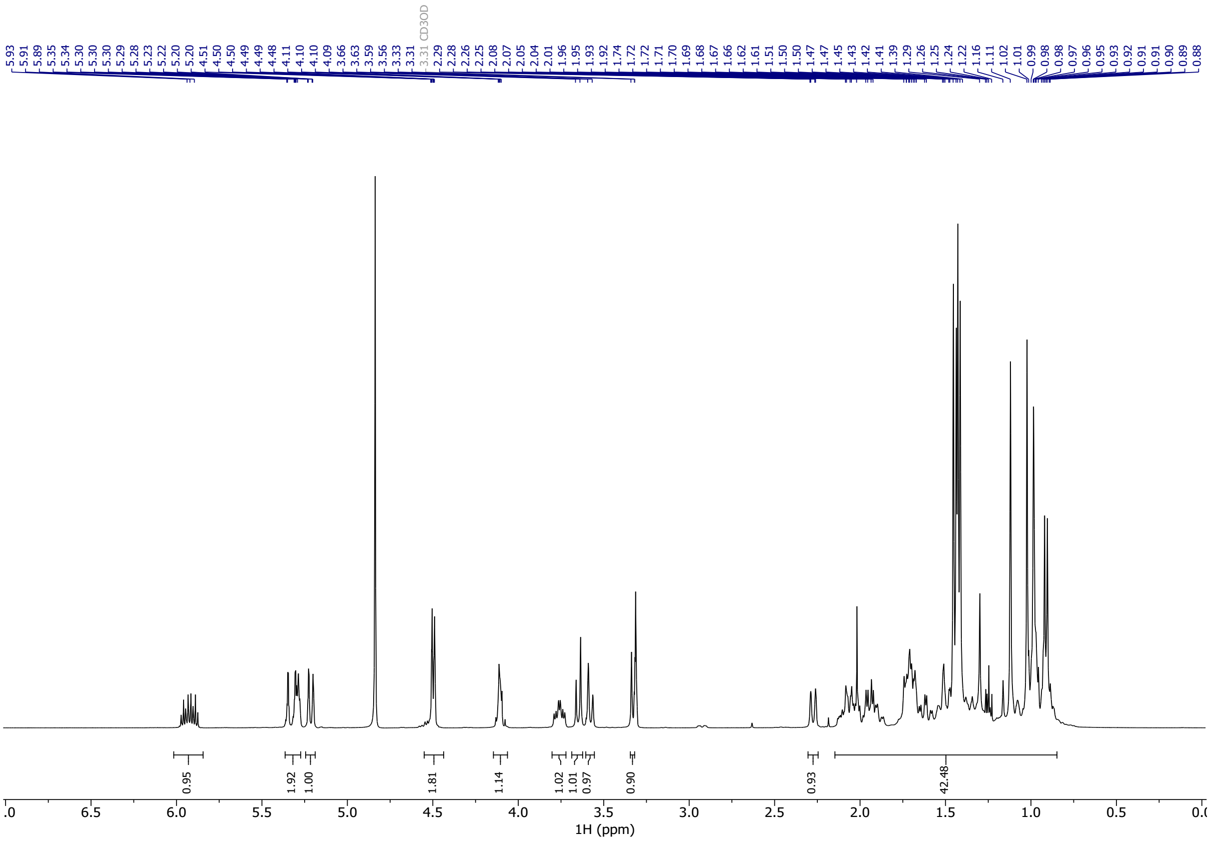


#### $^{13}C$ NMR Spectra of Compound 11

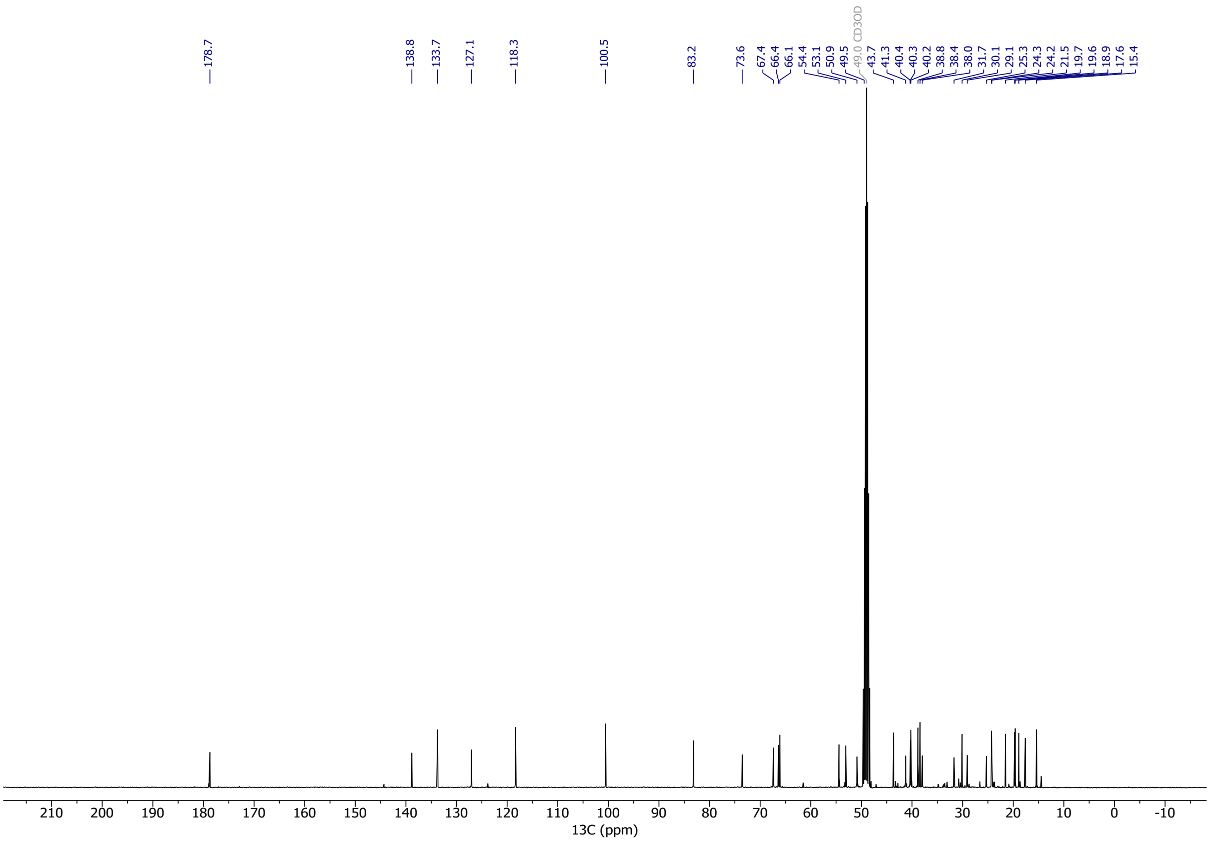

### $^{1}$H NMR Spectra of Compound 13

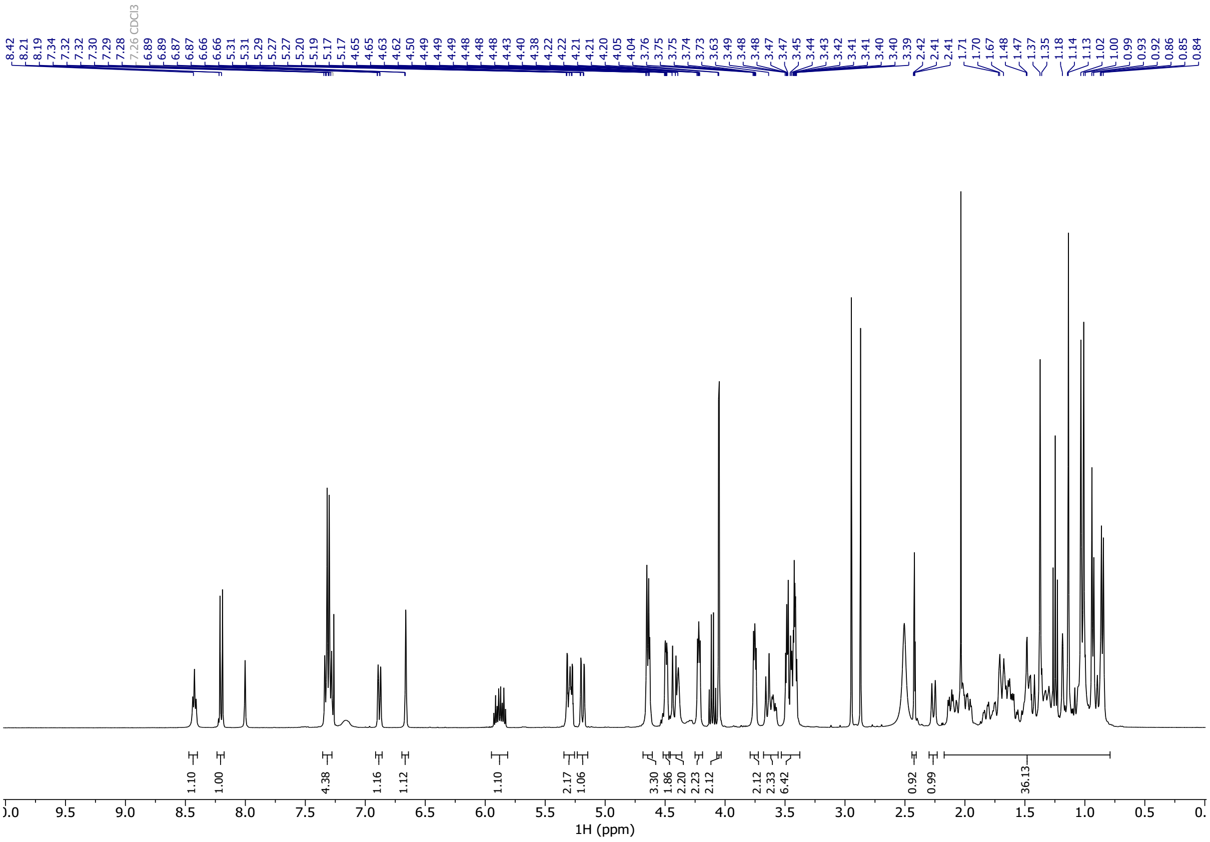


### $^{13}$C NMR Spectra of Compound 13

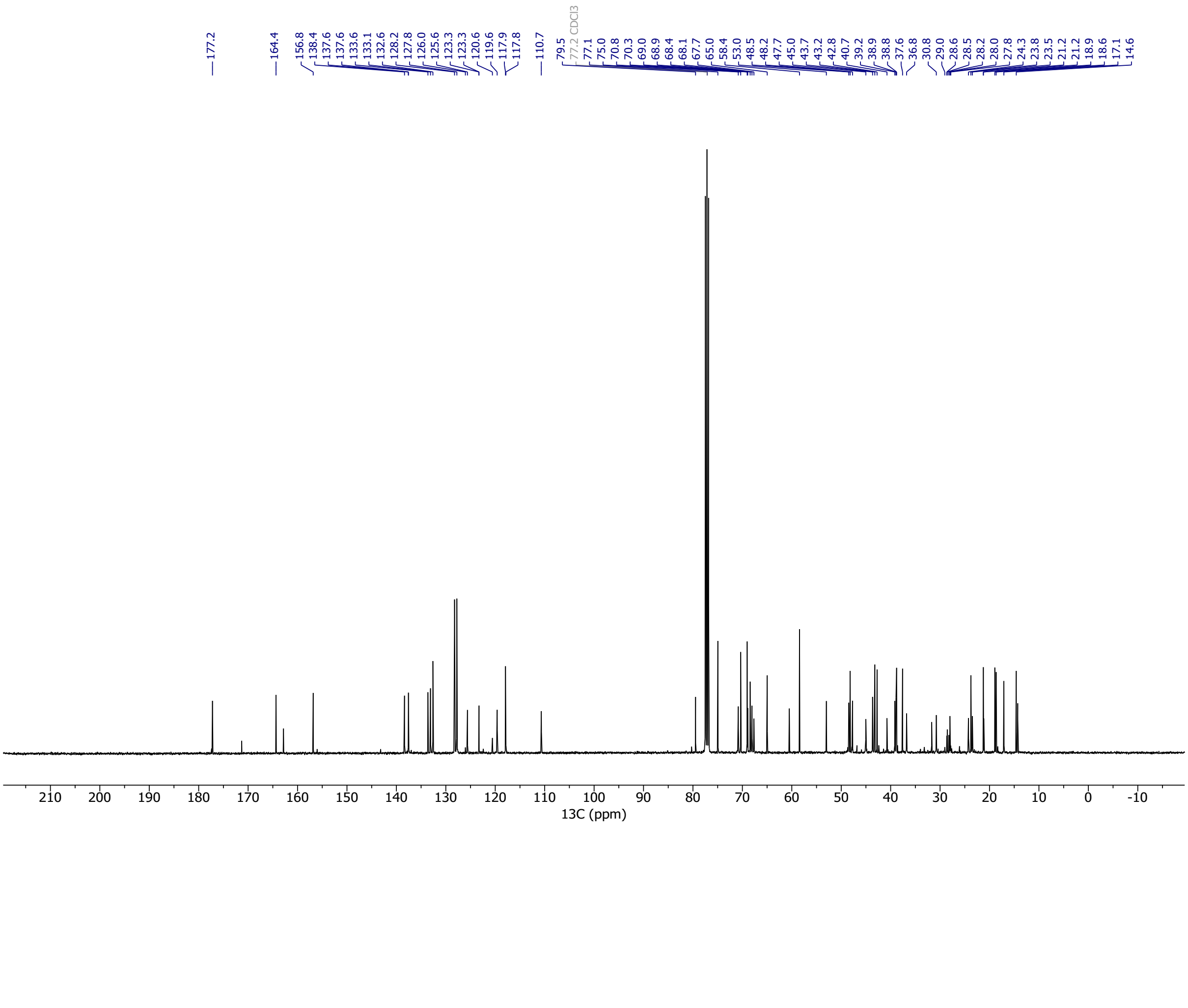

### [1]H NMR Spectra of Compound 14

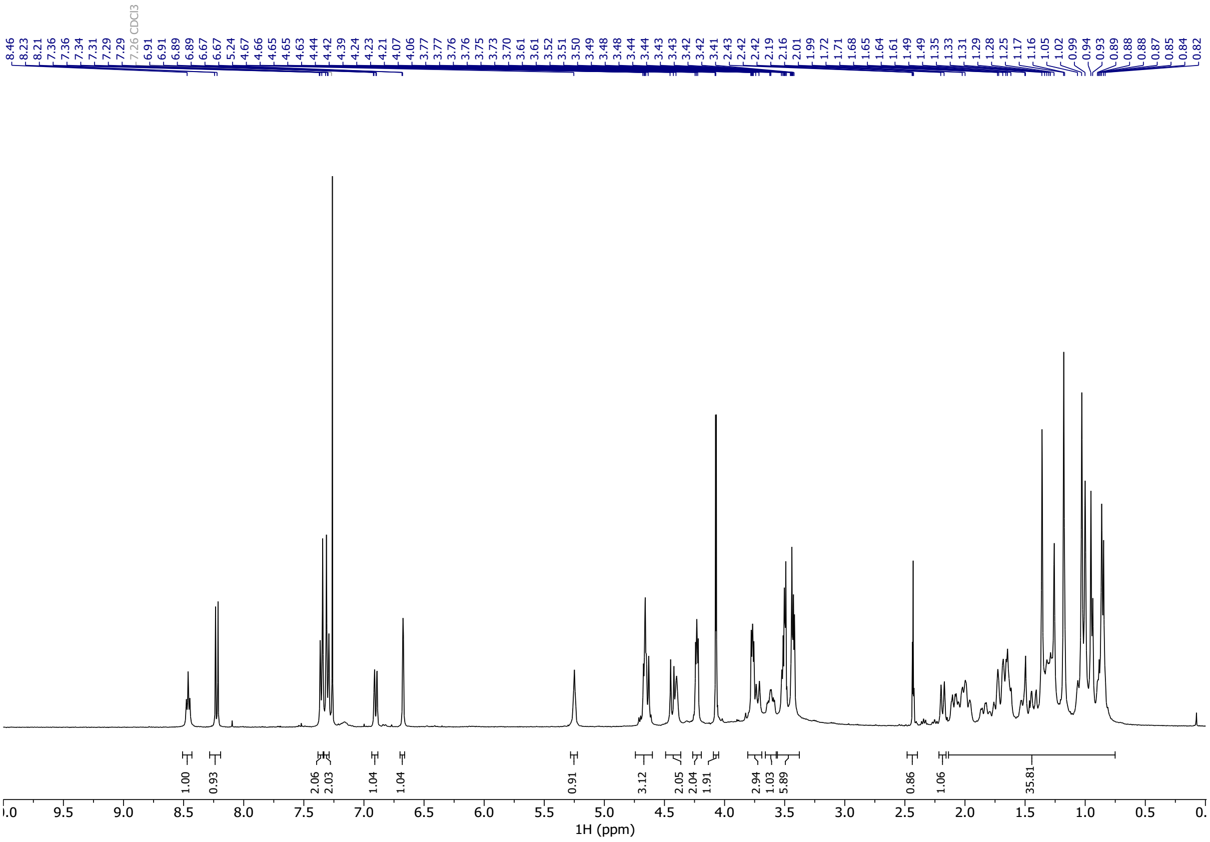

### [13]C NMR Spectra of Compound 14

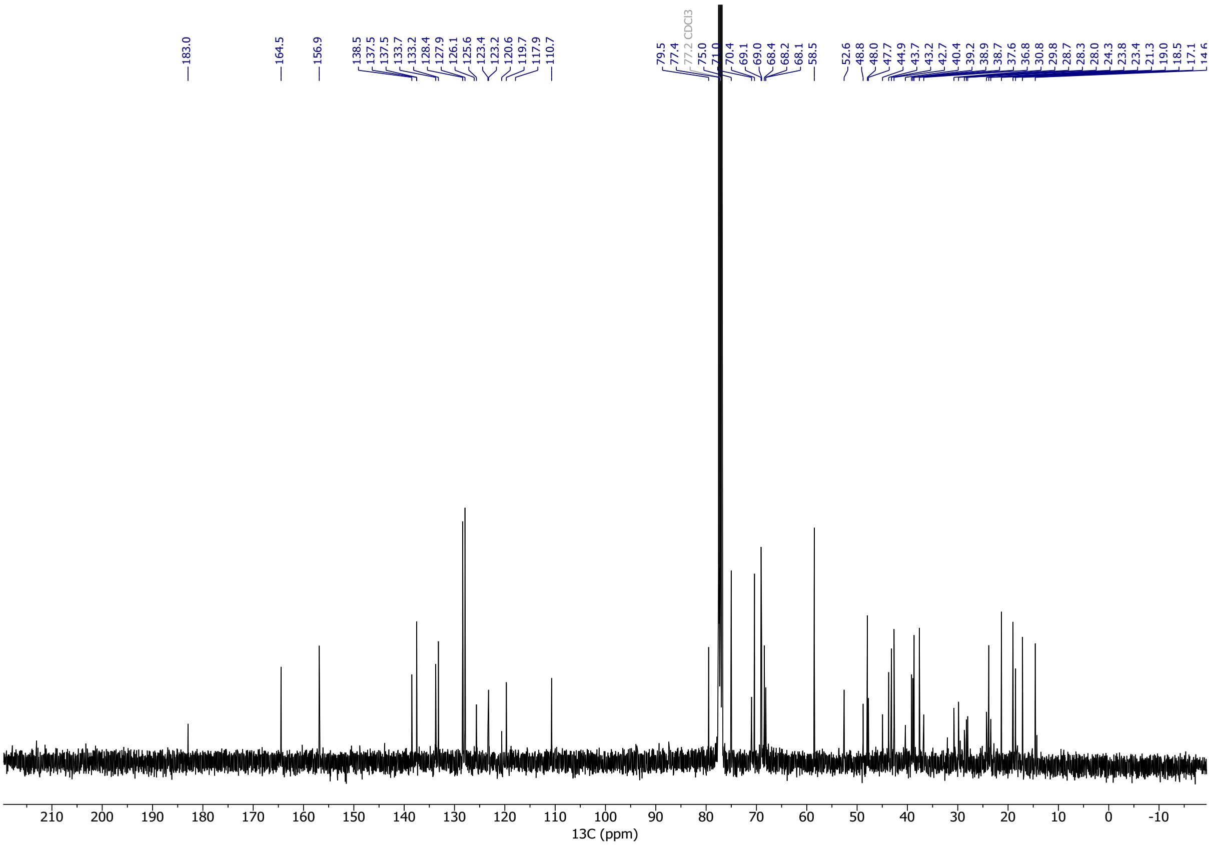

#### $^1$H NMR Spectra of Compound 15

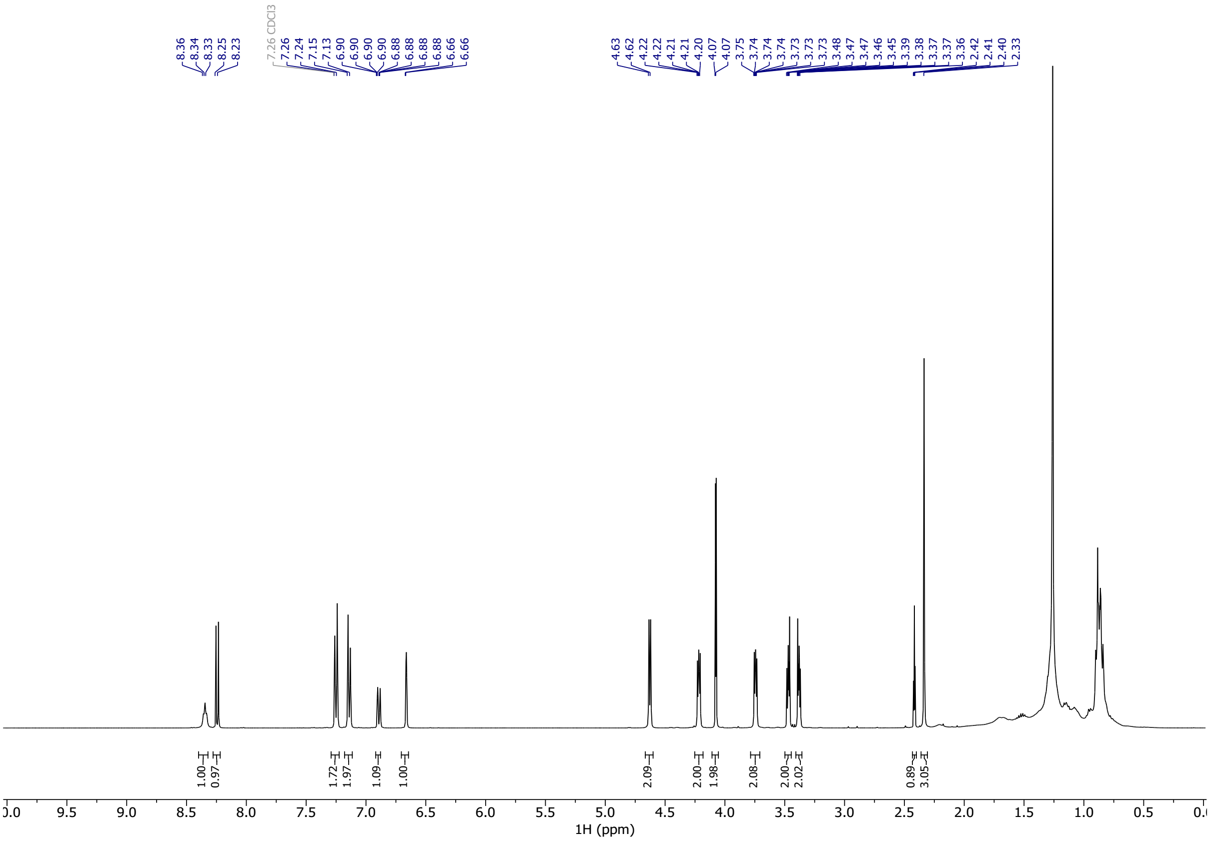


#### $^{13}$C NMR Spectra of Compound 15

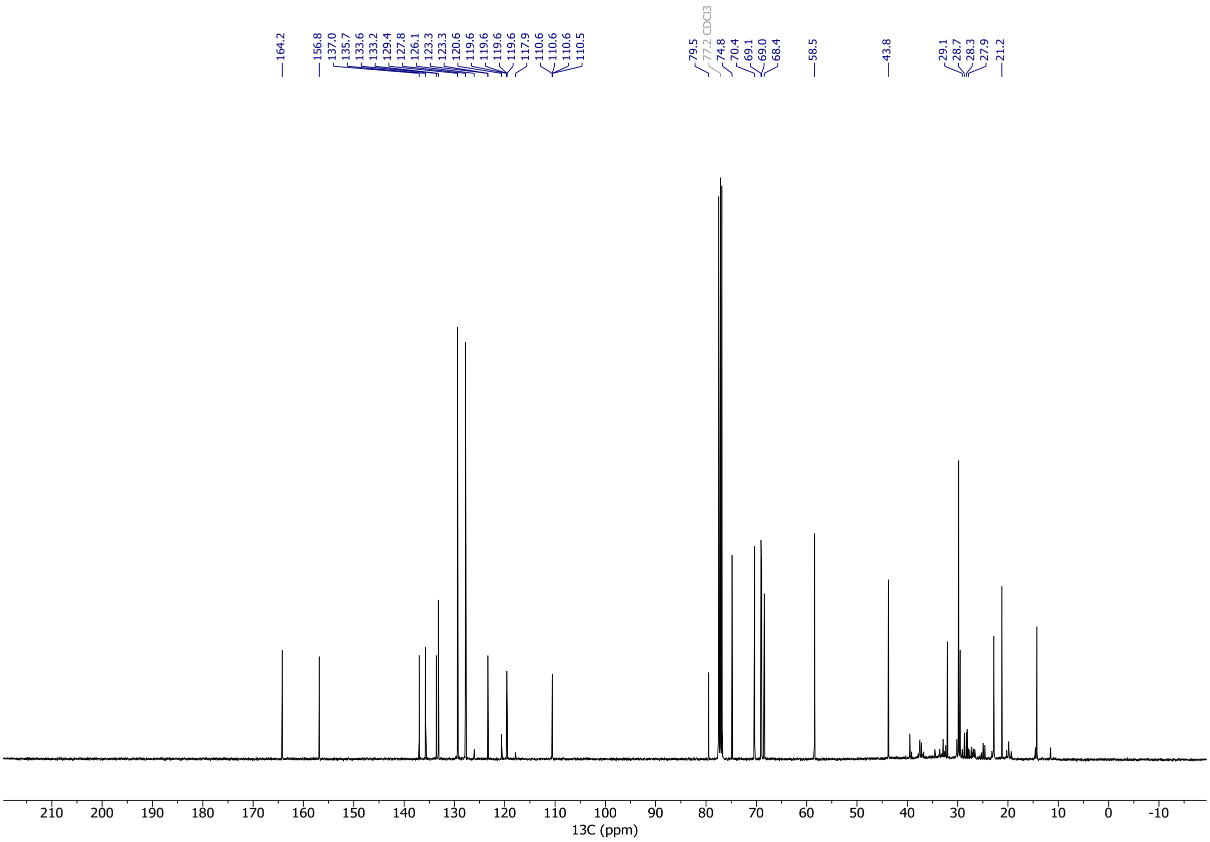
